# Chronic Electrophysiology Captures Progressive Variability in DNA-Damaged Cardiac Networks

**DOI:** 10.64898/2026.10.10.757663

**Authors:** Mabel Bartlett, Sara Gibson, Mats Forsell, Aaditya Nandakumar, Jacqueline Bliley, Daniel Shiwarski, Kritika Chaddha, Devora Cohen-Karni, Adam Feinberg, Pulkit Grover, Aditi Gurkar, Tzahi Cohen-Karni

## Abstract

Electrophysiological recordings provide a key functional readout of cardiac health. Most *in vitro* studies rely on acute measurements rather than recording evolving functional changes. Here, we implement a microelectrode array (MEA) based system for chronic electrophysiological monitoring of human induced pluripotent stem cell-derived cardiomyocyte (iPSC-CM) monolayers. Electrophysiology recordings were maintained for up to 40 days, enabling longitudinal assessment of field potential amplitude, field potential duration, beat frequency, conduction velocity, and waveform morphology. Untreated iPSC-CM monolayers exhibited stable electrophysiological behavior throughout chronic culture. Following exposure to the cardiotoxic chemotherapeutic doxorubicin, chronic recordings captured progressive deterioration in electrical activity, including reduced field potential amplitude, altered repolarization dynamics, increased beat-to-beat variability, and reduced conduction that persisted after drug removal. Electrode-resolved analyses further revealed pronounced progressive spatial variability, highlighting localized responses that would be obscured by population-averaged measurements. Endpoint assays indicated reduced calcium transient amplitude and altered calcium reuptake, decreased mitochondrial membrane potential, and loss of connexin 43, findings consistent with the impaired electrical activity observed in chronic recordings. Together, these results establish chronic electrophysiological monitoring as a powerful approach for resolving the spatiotemporal progression of cardiotoxicity and other dynamic cardiac processes *in vitro*.

## INTRODUCTION

Electrophysiology recordings, e.g. patch-clamp, voltage-sensitive dyes, field effect transistors (FETs) and microelectrode arrays (MEAs), are the current standard approaches for measuring and understanding cardiac electrical function *in vitro*. [1] Patch-clamp electrophysiology recordings provide high-fidelity intracellular recordings from individual cells by measuring transmembrane ionic currents. They are inherently low-throughput and invasive.[2] Voltage-sensitive dyes enable optical mapping of electrical activity by converting changes in membrane potential into fluorescent signals while calcium-sensitive dyes bind to intracellular calcium and emit fluorescent signals. Both require labeling and are limited by phototoxicity and photobleaching.[3] Field effect transistors (FETs) and microelectrode arrays (MEAs) enable label-free, multiplexed intracellular and extracellular field potential recordings. These two approaches have become the standard approach for characterizing the electrical function of cardiac and neural *in vitro* and *in vivo* models.[4–6] Current MEA techniques are commonly used for acute measurements, rather than for chronic monitoring which is key to further understanding progressive disease models. Chronic electrophysiology recordings offer the ability to assess electrical changes over long periods of time in lieu of activity snapshots, providing greater insight into dynamic changes over acute measurements. By quantifying and tracking variations in cardiomyocyte electrical function at the cellular level over extended periods of time, it becomes possible to capture dynamic spatial and temporal patterns that may serve as early indicators of tissue dysfunction. Drug-induced cardiotoxicity provides a clear example where chronically tracking tissue function can substantially enhance our understanding of disease progression, as side-effects are often seen in patients long after treatment concludes.[7]

Doxorubicin, an anthracycline commonly used to treat various cancers, induces cardiotoxicity in approximately 9% of patients. [7, 8] Doxorubicin-mediated cardiomyocyte damage is mainly caused by trapping topoisomerase IIβ in a cleavage complex, generating DNA double-strand breaks and thereby inhibiting replication and repair, in turn leading to cell apoptosis.[7, 9, 10] Additionally, doxorubicin generates reactive oxygen species (ROS), causing oxidative stress which can lead to increased inflammatory cytokines, mitochondrial damage, mishandling of intracellular calcium ions, impaired contractility, and congestive heart failure. [11–14] Chronic electrophysiology monitoring is therefore needed to capture the progressive mechanisms underlying doxorubicin-induced cardiotoxicity, providing an on-demand readout of ion channel dynamics along with indirect assessment of metabolic changes and cell coupling changes. [15] The use of induced pluripotent stem cell (iPSC) derived cardiac cells in *in vitro* models has aided in providing findings relevant to human physiology and has become the gold standard for *in vitro* cardiac models.[16] Acute recordings of iPSC-CM have been extensively demonstrated utilizing two dimensional (2D) and three dimensional (3D) MEA geometries and materials. [17–19]

Electrophysiological recordings offer the ability to assess the electrical signals of individual electrogenic cells, as well as the electrical communication of the network of signals which propagate throughout the tissue chronically and across the lifespan of the cells or tissue. The cardiomyocyte extracellular field potential recorded using passive MEAs reflects Na⁺, Ca²⁺, and K⁺ ionic transients, with the field potential amplitude (FPA) primarily associated with Na⁺ influx, and the field potential duration (FPD) governed by the interplay of Ca²⁺ and K⁺ currents (Supplemental Figure 5). [20] Multicellular (syncytium) behavior can be quantified through conduction velocity, calculated via the time latency between field potential onset at each recording site (electrode). [21, 22]

Long-term electrophysiological monitoring of engineered cardiac tissues presents a technical challenge because recording interfaces must maintain stable electrical coupling while accommodating tissue growth, remodeling, and continuous spontaneous contraction. Two-dimensional MEA platforms have been utilized to study long-term cardiac electrophysiology in a handful of studies, utilizing passive electrodes to record longitudinally and/or localized optoporation for action potential recordings to record for 45 days.[23, 24] To better replicate the native myocardial microenvironment, several groups have developed MEAs compatible with three-dimensional cardiac tissues, either encapsulating the tissue or embedded within it, using designs such as ultra-thin meshes, foldable and self-curling structures.[4, 5, 25] A shell MEA was utilized to chronically record for 9 days from a cardiac spheroid [26] representing a step away from acute monitoring toward long-term, chronic evaluations of *in vitro* models. (See Table 1 in Supplementary Information for a comprehensive list of chronic/long-term electrophysiology platforms) These studies demonstrate significant advances in interfacing with engineered cardiac tissues. Nevertheless, recording durations remain substantially shorter than the weeks-to months-long timescales required to study tissue maturation, chronic drug exposure, or progressive cardiotoxicity. We set out to implement a platform capable of chronic electrophysiological recordings from two-dimensional (2D) cardiac monolayers over the extended timescales needed to investigate long-term physiological remodeling and disease progression.

**Table 1:**
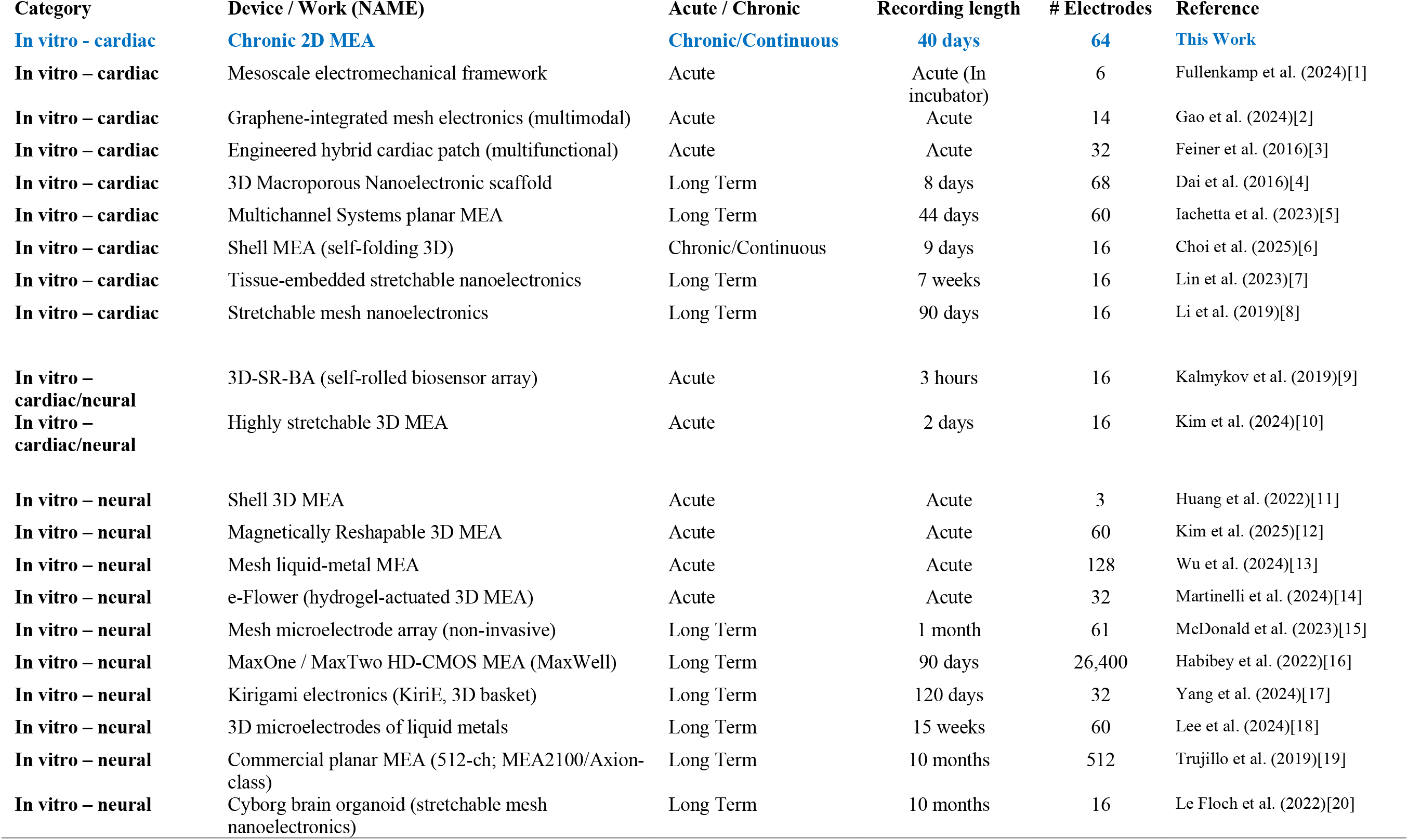
Notable Acute, Long Term (Non-Chronic, and Chronic Electrophysiology Platforms.

Here, we report chronic 2D recordings at high spatial resolution in an *in vitro* disease model spanning over one month with hourly to daily temporal resolution. In contrast to previous studies,[8, 23, 25, 27, 28, 29] we eliminate the need for repeated acute recordings, which typically yield discrete timepoints across the scale of minutes or hours, and demonstrate a continuous chronic monitoring approach that fully captures evolving temporal shifts in cardiac electrical behavior of the same specimen in the same locations over an extended time period (from recording onset through 40 days *in vitro*). [26] We report an integrated chronic electrophysiology platform for iPSC-derived cardiomyocytes that enables longitudinal detection of progressive dysfunction caused by doxorubicin that cannot be resolved by conventional acute assessments via extracellular field potentials (Figure 1). While expected variability in untreated tissues was observed, doxorubicin-administered samples exhibit a progressive increase in variance during functional decline and eventual arrest of electrophysiological behavior.

**Figure 1:**
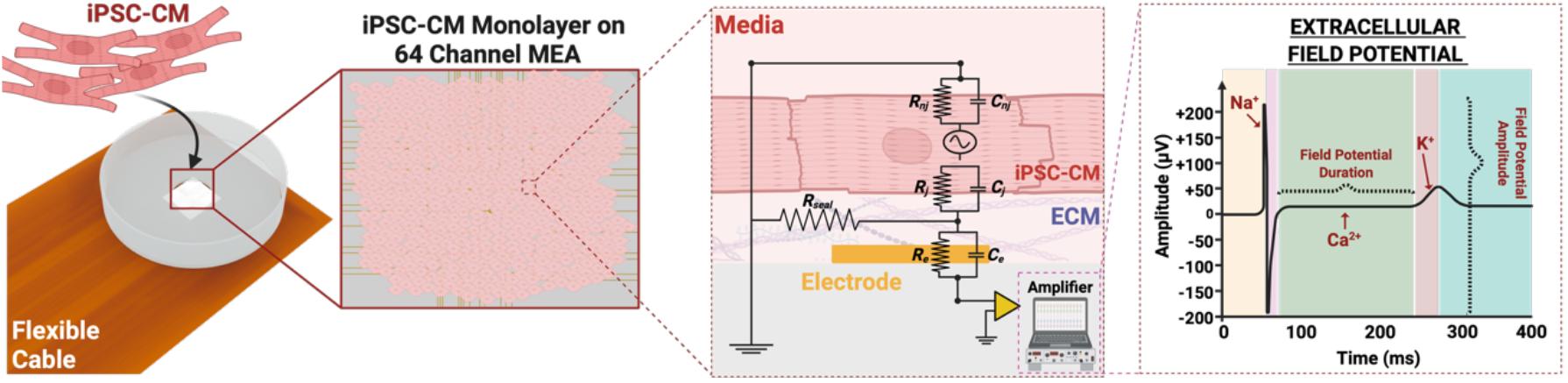
Chronic Electrophysiology experimental overview. Induced pluripotent stem cell (iPSC) derived cardiomyocytes are seeded onto 64-channel microelectrode arrays (MEAs). Extracellular field potentials are recorded from each electrode to assess individual cardiomyocyte electrical behavior.

Electrophysiological measurements were complemented by endpoint functional and structural assays to assess cellular processes underlying single-cell electrophysiological function, while structural analyses evaluated the tissue architecture and intercellular connectivity that govern monolayer conduction velocity.[15] Together, the implementation of chronic electrophysiology recording of iPSC-derived cardiomyocytes resolves the temporal progression of doxorubicin-induced cardiotoxicity and could be extended to other cardiac disease models.

## RESULTS AND DISCUSSION

Chronic electrophysiology recordings allow continuous monitoring of electrical activity over extended periods, rather than relying on brief snapshots of activity, enabling better characterization of gradual, progressive changes. Such high spatiotemporal resolution recordings enable researchers to analyze disease progression through two lenses: (1) comparing and tracking variability at the cellular level (at the microelectrode resolution) at any given time and (2) comparing multicellular-level changes over time by analyzing wave propagation (microelectrode array level).

### Microelectrode Arrays for Long Term Recordings

Two-dimensional passive microelectrode arrays (MEAs) allow spatial assessment of electrical behavior across a monolayer of cells by recording extracellular field potentials (Figure 2A). A 64-channel MEA was fabricated and packaged (for details see Materials and Methods) to record cardiac monolayer electrophysiology. Once seeded, cardiomyocytes form a continuously beating monolayer. Electrical signals were recorded with high spatial (250 µm pitch) and temporal resolution (50 µsec) with once hourly recording frequency. The fully assembled device (Supplemental Figure 2-3) has an 8 × 8 array of 50 µm × 50 µm electrodes that was fabricated using standard photolithography techniques (for the 2D MEA design see Supplemental Figure 1, for additional fabrication details see the Materials and Methods).

**Figure 2:**
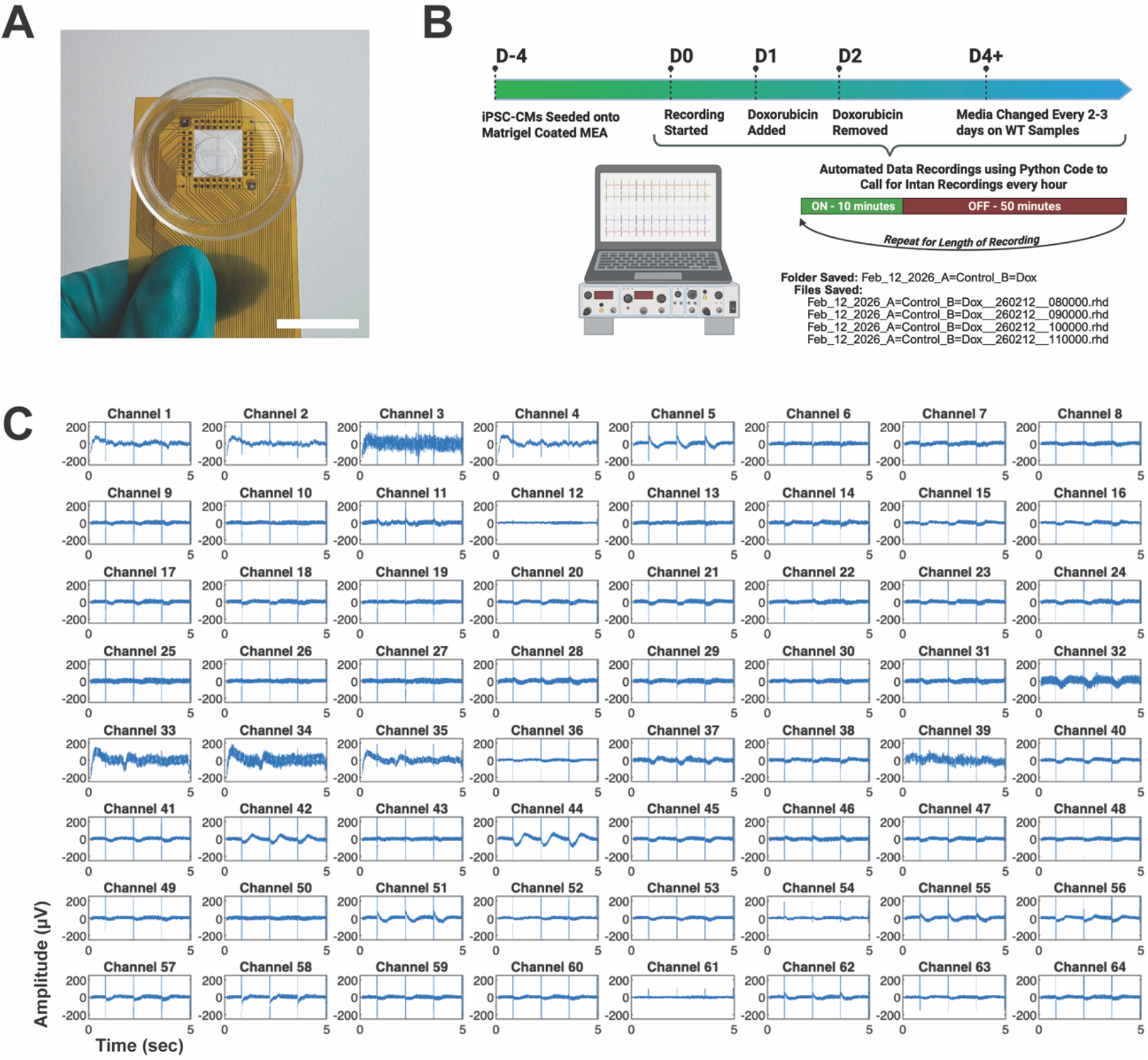
Chronic Electrophysiology Recording System. **(A)** Fully assembled 64-channel microelectrode array (MEA). Scale bar = 20 mm. **(B)** Experimental and recording timeline. **(C)** Representative 5 second trace of recording from untreated induced pluripotent stem cell derived cardiomyocytes (iPSC-CM).

To characterize the electrochemical properties of the microelectrodes, cyclic voltammetry (CV) and electrochemical impedance spectroscopy were performed in a three-electrode electrochemical cell (Supplemental Figures 2 and 3) (for details see Materials and Methods). [5] The microelectrodes were modified with either platinum black or poly(3,4-ethylenedioxythiophene):poly(sodium 4-styrenesulfonate) (PEDOT:PSS) to increase surface area, reduce impedance, and improve recording quality.[19,20] The Au microelectrodes weakly resolve oxidative and reduction peaks in potassium hexacyanoferrate(III), but fully resolve these peaks after modification with platinum black or PEDOT:PSS. The measured impedance at 1kHz was 130 ± 40.50 kΩ (n=10), 8.68 ± 5.35 kΩ (n=10), and 8.71 ± 1.93 kΩ (n=10) for bare Au, PEDOT:PSS-modified Au, and platinum black-modified Au electrodes, respectively, demonstrating reduced impedance following both surface modifications consistent with previous reports.[5, 30] The improved electrochemical activity and reduced impedance have been previously reported to yield higher signal amplitudes in electrophysiology recordings.[31]

**Figure 3:**
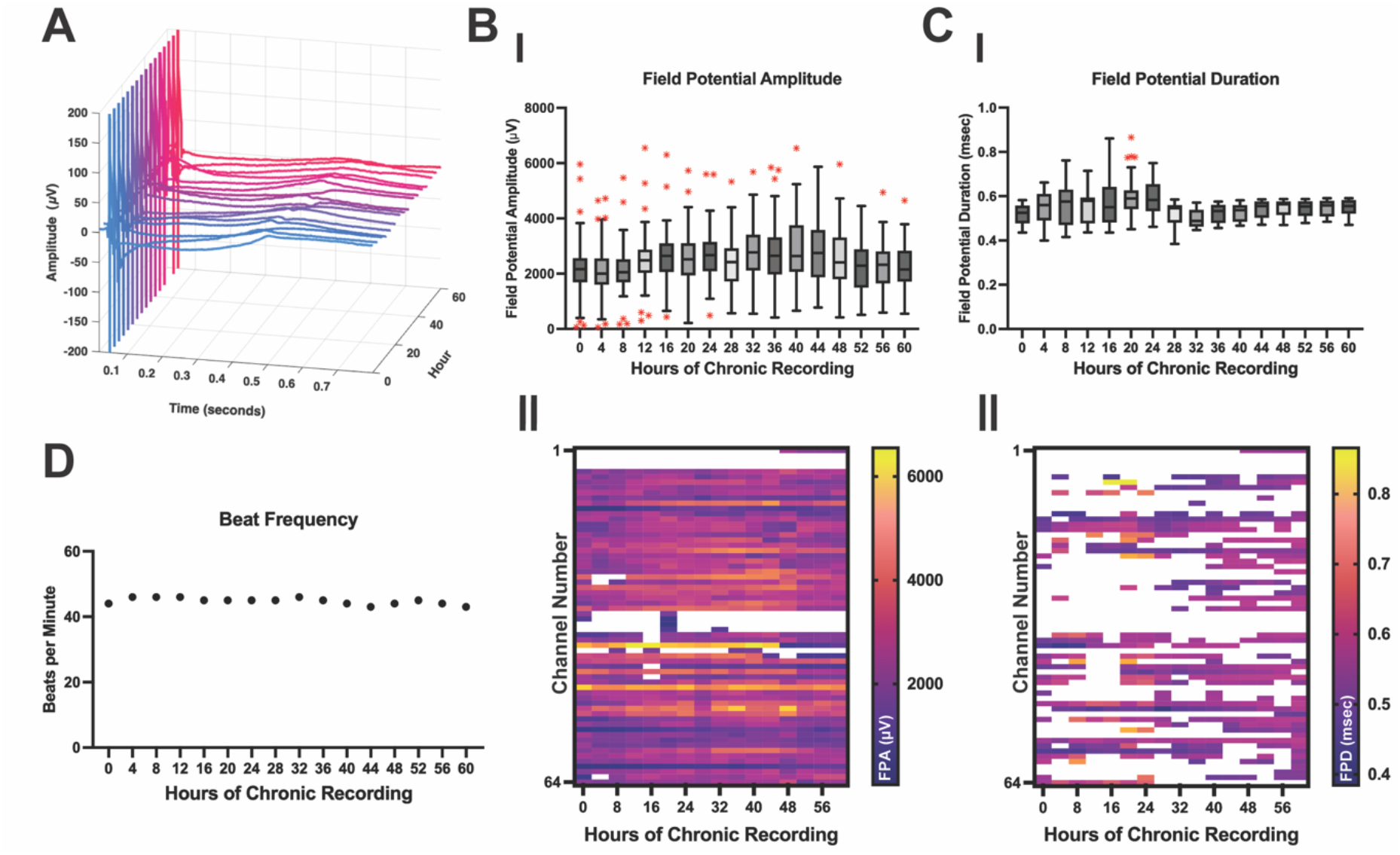
Single Electrode Level Metrics from Untreated iPSC-CM. **(A)** Representative extracellular field potential from a single channel over 60 hours of chronic recordings. **(B)** Field Potential Amplitude (I) Boxplot of field potential amplitude across 60 hours of chronic recording, with whiskers extending to 1.5× IQR and outliers shown in red. (II) Heatmap shows individual electrode amplitude values for each day of recording, white areas indicate no field potentials recorded from that channel at that timepoint. **(C)** Field Potential Duration (I) Boxplot of field potential duration across 60 hours of chronic recording, with whiskers extending to 1.5× IQR and outliers shown in red. (II) Heatmap shows individual electrode duration values for each day of recording, white areas indicate no potassium peak could be identified in the data. **(D)** Beat frequency over 60 hours of chronic recording.

### Long-term Single-Cell Electrophysiological Changes due to Doxorubicin Exposure and DNA Damage

Chronic electrophysiology recordings provide a robust approach to track cardiomyocyte electrical behavior at both cellular and tissue scales, revealing spatial and temporal changes in key ionic transients over time. Long-term electrophysiology recordings were used to assess the uniformity and consistency of untreated iPSC derived cardiac monolayers and track the development of doxorubicin induced cardiotoxicity following a 24-hour exposure to 0.5 µM doxorubicin (for details see Materials and Methods and Supplemental Figure 4). The extracellular field potentials recorded reveal three main transients across the cell membrane over time: Na^+^(upstroke), Ca^2+^(plateau), and K^+^(repolarization).[20] iPSC-CM monolayers were continuously maintained in an incubator while automated 10-minute recordings were taken every hour, as outlined in Figure 2B. A representative 64-channel recording is shown in Figure 2C, illustrating the spatial distribution of electrical activity across the MEA.

Chronic electrophysiological assessment of untreated iPSC-CMs was performed to establish baseline behavior and inherent variability over time, ensuring that electrophysiological trends seen in disease models could be attributed to disease pathologies. While variability in behavior is observed over time, stable behavior in this context refers to electrical activity that remains within the range of reported values for untreated iPSC-CM and shows no directional drift across the recording period, consistent with the intrinsic beat-to-beat variability known of the human heart. [32–34]

Analysis of individual field potentials yields two key parameters: field potential amplitude, which correlates with the magnitude of the Na^+^ current upstroke and downstroke, and field potential duration, which correlates with the duration of the Ca^2+^ plateau. (See Materials and Methods and Supplemental Figure 6 for analysis details.) Both parameters can be tracked over space and time and used to identify changes in cellular electrical behavior. Chronic single-electrode measurements captured from a representative untreated iPSC-CM monolayer over 60 hours illustrate the long-term electrophysiological behavior of healthy recordings (Figure 3). A representative extracellular field potential from a single cell (single electrode) demonstrates consistent waveform morphology throughout the 60-hour period displayed (Figure 3A). Extracellular field potentials across all active channels were analyzed from each timepoint displayed for extraction of single-cell electrophysiology metrics including FPA, FPD, and beat frequency (Figure 3 encompasses 52-57 active channels with n = 440-460 field potentials analyzed at each timepoint). Supplemental Figures 7 –10 present complete 20–40 day untreated iPSC-CM recordings, demonstrating that electrophysiological activity remained stable throughout extended chronic monitoring.

Proper function of Na^+^ ion channels, visible via FPA, is essential for the initiation of a field potential, rapid cell depolarization and the coordination of contraction at the cellular, tissue, and organ scales. Untreated iPSC-CM recordings showed no significant trend change in FPA across the recording ranging between a minimum of 2104.0 ± 988.7 µV (hour 4 of recording, n=52 channels, 460 field potentials per channel) to a maximum of 2852.0 ± 1120.0 µV (hour 40 of recording, n=56 channels, 440 field potentials per channel) (Figure 3BI), indicating consistent Na^+^ ion channel kinetics over time and matching values reported in previous studies. [5, 6, 35, 36] Similarly, analysis of the Na^+^ peak width, upstroke and downstroke velocities revealed no significant changes in behavior over the time frame of the recording (Supplemental Figures 11–16), indicating stable depolarization and repolarization dynamics across the chronic recording period.

To visualize the spatial and temporal evolution of electrophysiological changes across the monolayer, the FPA quantified for each electrode was displayed as heatmaps, where each row represents an individual electrode and each column represents a longitudinal timepoint (Figure 3BII). This representation enables rapid assessment of both temporal changes at an electrode and spatial variability across the monolayer. As evident in these heatmaps, the FPA exhibited both spatial (variation along each row/between electrodes) and temporal (variation along each column/between time points) heterogeneity across the healthy monolayer, demonstrating the importance of accounting for intrinsic variability when interpreting electrophysiological changes in both healthy and damaged tissues.

Field potential duration reflects cardiac repolarization and is primarily influenced by Ca^2+^ and K^+^ ion channel activity. FPD remained consistent during continuous 60-hour monitoring. The field potential duration varied between a minimum of 499.8 ± 43.9 ms (hour 32 of recording, n=23 channels, 460 field potentials per channel) and a maximum of 602.7 ± 100.0 ms (hour 24 of recording, n=30 channels, 450 field potentials per channel), remaining within the expected physiological range and showing no progressive changes indicative of cardiac stress. [5, 6, 35, 36] FPD values were similarly visualized as electrode-resolved heatmaps (Figure 3CII). Although spatial and temporal variability was observed, FPD remained within the expected physiological range with no variation outside of expected values for untreated iPSC-CM FPD, indicating healthy repolarization dynamics throughout the culture period. Intrinsic beat frequency remained stable over 60 hours (Figure 3D) and beat rate variability stayed within physiologically expected values across the recording (Supplemental Figures 17–18). Although individual electrodes exhibited expected spatial and temporal variability, the overall electrophysiological behavior of the untreated iPSC-CM monolayer remained stable at the multicellular level, establishing a baseline against which functional changes associated with cardiac stress, injury, or drug exposure can be distinguished. (See Materials and Methods for analysis details.)

While untreated iPSC-CM monolayers demonstrated stable electrophysiological behavior during chronic monitoring, the ability to capture progressive functional changes following cardiac injury or stress represents the key advantage of chronic recordings. Doxorubicin exposure can rapidly alter iPSC-CM function, with effects persisting beyond the initial treatment period. [37–39] Based on the half-life and peak concentration of doxorubicin in plasma after chemotherapeutic administration, [40, 41] the cardiac monolayers were exposed to 0.50 µM doxorubicin for 24 hours, followed by drug removal and continued electrophysiological monitoring to assess the long-term consequences of transient doxorubicin exposure on cardiac function (Figure 4).

**Figure 4:**
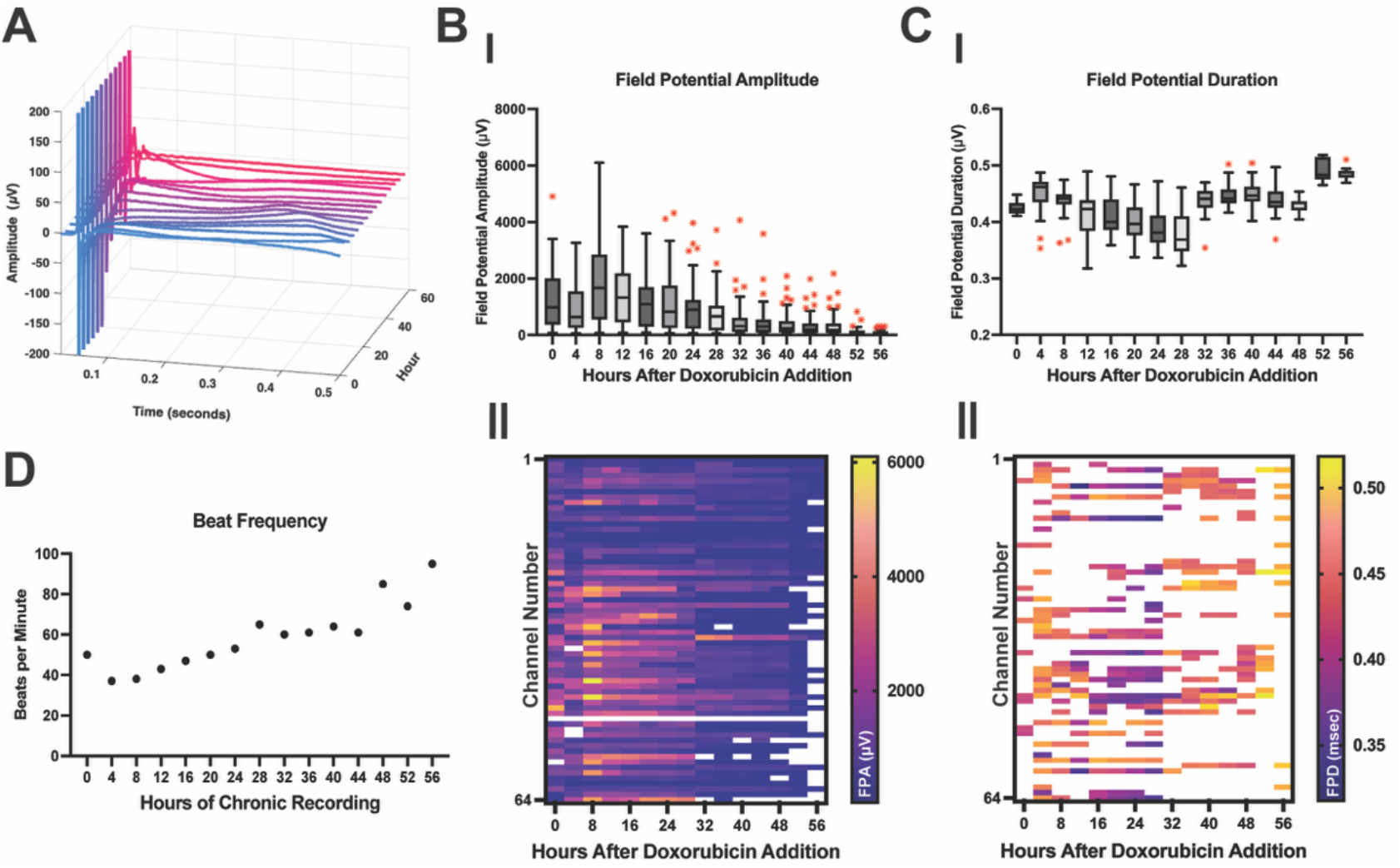
0.50 μM Doxorubicin Treated iPSC-CM Chronic Recording. **(A)** Representative extracellular field potential across all hours of recording. (B) Field potential amplitude, (I) Boxplot of field potential amplitude across 56 hours of chronic recording, with whiskers extending to 1.5× IQR and outliers shown in red. (II) Heatmap shows individual electrode amplitudes values for every 4 hours of recording, white areas indicate no field potentials recorded from that channel at that timepoint.. (C) Field potential duration, (I) Boxplot of field potential duration across 56 hours of chronic recording, with whiskers extending to 1.5× IQR and outliers shown in red. (II) Heatmap shows individual electrode durations values for every 4 hours of recording, white areas indicate no potassium peak could be identified in the data. (D) Beat frequency across 56 hours of chronic recording.

The extracellular field potential from a representative recording channel is shown to illustrate progressive changes in waveform morphology over the 56-hour recording (Figure 4A). Changes in morphology reflect alterations in the underlying ionic currents and support the quantitative assessment of field potential amplitude, field potential duration, and additional electrophysiological parameters (beat rate, beat rate variability, sodium peak width, upstroke and downstroke velocity). Field potential amplitude was evaluated to quantify changes in depolarization capacity during doxorubicin-induced injury, as decreases in signal magnitude can indicate altered Na⁺ channel activity and reduced cardiomyocyte excitability.[8, 42] Average field potential amplitude progressively decreases (Figure 4BI). In individual recordings, beat rate became abnormal (either increased or decreased) before electrical activity ceased entirely, with complete cessation of beating occurring between 28 – 60 hours (Supplemental Figures 19-22). Recordings were lost as cardiomyocytes died, metric values at later timepoints reflect only the channels that remained active. The FPA decreased from 1216.0 ± 1004 µV (hour 0, n = 62 channels, 500 field potentials per channel) to 107.3 ± 78.5 µV (hour 56, n = 32 channels, 950 field potentials per channel) with the statistically significant decline occurring 28 hours after the start of exposure (24 hours of exposure, 4 hours after removal). Visualizing the spatial and temporal changes via a heatmap of the FPA (Figure 4BII), a consistent decline in FPA across channels was observed, providing enhanced temporal and spatial resolution to previous endpoint-based studies investigating doxorubicin induced cardiotoxicity. [8, 35, 36, 43] Additional waveform analyses demonstrated increased sodium peak width and decreased upstroke and downstroke velocities over time (Supplemental Figures 23–25), consistent with progressive alterations in cardiomyocyte excitability and conduction. These functional changes may reflect impaired sodium channel activity and disruption of ion homeostasis, including altered Na⁺/K⁺ ATPase, Na⁺/Ca²⁺ exchanger (NCX), and L-type calcium channel function during doxorubicin-induced stress [44, 45]

FPD was monitored longitudinally as an extracellular measure of action potential duration, providing insight into repolarization abnormalities arising from altered calcium and potassium channel activity. FPD exhibited dynamic fluctuations throughout the recording period ranging from a minimum of 380.3 ± 3.9 msec (hour 28, n = 25 channels, 650 field potentials per channel) to a maximum of 490.9 ± 20.5 msec (hour 52, n = 8 channels, 740 field potentials per channel), with no consistent significant overall trend in change across the total time frame (Figure 4CI). FPD initially decreased over the first 28 hours and then increased at later time points, until activity ceased in individual recordings between 28 and 60 hours (Supplemental Figures 19-22). Because recordings were progressively lost over this period, later values reflect only the channels that remained active. [46, 47] The shifting trends in FPD may indicate inconsistent Ca^2+^ reuptake.[48] This irregular behavior is also consistent with previously described reduction in ATP or decreased K^+^ or NCX channel activity.[11, 15, 49, 50]

Beat frequency progressively increased over the course of the recording (Figure 4D), indicating changes in pacemaker activity and in Ca^2+^ and K^+^ handling.[11] Doxorubicin addition increases beat-to-beat variability beyond what is typically observed in iPSC-CM monolayers (Supplemental Figure 26), indicating increased fluctuations in rhythm and further supporting progressive electrophysiological dysfunction following doxorubicin exposure.

Chronic electrophysiological monitoring revealed the functional progression of doxorubicin-induced cardiac dysfunction. Subsequent single-cell analyses provided mechanistic insight into the cellular alterations associated with this evolving electrophysiological phenotype.

Cardiac electrophysiology is highly dependent upon intracellular calcium handling and the flux of calcium into and out of the cell.[15] The field potential duration taken from MEA recordings directly parallels the calcium peak width taken from calcium imaging. Assessment of calcium handling (Figure 5) was used to further evaluate calcium handling post doxorubicin exposure. Calcium handling was assessed in parts, through the unprocessed calcium transients from the fluorescence intensity in one section of the calcium image and secondly through the subsequent normalized calcium transients. A representative trace of calcium fluorescence intensity (Figure 5AI) shows a higher amplitude of the calcium intensity for untreated iPSC-CM compared to iPSC-CM administered 0.50 µM of doxorubicin. Peak amplitude is quantified (Figure 5AII) and a significant decrease (∼4.66x reduction) in peak amplitude is determined after 0.50 µM doxorubicin addition. This reduction in amplitude is attributed to decreased uptake of calcium into the cytosol.[15]

**Figure 5:**
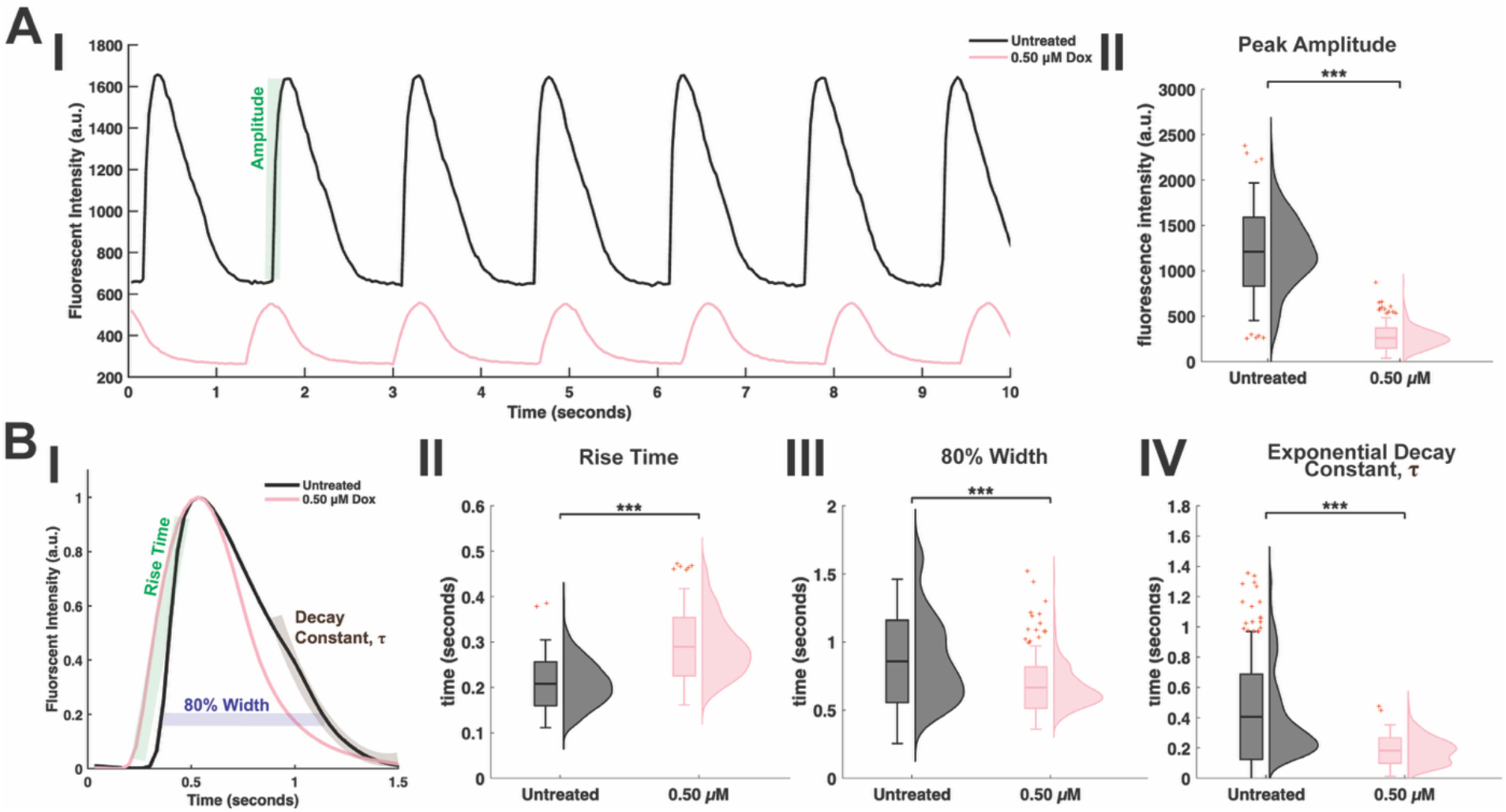
Doxorubicin Effects on Calcium Handling. **(A)** Calcium handling metrics taken from raw data of untreated and 0.50 μM doxorubicin exposed iPSC-CM calcium transients. **(I)** Representative calcium trace from untreated and 0.50 μM iPSC-CM after 24 hours of doxorubicin administration followed by 4 days in maintenance media. **(II)** Peak amplitude of untreated and 0.50 μM doxorubicin exposed iPSC-CM after 24 hours of doxorubicin administration followed by 4 days in maintenance media. **(B)** Calcium handling metrics taken for amplitude normalized calcium transients. **(I)** Representative normalized calcium transient for untreated and 0.50 μM doxorubicin exposed iPSC-CM **(II)** Rise time from 10% to 90% of peak height of untreated and 0.50 μM iPSC-CM after 24 hours of doxorubicin administration followed by 4 days in maintenance media. **(III)** Peak width at 80% of peak height of untreated and 0.50 μM doxorubicin exposed iPSC-CM after 24 hours of doxorubicin administration followed by 4 days in maintenance media. **(IV)** Exponential decay fit constant, *τ*, from the second half of the decay of untreated and 0.50 μM doxorubicin exposed iPSC-CM after 24 hours of doxorubicin administration followed by 4 days in maintenance media. 4 individual samples were imaged for each condition with each sample. Statistical significance was determined with a one-way ANOVA with Tukey’s multiple comparison, *** indicates statistical significance with a p-value < 0.001.

The normalized calcium transients are used to compare the shape of the calcium transient and associated ion channel dynamics.[51] When normalized, representative untreated and doxorubicin-administered iPSC-CM calcium transients show a clear morphological difference. (Figure 5BI). L-type Ca^2+^ channel activity and calcium induced calcium release during depolarization can be assessed via the peak rise time. The peak rise time increased significantly after doxorubicin administration (∼1.39x increase) (Figure 5BII), indicating a slower release of calcium into the cytosol via L-type Ca^2+^ channels and from the sarcoplasmic reticulum via the ryanodine receptor (RyR). Calcium release from and reuptake into the sarcoplasmic reticulum has also been correlated to the calcium transient duration at 80% of reuptake.[51] The calcium transient duration was assessed by calculating the width of the calcium transient at 80% of repolarization (Figure 5BIII). This decrease (∼1.29x reduction) was a significant reduction in width, indicating arrhythmogenic behavior and has been correlated with early afterdepolarizations.[52] Sarcoplasmic/endoplasmic reticulum Ca2+-ATPase (SERCA) and sodium-calcium exchanger (NCX) activity can be assessed via the exponential decay of the calcium repolarization.[51] The exponential decay constant, tau (τ), for the second half of the decay fit was therefore calculated to indicate how quickly SERCA and NCX remove calcium from the cell (Figure 5BIV).[51] The decay constant was observed to significantly decrease after 0.50 µM doxorubicin exposure (∼2.22x reduction). This decrease in τ indicates faster decay, and therefore in the context of calcium repolarization, faster removal of calcium from the cell as the calcium transient fluorescent intensity returns to baseline.

Doxorubicin exposure significantly altered calcium handling dynamics as evidenced by the significant changes in amplitude, rise time, peak width, and decay kinetics. These alterations suggest disruption to key ion channels, pumps, and exchangers which facilitate proper excitation and contraction of a cardiomyocyte. The altered calcium dynamics may underlie the fluctuating field potential duration observed in electrophysiology recordings from doxorubicin-administered iPSC-CM where FPD is seen to be highly variable compared to untreated iPSC-CM.

Mitochondria play a key role in the flux of calcium into and out of the cell due to their involvement in excitation-contraction coupling and ATP synthesis. [15, 53] Doxorubicin has been found to directly interact with mitochondria via binding with phospholipid cardiolipin in the inner mitochondrial membrane, leading to accumulation of doxorubicin and inhibition of complexes I, II, and IV of the electron transport chain.[49, 54] Additionally, multiple studies have found doxorubicin to dysregulate the GTPase family of proteins (Drp1, Opa1, and Mfn2), specifically downregulation of fusion proteins (Opa1 and Mfn2), and upregulation of fission proteins (Drp1), resulting in overall increased fragmentation of the mitochondrial network and decreased ATP production via oxidative phosphorylation.[46, 55]

Mitochondrial damage can be confirmed directly by assessing mitochondrial activity.[11, 56] The negative charge, created by the proton pumps of the electron transport chain, creates the mitochondrial membrane potential which was fluorescently labeled here. [57] Mitochondrial membrane potential (MMP) was imaged to assess mitochondrial functionality and mitochondrial network morphology as shown in Figure 6A (see Materials and Methods for details). A ca. 1.5x decrease in MMP was seen between untreated iPSC-CMs and the 0.10 µM doxorubicin treated iPSC-CMs, and a ca. 2.7x decrease in MMP was seen between untreated and 0.50 µM doxorubicin treated iPSC-CMs, indicating a statistically significant dose-dependent relationship between doxorubicin and MMP (Figure 6B). Doxorubicin-induced mitochondrial dysfunction may further compound the calcium handling disruptions described above (Figure 5). Mitochondria contribute to intracellular Ca^2+^ buffering, and both membrane potential and network integrity influence this capacity. The decreased MMP and fragmented mitochondrial network observed after doxorubicin exposure may therefore impair Ca^2+^ buffering and alter calcium handling. Separately, dysfunctional mitochondria exhibit impaired calcium uptake, further reducing cytosolic calcium clearance. Both effects occur in a dose-dependent manner. This interaction further exemplifies the interconnectedness between electrophysiology, calcium-handling, and metabolic behavior within cardiomyocytes.[11, 58, 59]

**Figure 6:**
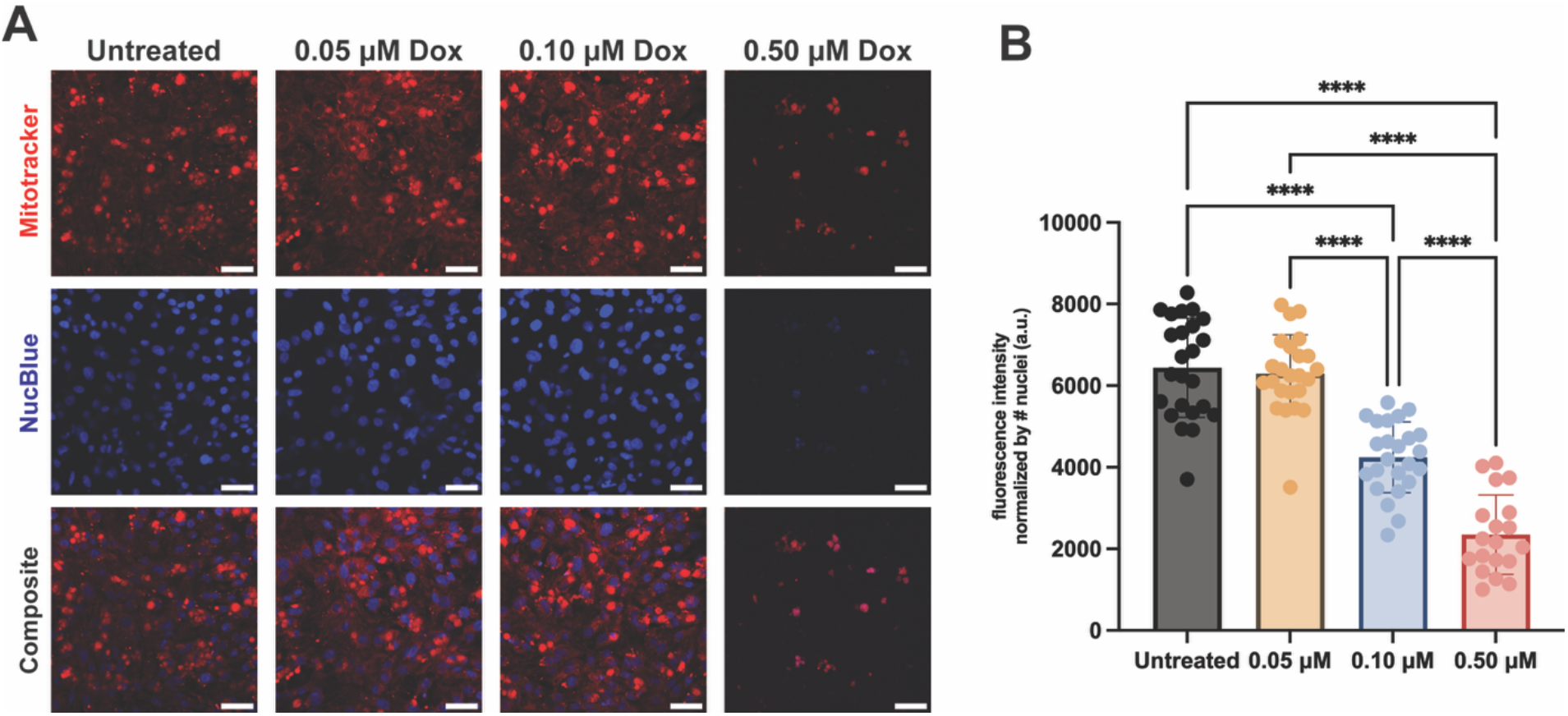
Doxorubicin Effect on Mitochondrial Membrane Potential. **(A)** Representative images of mitochondrial membrane potential using Mitotracker Red CXMRos in untreated and doxorubicin treated (0.05 µM, 0.10 µM, 0.50 µM) iPSC-CMs. **(B)** Quantification of the fluorescence intensity of mitochondrial membrane potential normalized per image by the nuclei count. 3 samples were imaged per condition, with 4 fields of view analyzed per sample. Statistical significance was determined with a one-way ANOVA with Tukey’s multiple comparison, ns indicates non-significance, **** indicates statistical significance with a p-value < 0.0001.

This doxorubicin-induced decline in MMP, indicative of mitochondrial dysfunction, has also been linked to increased concentrations of reactive oxygen species which are created in the mitochondria after doxorubicin exposure. [60] Mean fold change in ROS levels after doxorubicin exposure was measured via fluorescence imaging, confirmation a ca. 1.5 and 2 increase in ROS over time (Supplemental Figure 29), and a ca. 1.5, 4.5, and 7.4 with increasing concentrations (0.05, 0.10, 0.5 uM) of doxorubicin (Supplemental Figure 30).

### Network (Multi-Cellular) Level Analysis

While single-electrode analyses resolve temporal changes at the single-cell level, multicellular-level assessment reveals how coordinated behavior across the monolayer breaks down during doxorubicin exposure.

As doxorubicin-induced cellular damage accumulates, corresponding changes in electrical behavior emerge, capturing the temporal progression of cardiotoxicity. Looking at the entire monolayer, the conduction velocity and propagation direction can be assessed over time (Figure 7). Time latency across the monolayer is determined based on the Na^+^ peak onset. An isochronal map can be plotted to visualize the wave propagation across the monolayer. (See Materials and Methods and Supplemental Figure 28 for analysis details.) Figure 7A shows the conduction velocity over 60 hours for untreated and doxorubicin exposed iPSC-CM. The conduction velocity of untreated iPSC-CM varies between a minimum of 6.66 ± 0.433 cm/s (hour 12, n=12 beats) to a maximum of 18.74 ± 1.822 cm/s (hour 44, n=12 beats) across the 60-hour recording subset. The untreated iPSC-CM isochronal maps show time latency and propagation every twelve hours, demonstrating changes in the monolayer as time progresses, consistent with literature (Figure 7BI).[5, 26] Accompanying quiver maps were generated to display the direction and magnitude of propagation across the monolayer, providing a spatially resolved representation of conduction dynamics across the sample (Supplemental Figure 27). Changes in iPSC-CM conduction velocity and network dynamics are observed following doxorubicin exposure (Figure 7 AII, BII). The initial conduction velocity significantly decreases from 7.04 ± 0.59 (n=10 beats) to 3.63 ± 1.79 cm/s 32 hours after doxorubicin exposure (24 hours of exposure, 8 hours after removal). The slowest conduction velocity of 1.818 ± 1.30 cm/s was measured at 52 hours after doxorubicin exposure (n=10 beats) (Figure 7AI). Decrease in conduction velocity has been reported in both mouse heart models and iPSC-CM cardiac models after doxorubicin exposure.[61] This decrease can be attributed to changes in gap junction expression and organization, as well as changes in calcium handling.[62, 63] The isochronal maps of doxorubicin treated iPSC-CM (Figure 7BII) show how the propagation across the monolayer changes over time and displays a shift in the direction of average propagation, possibly indicating monolayer disruption. By hour 40, the signal propagation wave is seen to diverge, a phenomenon which has been reported to increase with doxorubicin due to downregulated and delocalized gap junctions, reducing electrical coupling between cells.[62] This decrease in conduction velocity and shifts in propagation direction indicate network instability related to cell-cell coupling, most likely due to doxorubicin-induced ROS generation causing DNA damage, which in turn dysregulates gap junction protein expression and disrupts intercellular coupling.[62]

**Figure 7:**
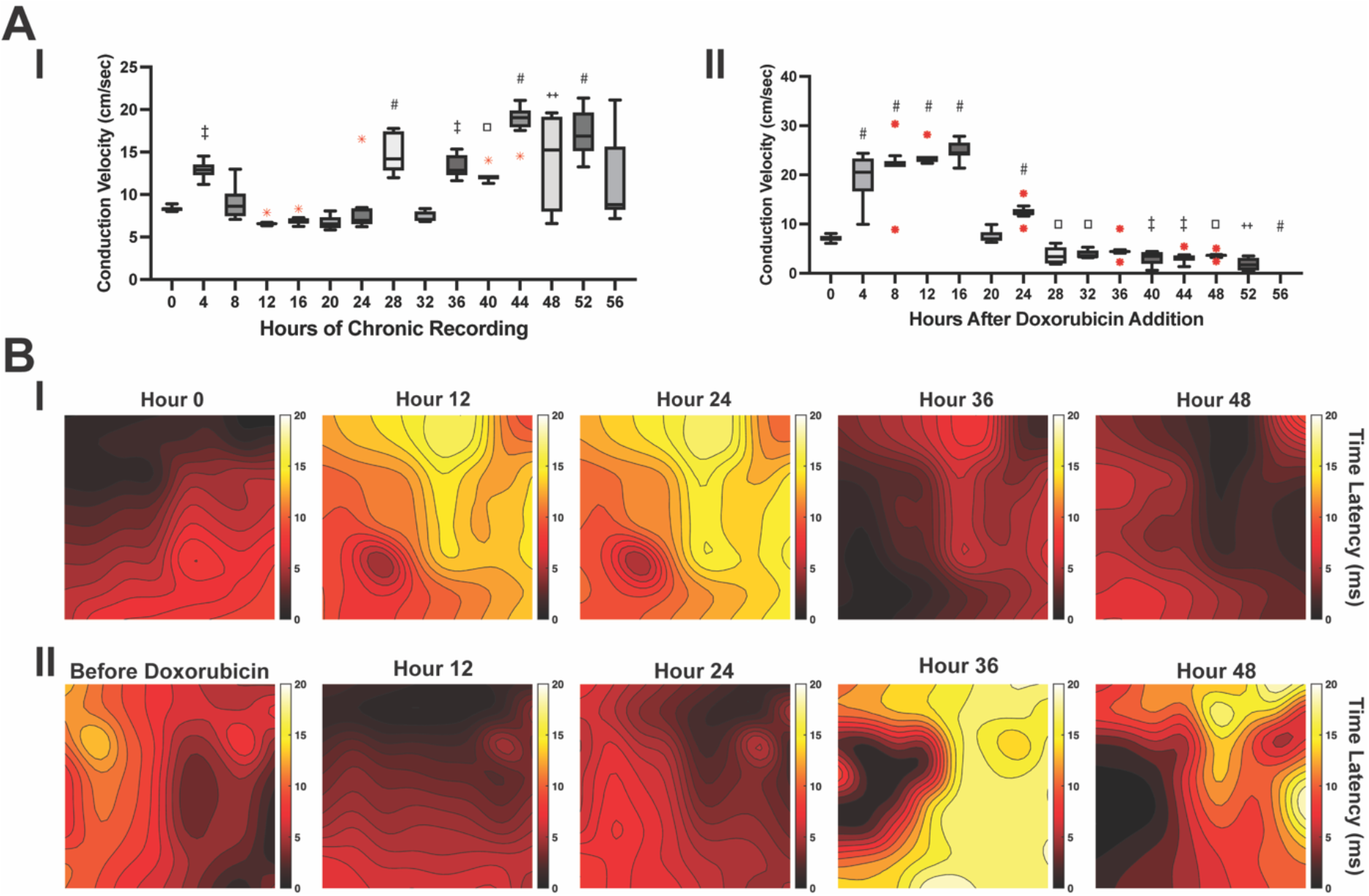
Conduction velocity dynamics across untreated and doxorubicin exposed iPSC-CM over time. **(A)** Conduction velocity every 4 hours across the monolayer. **(I)** untreated iPSC-CM monolayer **(II)** iPSC-CM exposed to 0.5 µM doxorubicin. **(B**) Isochronal maps every 12 hours of recording based on the time latency at each electrode **(I)** untreated iPSC-CM monolayer **(II)** iPSC-CM exposed to 0.5 µM doxorubicin. Statistical significance was determined with a one-way ANOVA with Tukey’s multiple comparison, ◻ indicates statistical significance with a p-value < 0.05, **‡** indicates statistical significance with a p-value < 0.01. **++** indicates statistical significance with a p-value < 0.001, **#** indicates statistical significance with a p-value < 0.0001.

Coordinated cardiac function relies on both electrical coupling and structural organization. Changes in cardiomyocyte architecture and junctional integrity were assessed following doxorubicin exposure. A major driver of cell-cell communication and connectivity in cardiomyocytes is gap junctions between cardiomyocytes. Gap junctions connect cells via intercellular channels made of connexin proteins, which allow the movement of ions between the cytosol of two adjacent cells and therefore electrically couple cells.[64] Connexin 43 (Cx43) is the major component of gap junctions and can be seen to decrease in expression after doxorubicin exposure (Figure 8A). 5 days post doxorubicin exposure, Cx43 expression was significantly reduced in the 0.10 µM group by 19% compared to the untreated samples and by 53% in the 0.50 µM group compared to untreated iPSC-CM (Figure 8B). This reduction in Cx43 expression is consistent with the reduction in conduction velocity seen in chronic electrophysiology recordings, as reduced gap junction coupling limits the efficient propagation of depolarization between cardiomyocytes.[65] (See Materials and Methods for details.)

**Figure 8:**
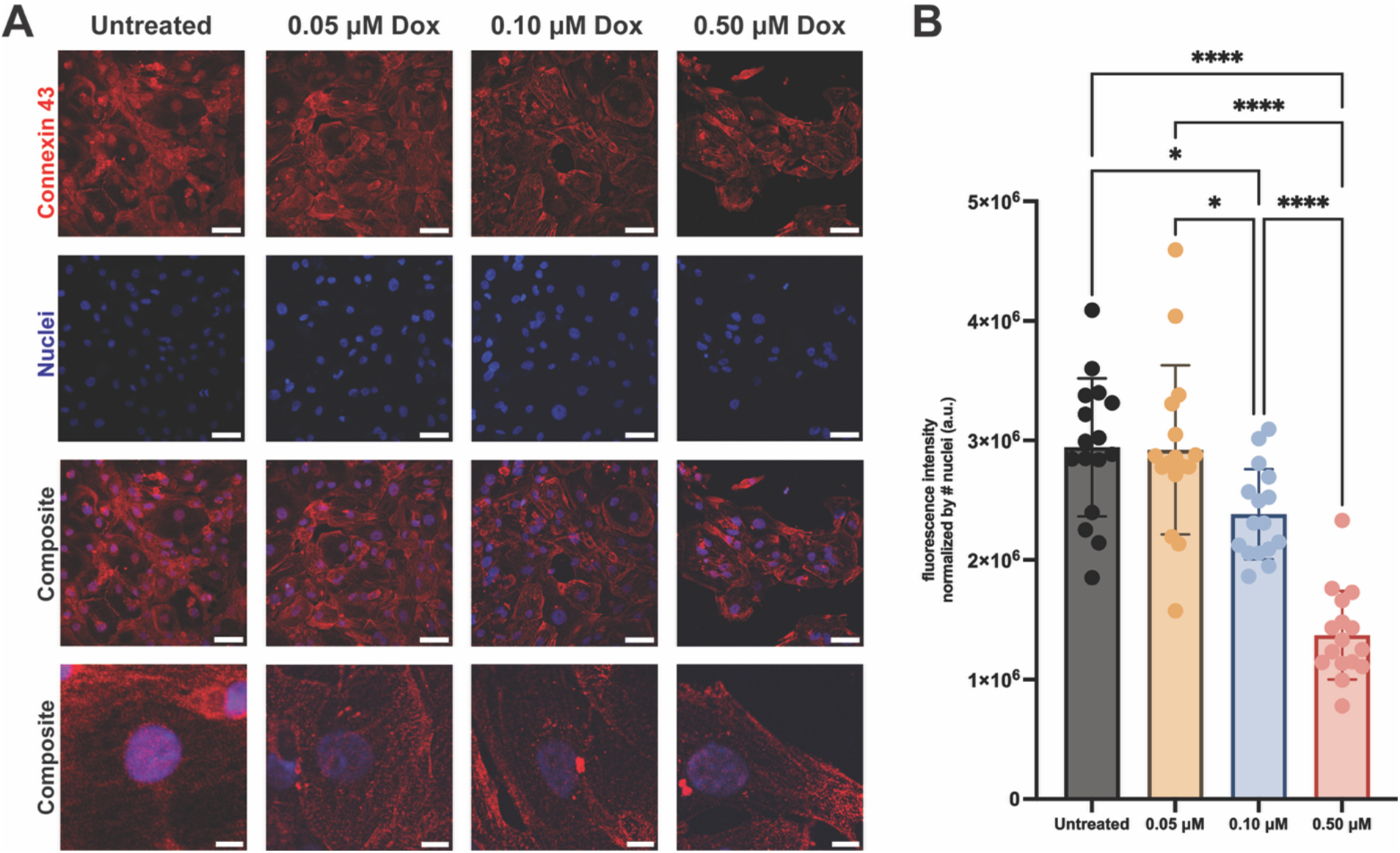
Doxorubicin Impact on Connexin43 Expression in Cardiomyocytes. **(A)** Representative images of immunofluorescence detection of connexin 43 in untreated and doxorubicin treated (0.05 µM, 0.10 µM, 0.50 µM) iPSC-CMs. **(B)** Quantification of the fluorescence intensity of connexin 43 normalized per image by the nuclei count. 4 samples were imaged per condition, with 4 fields of view analyzed per sample. Statistical significance was determined with a one-way ANOVA with Tukey’s multiple comparison, ns indicates non-significance, * indicates statistical significance with a p-value < 0.05, **** indicates statistical significance with a p-value < 0.0001.

Another major contributor to doxorubicin-induced cardiotoxicity is the disruption of proteostasis following DNA damage (Supplemental Figure 31).[9] Accumulated DNA damage impairs the transcription of proteins involved in quality control pathways, leading to a buildup of misfolded or dysfunctional proteins that further compromise cardiomyocyte function. [66] Anthracyclines have been found to cause sarcomere disruption, specifically loss of myofibrils, disorganization of the sarcomeric proteins, and insufficient production of new proteins to replace damaged proteins.[66, 67] Sarcomere proteins alpha-actinin and f-actin were immunofluorescently labeled to assess protein production and organization (Figure 9) (See Materials and Methods for details). Doxorubicin exposure caused a significant mean fold change decrease in alpha-actinin expression for each concentration, whereby there was a ca. 4.35-, 4.93-, and 2.54-fold decrease for 0.05 µM, 0.10 µM, 0.50 µM, respectively (Figure 9B). To a lesser extent, doxorubicin exposure still caused a mean fold change decrease in the expression of f-actin, whereby a ca. 0.59-, 0.54-, and 0.55-fold decrease was quantified for 0.05 µM, 0.10 µM, 0.50 µM, respectively (Figure 9C). The decrease of these structural proteins within the sarcomere indicates a reduction in the replacement of these proteins within the sarcomere.[66] Together, these assays establish what doxorubicin does to cardiomyocyte structure; chronic electrophysiology establishes when, and how gradually, it happens.

**Figure 9:**
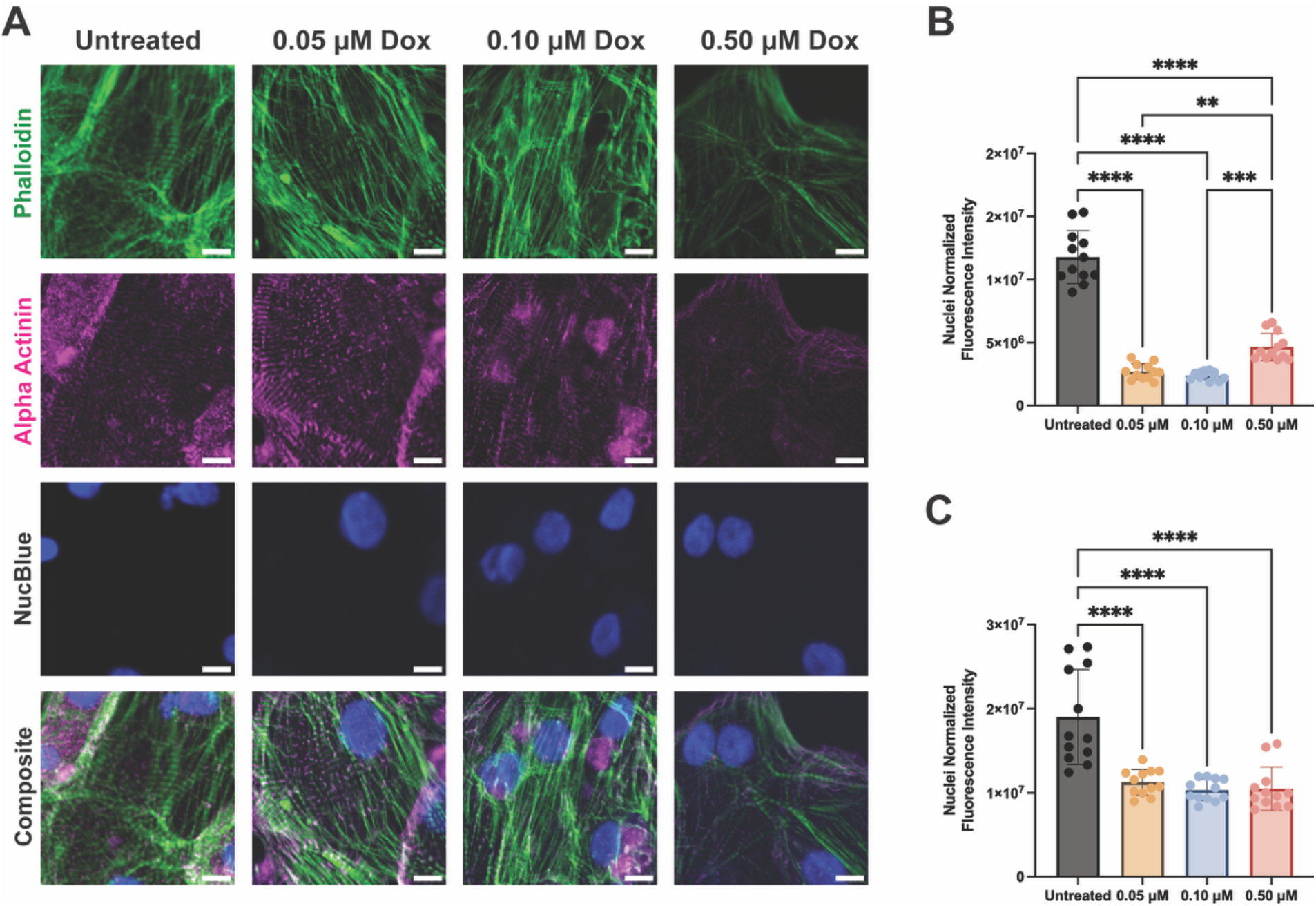
Doxorubicin Impact on Cardiomyocyte Structural Proteins. **(A)** Representative images of immunofluorescence detection of alpha-actinin and phalloidin in untreated and doxorubicin treated (0.05 µM, 0.10 µM, 0.50 µM) iPSC-CMs. **(B)** Quantification of the fluorescence intensity of alpha-actinin normalized per image by the nuclei count. 4 samples were imaged per condition, with 4 fields of view analyzed per sample. **(C)** F-actin fluorescence intensity quantification (phalloidin staining) normalized by nuclei count. 4 samples were imaged per condition, with 4 fields of view analyzed per sample. Statistical significance was determined with a one-way ANOVA with Tukey’s multiple comparison, ns indicates non-significance, ** indicates statistical significance with a p-value < 0.01, *** indicates statistical significance with a p-value < 0.001, **** indicates statistical significance with a p-value < 0.0001.

### Conclusions and Future Directions

This study highlights the knowledge that can come from continuous extracellular field potential recording spanning over 40 days in iPSC-CM monolayers, establishing a benchmark for longitudinal electrophysiological assessment *in vitro*. The spatial and temporal resolution of this study enables both single cell and monolayer level analysis. Ionic transients (Na^+^, Ca^2+^, K^+^) were assessed via single-electrode metrics (FPA, FPD) to demonstrate baseline cell-to-cell variability in healthy samples and increased variability with the development of doxorubicin induced cardiotoxicity. Doxorubicin induced progressive significant reductions to field potential amplitude and conduction velocity, upstroke and downstroke velocity, which became statistically significant over extended time scales. These changes were accompanied by significant alterations in field potential duration, beat frequency, and beat rate variability, consistent with changes in ion channel expression and permeability. Wholescale evaluation via conduction velocity analysis shows detailed changes to cell-cell communication and network behavior. The progressive electrophysiological deterioration observed following doxorubicin administration consistent with accumulation of unrepaired DNA double-strand breaks and sustained oxidative stress, reinforcing the utility of chronic monitoring for capturing the functional consequences of underlying molecular damage as it develops.

To complement the electrophysiological findings, calcium handling, mitochondrial dynamics, and reactive oxygen species were assessed throughout disease progression, revealing how underlying metabolic alterations drive the observed electrical changes, while gap junction, alpha-actinin, and f-actin expression were assessed to understand protein regulation changes during exposure. These endpoint fluorescence assays underscore the advantage of continuous monitoring afforded by chronic electrophysiology over time temporal limitations of acute, snapshot measurements.

Chronic electrophysiological monitoring demonstrates immense value for capturing the evolving electrical behavior of cardiac tissues over extended time. Incorporating chronic electrophysiology into disease modeling would clarify how electrically active tissues behave during both healthy function and disease development and could reveal the presence of cardiac injury before it becomes irreversible. Combined with patient-derived iPSC-CMs, this approach also offers a compelling avenue for personalized risk stratification in chemotherapeutic regimens. Advances in data processing and analysis methods would enable higher temporal resolution, while live data analysis could reveal nuanced pathological electrophysiological phenotypes in real-time. Further development of electrophysiology recording platforms for three-dimensional engineered heart tissues paired with chronic electrical monitoring will enable deeper insight into complex electrical dynamics for physiologically relevant cardiac systems.

## MATERIALS AND METHODS

### Device Fabrication

The microelectrode array design is shown in Figure S1. The array consists of 64, square 50 µm × 50µm electrodes in an 8×8 array with 250 µm center to center spacing between electrodes. The MEAs were microfabricated in a cleanroom facility. A 4” fused silica wafer (University Wafer, catalog no. 518) was cleaned with acetone in an ultrasonic bath for 10 minutes, rinsed with isopropyl alcohol, and N_2_ blow-dried. The wafer is coated with a layer of Hexamethyldisilazane (HMDS) to promote adhesion. A 300 nm lift-off resist layer (LOR3A, MicroChem) was then spun onto the wafer at 4000 rpm for 40 seconds and soft baked at 200°C for 5 minutes. A 1300 nm positive photoresist (Shipley 1813, MicroChem) was then spun onto the wafer at 4000 rpm for 40 seconds and soft baked for 90 seconds at 115°C. Electrodes, interconnects, and bond pads were then defined via ultraviolet (UV) exposure using a Heidelberg MLA 150 at a dose of 180mJ/cm^2^, followed by development for 20 seconds in CD-26 developer (MicroChem) and rinsing in deionized water, and the wafer is N_2_ blow-dried. A 1 minute barrel etching step (IPC Barrel Etcher) was carried out at 50W prior to metal deposition. A metal stack of 20 nm chromium and 80 nm gold was deposited using an electron beam evaporator (Kurt J. Lesker PRO Line PVD 75 E-beam Evaporator). The photoresist double-layer and excess metallic layer was then removed using a warmed 70°C bath of Remover PG (MicroChem) to define the electrodes, interconnects, and bond pads. The wafer was washed in acetone, IPA, and N_2_ blow-dried. A passivation layer of 2 μm SU-8 was defined over the metal interconnects (SU-8 2002, Kayaku Advanced Materials). The SU-8 2002 photoresist layer was spun onto the wafer at 3000 rpm for 30 seconds and soft baked for 5 minutes at 65°C and 5 minutes at 95°C. The passivation layer was defined using a Heidelberg MLA 150 with an exposure dose of 1800 mJ/cm^2^. Two post-exposure bakes were done, the first for 5 minutes at 65°C and the second for 1 minute at 95°C before development for 2 minutes in SU-8 Developer (Kayaku Advanced Materials). The wafer was hard baked at 200°C for 30 minutes and then allowed to cool down to room temperature. The wafers were then diced to separate the wafer into individual device chips.

### Device Packaging

The chips were cleaned in acetone, IPA, and N_2_ blow-dried. Screen printing of silver epoxy adhesive (MG Chemicals, catalog no. 8331S) was done to bond the devices to a custom flat-flexible cable (FFC) (Custom design on Autodesk Eagle, Fabricated by PCBWay Inc.). The silver epoxy was cured for 2 hours at 65°C. The chips were further adhered to the FFC using medical grade epoxy and left to cure at 60°C for at least 2 hours (Clear Epoxy, Loctite® M-31 CL, part no. 30673). A 10 mm Mattek dish (Mattek, P35G-1.5-10-C) with the glass bottom removed was then adhered on top of the chip using KwikSil (World Precision Instruments, KWIK-SIL-S).

### Device Characterization

Cyclic voltammetry (CV) experiments were performed in a three-electrode cell setup using a potentiostat (Reference 620, Gamry Instruments or a Palmsens4, Palmsens). Electrolyte solution of 1 M KCl (≥99%, Sigma-Aldrich, catalog no. P5405) was prepared in deionized water. Analyte solution of 5 mM K_3_[Fe(CN)_6_] (Sigma-Aldrich, catalog no. 244023) was prepared in 1 M KCl solution. Platinum wire (CH Instruments, part no. CHI115) and Ag/AgCl electrodes (CH Instruments, part no. CHI111) were used for counter and reference electrodes, respectively. CV measurements were assessed within a potential range from −0.2 to +0.2 V versus Ag/AgCl at 200 mV/s scan rate in the presence of 5 mM K_3_[Fe(CN)_6_] in 1 M KCl solution. Charge storage capacity was determined from integrating the cyclic voltammetry data and dividing by the scan rate and electrode area. Electrochemical impedance spectroscopy was performed in a three-electrode cell setup using phosphate-buffered saline (PBS) (1×) (ThermoFisher Scientific, catalog no. 10010023) as an electrolyte solution. Pt wire and Ag/AgCl electrodes were used for counter and reference electrodes, respectively. The frequency was scanned from 1 to 100,000 Hz with *V*_DC_ of 0 V. Impedance at 1kHz was determined from the 1kHz value in the full impedance spectrum.

### Device Modification: Platinum Black Deposition

Electrodeposition was performed in a three-electrode cell setup using a potentiostat. Pt wire, Ag/AgCl electrode, and Au microelectrodes were used as counter, reference, and working electrodes, respectively. The analyte solution of 25 mM Chloroplatinic acid hydrate (H_2_PtCl_6_) (Sigma Aldrich, catalog no. 520896) prepared in DI water. Platinum black deposition was performed on all 64 channels at once via chronoamperometry at -0.3V for 60 seconds. Devices were washed three times in deionized water for 5 minutes per wash.

### Device Modification: PEDOT:PSS Deposition

Electrodeposition was performed in a three-electrode cell setup using a potentiostat. Pt wire, Ag/AgCl electrode, and Au microelectrodes were used as counter, reference, and working electrodes, respectively. A solution of 0.01 M EDOT (97%, Sigma-Aldrich, catalog no. 483028) and 0.02 M PSS (Sigma-Aldrich, catalog no. 243051) was prepared in DI water. PEDOT:PSS electrodeposition was performed by using a constant current density of 0.5 mA/cm^2^ applied for 10 min per electrode.

### Electrophysiology Recordings Cell Culture

iCell® Cardiomyocytes^2^ (Fujifilm Cellular Dynamics, catalog no. 01434) were used for this study. These cells were thawed and cultured according to the User’s Guide for iCell® Cardiomyocytes^2^. Prior to seeding cells for electrophysiological recordings, devices were treated with UV/Ozone for 10 minutes at 70°C to activate the surface of the glass. Distilled water was immediately drop-casted over the surface to maintain the activation. Devices were moved into a biological safety cabinet and UV sterilized for 45 minutes. The devices were washed with ethanol and then rinsed 4 times with 1x PBS prior to coating the surface with Matrigel (Corning® Matrigel® Growth Factor Reduced (GFR) Basement Membrane Matrix, Phenol Red-free, LDEV-free, 10 mL, product no. 356231) at a dilution of 1:400 to 1:800 to promote adherence of cardiomyocytes onto the device. The Matrigel coating was incubated for 30 minutes prior to removing the coating and seeding cardiomyocytes. Removal of the excess Matrigel solution was done immediately before seeding. iCell Cardiomyocytes^2^ were thawed according to the iCell Cardiomyocytes^2^ User Manual with a seeding density of 400k cells/cm^2^ and seeded in 200 µL iCell Cardiomyocytes^2^ plating medium. 24 hours after thawing the cells, media was changed to 2 mL of iCell Cardiomyocytes^2^ maintenance medium. Media was changed and replenished on day 3 with maintenance media. For doxorubicin-administered samples, a 24-hour 0.50 µM doxorubicin (in DMSO) (Sigma-Aldrich, catalog no. D5220) treatment was administered. The doxorubicin containing media was removed and replaced with iCell Cardiomyocytes^2^ maintenance medium after 24 hours of exposure. Media was changed every 3 days until recordings were terminated. 4 Samples were recorded for both untreated and doxorubicin treated electrophysiology recordings from 3 different vials of iCell® Cardiomyocytes^2^.

### Electrophysiological Recordings Set Up

The recording devices were kept inside of a cell incubator kept at 37°C and 5% CO_2_. Four days after seeding, devices were connected to the Intan Recording system (RHD 1024-channel recording controller, Intan Technologies, catalog no. C3008). The FFC was designed to extend from inside the incubator to the outside of the incubator where it was inserted into a zero-insertion force (ZIF) socket which connected to two 32-channel amplifiers (Intan Technologies, catalog no. C3314) or two devices were connected to a 128-channel amplifier (Intan Technologies, catalog no. C3316). An external platinum wire reference electrode was used, submerged within the media throughout the course of the recording (Kurt J. Lesker, catalog no. EVMPT40020). The electrical signals were recorded using an Intan acquisition system with a sampling rate of 20kHz. The electrical recordings were done chronically, with a custom script controlling the Intan to record data from every channel for 10 minutes each hour across the days-long recordings.

### Electrophysiological Recording Data Analysis

Representative field potential traces were filtered using a notch filter for 60Hz harmonics. Representative channels were chosen based on functional channels present across the whole recording timeframe.

#### Average Field Potential Trace

Data files are loaded into a custom MATLAB (MathWorks, Inc.) script which concatenates multiple Intan Recording files together. The Na^+^ peaks for each active channel in the data are found using the findpeaks MATLAB function based on a ‘MinPeakDistance’ depending on the beating rate of the recording, set manually. This returns the time points of these peaks and the amplitude at the peak. The Na^+^ peaks were then overlaid to align the peak of each field potential and include about 0.7 seconds of recording around each peak (this value being variable based on the recording).

#### Field Potential Amplitude

Data files are loaded into a custom MATLAB script which concatenates multiple Intan Recording files together. The Na^+^ peaks for each active channel in the data are found using the findpeaks MATLAB function based on a ‘MinPeakDistance’ depending on the beating rate of the recording, set manually. This returns the time points of these peaks and the amplitude at the peak. The values of the field potentials were then inverted across the length of the data input, and the negative peak for each positive peak found. The positive and negative peaks were aligned based on the time stamp of each peak and the field potential amplitudes were found by adding the amplitude of the positive and negative peak positions of the amplitude together.

#### Beat to Beat Variability

The beat-to-beat variability was calculated per channel using the root mean square of successive differences (RMSSD). The peaks in each channel are found using the findpeaks MATLAB function and then the difference in time between successive peaks are found. The absolute value of the difference between successive differences is then found, squared, and all of these values are averaged together across the beats for each channel. The square root is then found of the averaged value.

#### Upstroke/Downstroke Velocity

The upstroke and downstroke velocity were calculated based on the first derivative of the electrophysiology recordings using the gradient MATLAB function. For each channel the first derivative was taken, and the upstroke velocity was determined by the positive peak of the first derivative, and the downstroke velocity was determined by the negative peak of the first derivative for each peak at each channel, determined via the findpeaks MATLAB function. Values were averaged by recording channel.

#### Sodium Width

The spike duration was determined based on the time difference between the locations of the first derivative positive and negative values corresponding to upstroke and downstroke velocity.

#### Field Potential Duration

Data files are loaded into a custom MATLAB script which concatenates multiple Intan Recording files together. The data was filtered through a 1kHz low pass filter using the MATLAB functions butter and filtfilt. The Na^+^ peaks in the data were found by using the absolute value of the recording data with a minimum height of 2 times the standard deviation of the data and a minimum peak distance of the sampling frequency. The K^+^ peak for each Na^+^ peak was found by finding the maximum value within a manually defined time window after each Na^+^ peak, typically set to between .3 to .7 seconds after the Na^+^ peak. The field potential duration was calculated by subtracting the time associated with the K^+^ from the time associated with the Na^+^ peak.

#### Signal Propagation Analysis

Data files are loaded into a custom MATLAB script which concatenates multiple Intan Recording files together. The Na^+^ peaks for each active channel in the data are found using the findpeaks MATLAB function based on a ‘MinPeakDistance’ depending on the beating rate of the recording, set manually. This returns the time points of these peaks and the amplitude at the peak. The starting channel for each field potential in the recording was identified by finding the minimum timed peak for each collective beat of the cardiac monolayer. The most frequently determined starting channel was used to determine the direction of the signal across the monolayer. The time latency between the first channel and each other channel was calculated for each field potential set in the data. These times were then either averaged for each channel or used as individual values for creating videos of the signal propagation. The time latency values were assigned into a matrix which associated each electrode position with each channels time latency. This matrix was used to plot a map of values which could be interpolated using the MATLAB function scatteredInterpolant with a linear interpolation method and linear extrapolation method. The MATLAB function contourf was used to create an isochronal map with 15 levels utilizing the scatteredInterpolant results after a 2D Gaussian smoothing kernel with a sigma of 2 was used to smooth the image (MATLAB function imgaussfilt). To determine the direction of the signal propagation, the gradient of the 2D Gaussian smoothed scatteredInterpolant results were used to create a quiver plot, and the average quiver was used to determine the average direction and magnitude of the signal propagation, scaled into centimeters per second based on the map of the electrodes.

### Seeding Cardiomyocytes onto Glass and Doxorubicin Treatment

Glass dishes (Mattek, part no: P35G-1.5-7-C) with a diameter of 7mm were treated with UV/Ozone for 10 minutes at 70°C to activate the surface of the glass. Distilled water was immediately drop-casted over the surface to maintain the activation. Dishes were moved into a biological safety cabinet and UV sterilized for 45 minutes. The dishes were washed 3 times with 1x PBS prior to coating the glass with 200 µL of 0.5 µg/mL of fibronectin (Advanced BioMatrix, catalog no. 5050) or Matrigel (Corning® Matrigel® Growth Factor Reduced (GFR) Basement Membrane Matrix, Phenol Red-free, LDEV-free, 10 mL, product no. 356231) to promote adherence of cardiomyocytes on the glass surface. The fibronectin coating was incubated for 1 hour prior to removing the coating and seeding cardiomyocytes and Matrigel solution was incubated for 30 minutes prior to removal. Removal of the excess solution was done immediately before seeding. iCell Cardiomyocytes^2^ were thawed according to the iCell Cardiomyocytes^2^ User Manual with a seeding density of 300k cells/cm^2^. 24 hours after thawing the cells, 2 mL of maintenance media was added to the dish. The next day, the media was removed, and 1 mL of maintenance media was added to each dish. Media was changed on day 3 to maintenance media. Four days after seeding, the cells were treated with 0.05µM, 0.1µM, or 0.5µM of doxorubicin (Sigma-Aldrich, catalog no. D5220). Media was changed after 24 hours of doxorubicin exposure. Maintenance media was changed every two days until imaging/termination.

### Calcium Imaging

Cardiomyocytes were taken at day 5 after a 24 hour doxorubicin treatment (0 µM, 0.05µM, 0.10 µM, 0.50 µM) for calcium imaging. CalBryte^TM^ 520 AM (AAT Bioquest, catalog no. 20651) was used to stain the cells at a concentration of 5 µM in iCell Cardiomyocyte^2^ Maintenance Medium. Cells were rinsed with 1x PBS 3 times prior to Tyrode’s solution (Sigma Aldrich, catalog no: T2145-10L) being added to each dish. Cells were imaged at 40X using an upright confocal microscope (Nikon A1R) taking 40 second videos at 30 frames per second, while the temperature inside the culture chamber was maintained by constant perfusion of Tyrode’s solution maintained at 37°C using an inline heater (ThermoClamp, Automate Scientific).

### Reactive Oxygen Species Imaging – 12 hour and 24-hour Timepoints

Cardiomyocytes were taken at 12 hours and 24 hours after either no treatment or 0.50 µM doxorubicin treatment for assessing reactive oxygen species. Cells were rinsed with 1x PBS (ThermoFisher Scientific, catalog no. 10010023) and stained with 2 µM of CM-H2DCFDA (ThermoFisher Scientific, catalog no. C6827) in Tyrode’s solution (Sigma Aldrich, catalog no: T2145-10L) for 30 minutes at 37°C, 5% CO_2_. NucRed (ThermoFisher Scientific, catalog no. R37106) was used as a nuclear stain and incubated for 15 minutes. Cells were rinsed with 1x PBS 3 times prior to Tyrode’s solution being added to each dish. Cells were imaged at 40X using an upright confocal microscope (Nikon A1R), and the temperature inside the culture chamber was maintained by constant perfusion of Tyrode’s solution maintained at 37°C using an inline heater (ThermoClamp, Automate Scientific).

### Reactive Oxygen Species Imaging – 5 day Timepoint

Cardiomyocytes were taken at day 5 after doxorubicin treatment (0 µM, 0.05µM, 0.10 µM, 0.50 µM) for assessing reactive oxygen species. Cells were rinsed with 1x PBS (ThermoFisher Scientific, catalog no. 10010023) and stained with 2 µM of CM-H2DCFDA (ThermoFisher Scientific, catalog no. C6827) in Tyrode’s solution (Sigma Aldrich, catalog no: T2145-10L) for 30 minutes at 37°C, 5% CO_2_. NucRed (ThermoFisher Scientific, catalog no. R37106) was used as a nuclear stain and incubated for 15 minutes. Cells were rinsed with 1x PBS 3 times prior to Tyrode’s solution being added to each dish. Cells were imaged at 40X using an upright confocal microscope (Nikon A1R), and the temperature inside the culture chamber was maintained by constant perfusion of Tyrode’s solution maintained at 37°C using an inline heater (ThermoClamp, Automate Scientific).

### Mitochondrial Membrane Potential Imaging

Cardiomyocytes were taken at day 5 after doxorubicin treatment (0 µM, 0.05µM, 0.10 µM, 0.50 µM) for assessing mitochondrial membrane potential. Cells were stained with 100 nM of Mitotracker CXMROS (ThermoFisher Scientific, catalog no. M7512) for 40 minutes at 37°C, 5% CO_2_. The nuclei were stained with 1 µM Hoechst dye (ThermoFisher Scientific, catalog no. H3570) for 20 minutes at 37°C, 5% CO_2_. The cells were washed 3 times with 1x PBS (ThermoFisher Scientific, catalog no. 10010023). The cells were then fixed in 4% paraformaldehyde (ThermoFisher Scientific, cat. # J19943.K2) for 15 minutes. Cells were then washed 3 times with 1x DPBS (ThermoFisher Scientific, cat. #14040117). Cells were kept in 1x DPBS for imaging. Cells were imaged using an upright confocal microscope (Nikon A1R) at 40x.

### γ-H2AX Imaging

Cardiomyocytes were taken at day 5 after doxorubicin treatment (0 µM, 0.05µM, 0.10 µM, 0.50 µM) for assessing DNA-damage via *γ-H2AX Imaging*. Cells were fixed in 4% paraformaldehyde (ThermoFisher Scientific, cat. # J19943.K2) for 15 minutes. Cells were then washed 3 times with 1x DPBS (ThermoFisher Scientific, cat. #14040117). Cells were permeabilized with 0.2% Triton-X (Invitrogen™, cat. # HFH10) for 15 minutes. Cells were then washed 3 times with 1x DPBS. Blocking was done with 2% BSA (Invitrogen™, cat. # 15561020) in 1x DPBS for 45 minutes. Cells were washed with 1x DPBS 3 times. Primary staining was done at 4°C overnight in 1x DPBS with 0.5% BSA at a dilution of 1:200 using Phospho-Histone H2A.X (Ser139) Rabbit mAb (Cell Signaling Technology, cat. # 9718). After washing the cells with 2% BSA (in 1x DPS) 3 times, the secondary antibody, Anti-rabbit IgG, Alexa Fluor ® 647 Conjugate, was stained at room temperature for 60 minutes at a dilution of 1:1000 in 0.5% BSA. NucBlue (Invitrogen NucBlue™ Live ReadyProbes Reagent (Hoechst 33342)) was used as a nuclear stain. 2 drops were used per mL of 0.5% BSA in 1x PBS. Cells were washed 3 times with 2% BSA and 3 times with 1x DPBS before being stored in 1x DPBS at 4°C. Cells were imaged using an upright confocal microscope (Nikon A1R) at 40x.

### a-Actinin / Phalloidin Imaging

Cardiomyocytes were taken at day 5 after doxorubicin treatment (0 µM, 0.05µM, 0.10 µM, 0.50 µM) for assessing alpha-actinin and phalloidin expression and organization. Cells were fixed in 4% paraformaldehyde (ThermoFisher Scientific, cat. # J19943.K2) for 15 minutes. Cells were then washed 3 times with 1x DPBS (ThermoFisher Scientific, cat. #14040117). Cells were permeabilized with 0.1% Triton-X (Invitrogen™, cat. # HFH10) for 5 minutes. Cells were then washed 3 times with 1x DPBS. Blocking was done with 2% BSA (Invitrogen™, cat. # 15561020) in 1x DPBS for 45 minutes. Cells were washed with 1x DPBS 3 times. Primary staining was done at 4°C overnight in 1x DPBS with 0.5% BSA at a dilution of 1:100 using alpha-Actinin (D6F6) XP® Rabbit mAb (Cell Signaling Technology, cat. # 6487). After washing the cells with 2% BSA (in 1x DPS) 3 times, the secondary antibody, Anti-rabbit IgG, Alexa Fluor ® 647 Conjugate, was stained at room temperature for 60 minutes at a dilution of 1:1000 in 0.5% BSA. Phalloidin was stained using 1X Alexa Fluor ® 488 Phalloidin (Cell Signaling Technology, cat. # 8878) for 15 minutes at room temperature in 0.5% BSA in 1x PBS. NucBlue (Invitrogen NucBlue™ Live ReadyProbes Reagent (Hoechst 33342)) was used as a nuclear stain. 2 drops were used per mL of 0.5% BSA in 1x PBS. Cells were washed 3 times with 2% BSA and 3 times with 1x DPBS before being stored in 1x DPBS at 4°C. Cells were imaged using an upright confocal microscope (Nikon A1R) at 40x.

### Cx43 Imaging

Cardiomyocytes were taken at day 5 after doxorubicin treatment (0 µM, 0.05µM, 0.10 µM, 0.50 µM) for assessing gap junction expression via connexin 43 presence. Cells were fixed in 4% paraformaldehyde (ThermoFisher Scientific, cat. # J19943.K2) for 15 minutes. Cells were then washed 3 times with 1x DPBS (ThermoFisher Scientific, cat. #14040117). Cells were permeabilized with 0.1% Triton-X (Invitrogen™, cat. # HFH10) for 5 minutes. Cells were then washed 3 times with 1x DPBS. Blocking was done with 2% BSA (Invitrogen™, cat. # 15561020) in 1x DPBS for 45 minutes. Cells were washed with 1x DPBS 3 times. Primary staining was done at 4°C overnight in 1x DPBS with 0.5% BSA at a dilution of 1:100 using Connexin 43 mAb (Thermofisher #14-4759-82). After washing the cells with 2% BSA (in 1x DPS) 3 times, the secondary antibody, Anti-mouse IgG, Alexa Fluor ® 555 Conjugate (Cell Signaling #4409), was stained at room temperature for 60 minutes at a dilution of 1:1000 in 0.5% BSA. NucBlue (Invitrogen NucBlue™ Live ReadyProbes Reagent (Hoechst 33342)) was used as a nuclear stain. 2 drops were used per mL of 0.5% BSA in 1x PBS. Cells were washed 3 times with 2% BSA and 3 times with 1x DPBS before being stored in 1x DPBS at 4°C. Cells were imaged using an upright confocal microscope (Nikon A1R) at 40x.

### Calcium Imaging Analysis

Videos of the calcium transient are taken and the fluorescence intensity of the video over time is measured using Fiji ImageJ (National Institutes of Health) for 4 separate videos/ROIs per sample. Average fluorescence calcium transients were calculated by thresholding each video based on the max intensity projection of each video. Each video was divided into a 5×5 grid of areas, and each area was used for analysis based on the transient at each peak. The metrics from the calcium imaging are all found from baseline subtracted traces. The peak amplitude was determined using the findpeaks MATLAB function to locate and determine the fluorescence intensity of each peak. The normalized fluorescence trace was found by normalizing baseline to 0 and the peak maximum to 1. The rise time was determined by averaging the normalized calcium transients for each video and determining the time at which the increase in fluorescence reached 10% and 90% of the peak maximum. The rise time is then taken from the difference between these two values. The 80% peak width is found from the normalized averaged peak for each area of each video taken. The point during the influx and decay at which the fluorescence intensity is 20% of the peak amplitude is found and then the width is then determined from subtracting the decay crossing at 20% from the influx crossing the 20% threshold. The exponential fit of the peak decay is found from the normalized averaged peak for each area of each video taken. The exponential fit is done on the second half of the decay using the nlinfit function in MATLAB. The exponential decay constant, tau, is determined based on the fit for the second half of the decay. Data was compared between untreated and 0.5 µM and boxplots were used to display the mean as the center of the boxplot and whiskers showing the standard deviation of the data with outliers marked in red “+” and were determined by values outside of the 1.5 × IQR of the data.

### Fluorescence Intensity Image Analysis

The fluorescence intensity of each image was determined using a custom MATLAB code which calculates the intensity of each image and subtracts the background intensity and removes any debris in the image. The number of nuclei in each image was also measured using a custom MATLAB code which thresholds each nuclei image and counts the nuclei per image. The fluorescence intensity is then averaged per image by the number of nuclei in the image or by the number of pixels of nuclei for the γH2AX imaging set.

### Statistical Analysis

Statistical analysis was done in GraphPad^®^. Statistical significances were determined with either student t-tests, one-way ANOVA with Dunnett’s multiple comparison test or with a one-way ANOVA with Tukey’s multiple comparison depending on the experiment. Individual experiments have the specific analysis noted. In graphs, ns indicates non-significance, * indicates statistical significance with a p-value < 0.05, ** indicates statistical significance with a p-value < 0.01, *** indicates statistical significance with a p-value < 0.001, **** indicates statistical significance with a p-value < 0.0001.

## Supplementary Information

**Figure S1:**
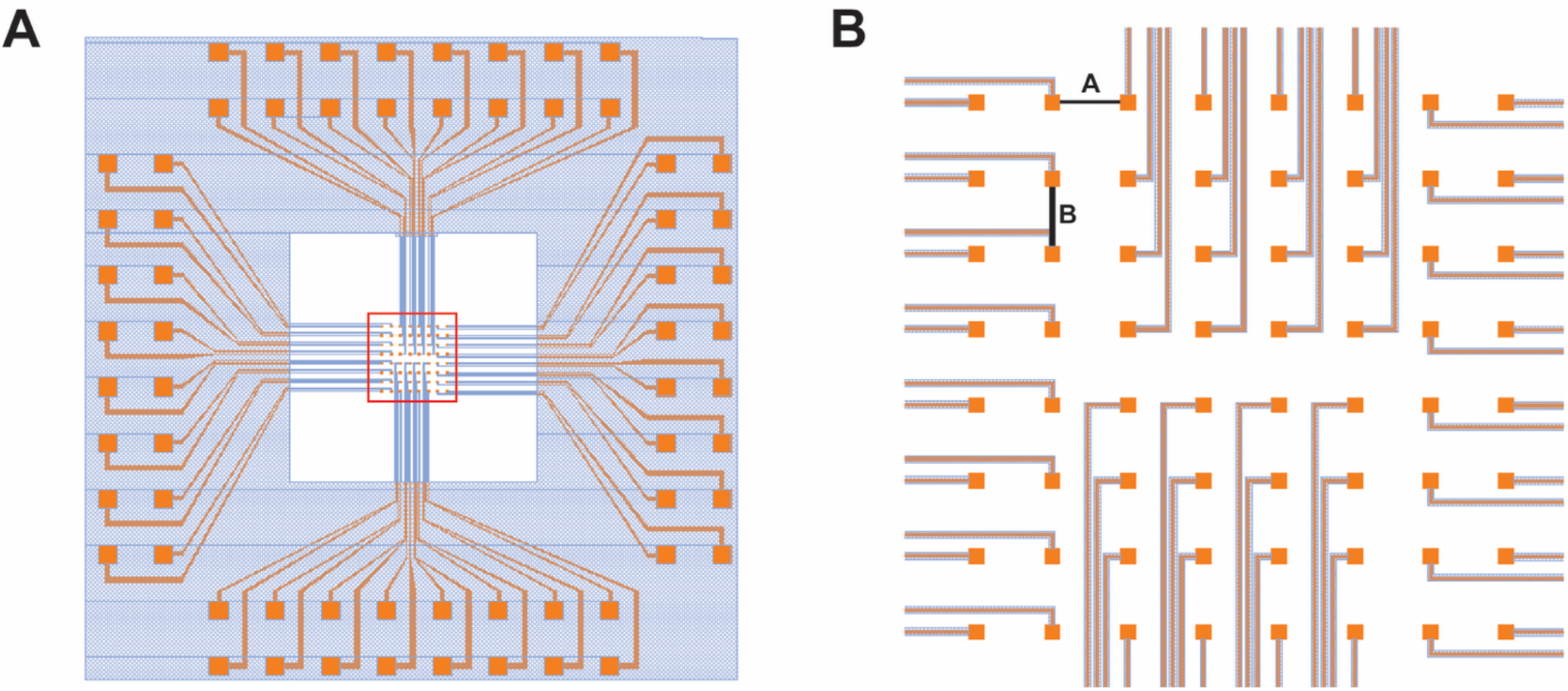
KLayout Microelectrode Array Design. **(A)** KLayout Schematic of 64 channel microelectrode arrays. Gold areas represent the metal lines, bond pads, and electrodes which when fabricated layer consists of 20nm Cr and 80nm Au. Blue areas represent the insulation layer, when fabricated this layer consists of 2 microns of SU-8 polymer. **(B)** KLayout schematic of 8×8 array of electrodes. Both distances A and B are 200 microns.

**Figure S2:**
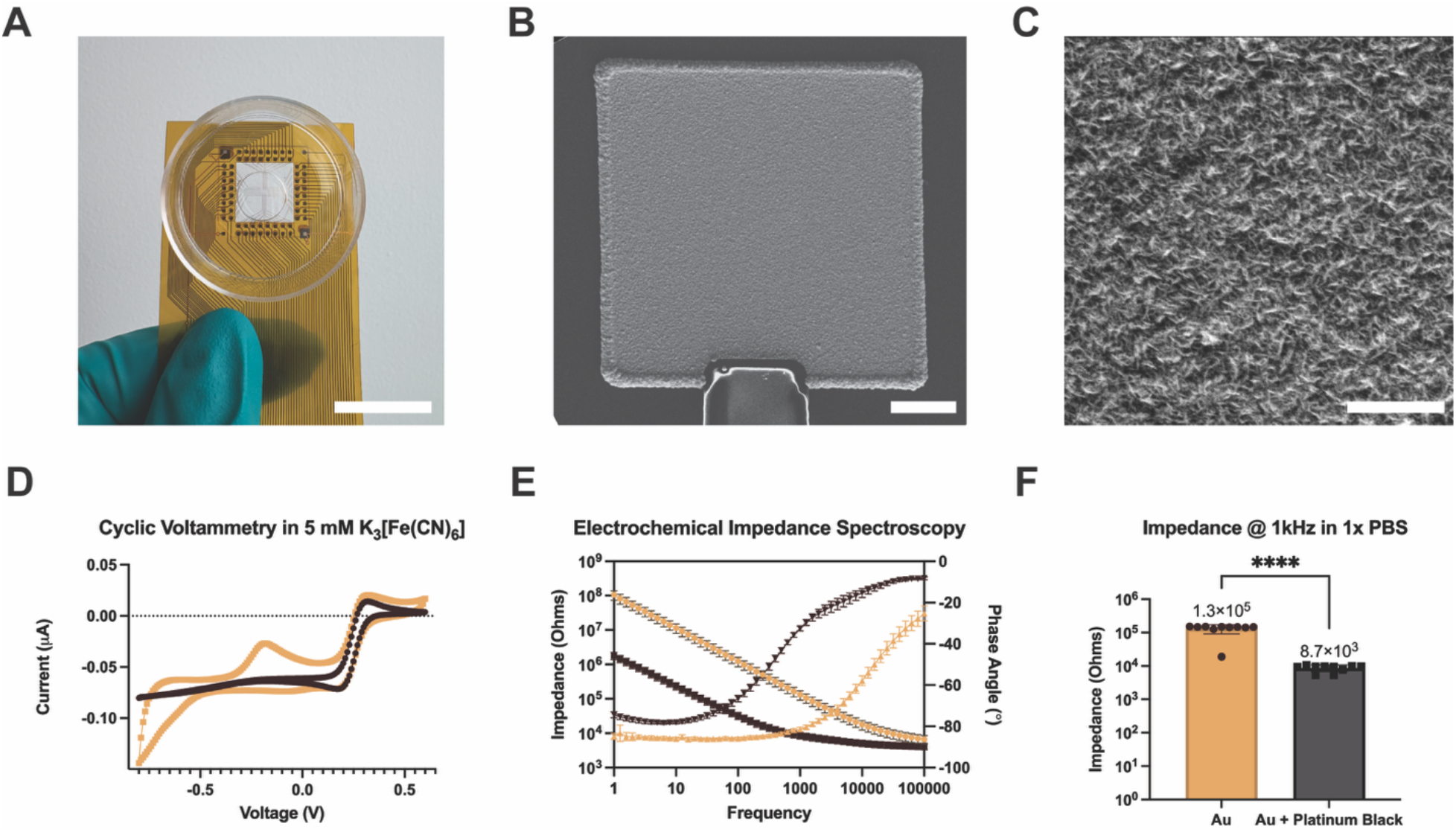
Two-dimensional Microelectrode Array Characterization. **(A)** Image of packaged microelectrode array (MEA), with 3D printed well, and flat flexible cable. Scale bar = 20 mm. **(B)** Scanning electron microscopy image of platinum black on Au electrode. Scale bar = 10 microns. **(C)** Scanning electron microscopy image of platinum black on Au electrode. Scale bar = 1 micron. **(D)** Cyclic voltammetry of 50 × 50 micron Au electrodes before and after modification with Platinum Black carried out with commercial reference electrode Ag/AgCl and a Pt wire reference electrode, n = 10 per condition. **(E)** Electrochemical impedance spectroscopy showing impedance magnitude and phase angle of 50 × 50 micron Au electrodes before and after modification with platinum black in 1x PBS carried out with commercial reference electrode Ag/AgCl and a Pt wire reference electrode, n= 10 per condition. **(F)** Electrochemical impedance spectroscopy showing impedance at 1kHz of 50 × 50 micron Au electrodes before and after modification with platinum black in 1x PBS carried out with commercial reference electrode Ag/AgCl and a Pt wire reference electrode, n = 10 per condition. Statistical significance was determined with a student t-test, **** indicates statistical significance with a p-value < 0.0001.

**Figure S3:**
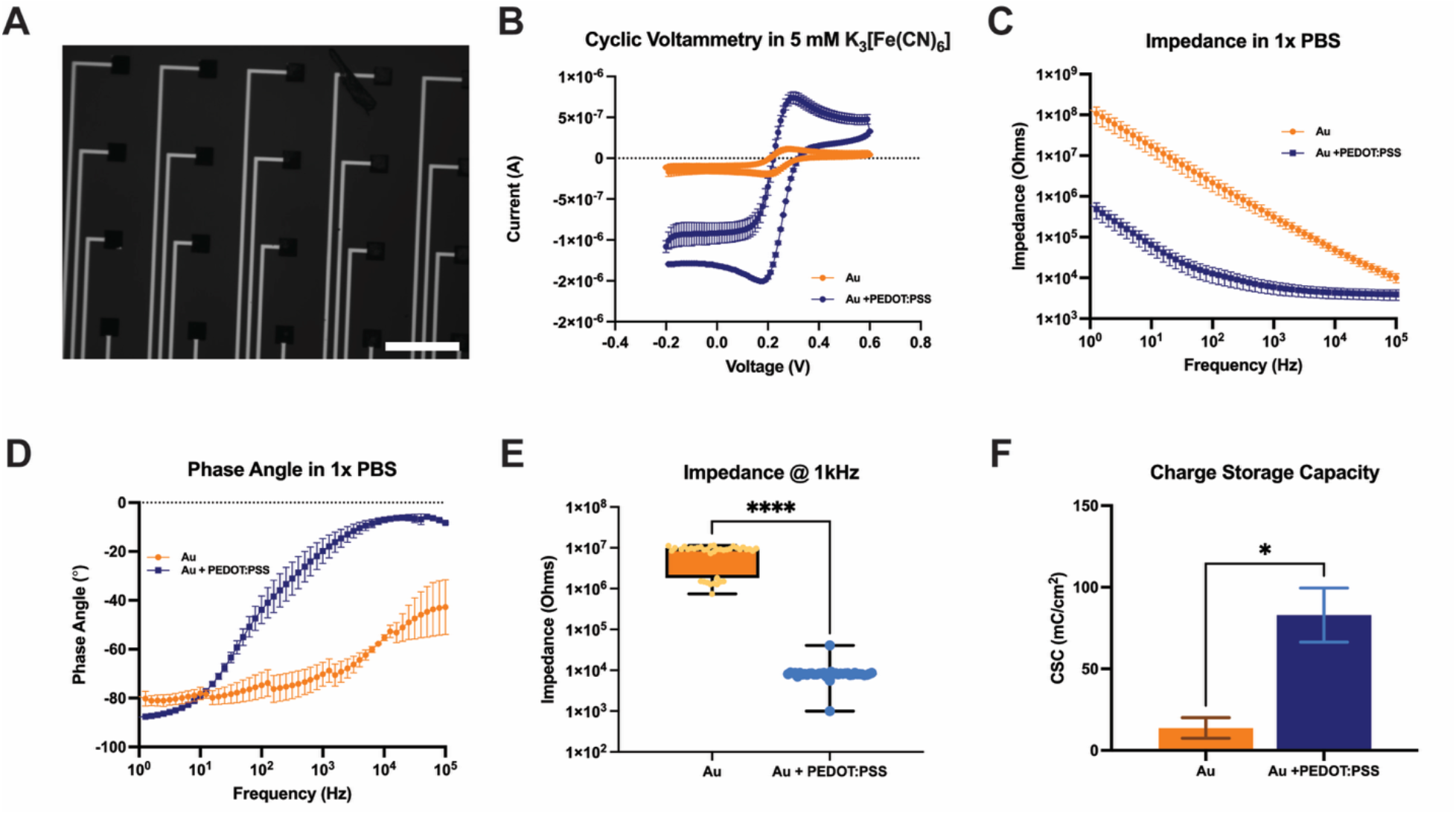
Two-Dimensional Microelectrode Array Characterization after PEDOT:PSS Deposition. **(A)** Brightfield image of 50 × 50 micron Au electrodes modified with poly(3,4-ethylenedioxythiophene): poly(sodium 4-styrenesulfonate) (PEDOT:PSS) by electrodeposition. Scale bar = 200 microns. **(B)** Cyclic voltammetry of 50 × 50 micron Au electrodes before and after modification with PEDOT:PSS in 5 mM K_3_[Fe(CN)_6_] carried out with commercial reference electrode Ag/AgCl and a Pt wire reference electrode, n= 3 per condition. **(C)** Electrochemical impedance spectroscopy showing impedance magnitude of 50 × 50 micron Au electrodes before and after modification with PEDOT:PSS in 1x PBS out with commercial reference electrode Ag/AgCl and a Pt wire reference electrode, n= 3 per condition. **(D)** Phase Angle of 50 × 50 micron Au electrodes before and after modification with PEDOT:PSS in 1x PBS carried out with commercial reference electrode Ag/AgCl and a Pt wire reference electrode, n= 3 per condition. **(E)** Electrochemical impedance spectroscopy showing impedance at 1kHz of 50 × 50 micron Au electrodes before and after modification with PEDOT:PSS in 1x PBS carried out with commercial reference electrode Ag/AgCl and a Pt wire reference electrode, n= 40 per condition. Statistical significance was determined with a student t-test, **** indicates statistical significance with a p-value < 0.0001. **(F)** Charge storage capacity calculated from cyclic voltammetry of 50 × 50 micron Au electrodes before and after modification with PEDOT:PSS in 1x PBS carried out with commercial reference electrode Ag/AgCl and a Pt wire reference electrode, n= 3 per condition. Statistical significance was determined with a student t-test, * indicates statistical significance with a p-value < 0.05.

**Figure S4:**
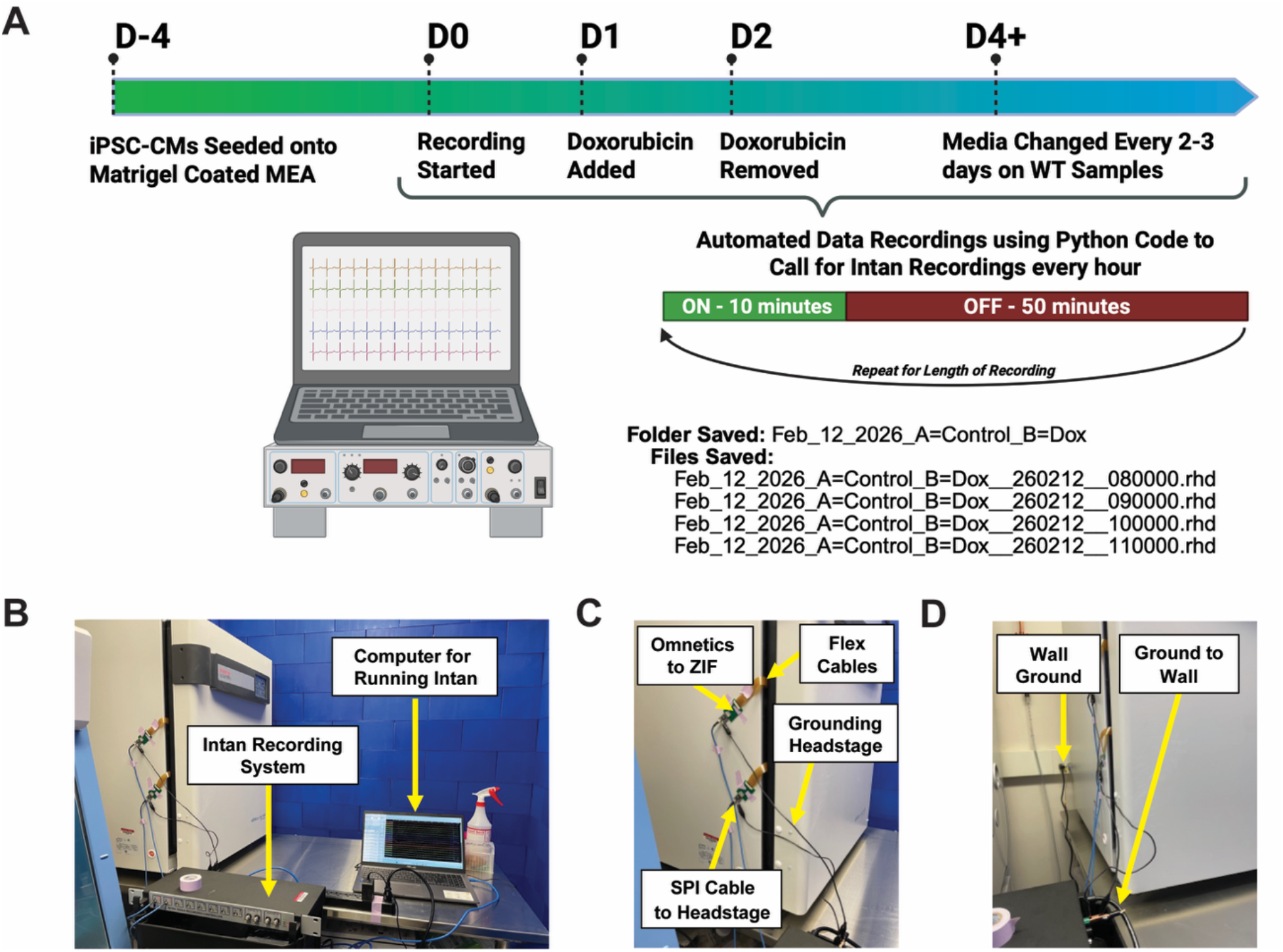
Chronic Electrophysiological Set Up. **(A)** Experimental and recording timeline. **(B)** Image of Intan Recording System set up outside of incubator. **(C)** Image of device flat flexible cable exiting the computer, connecting to Omnetics of Intan Recording System. **(D)** Image of grounding set up used for chronic recordings.

**Figure S5:**
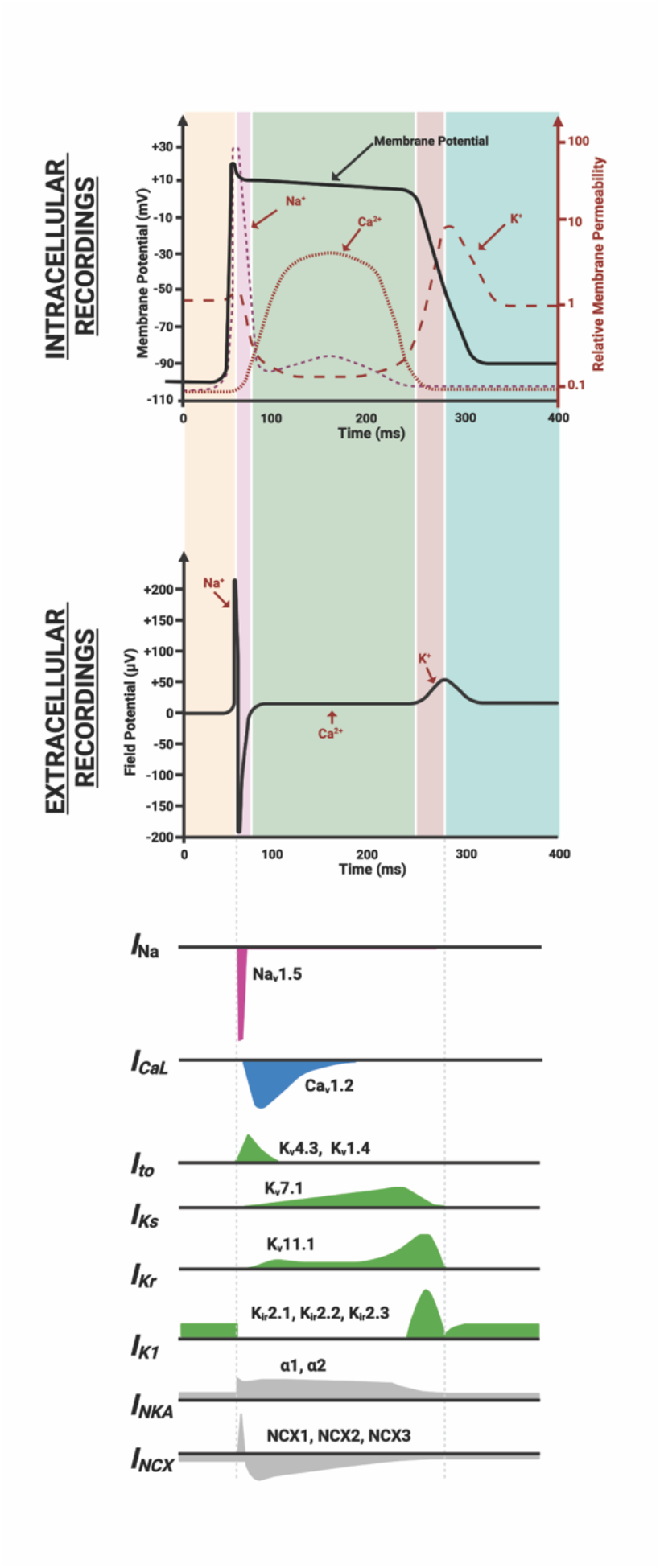
**Cardiac Action Potential to Cardiac Extracellular Field Potential Comparison.**

**Figure S6:**
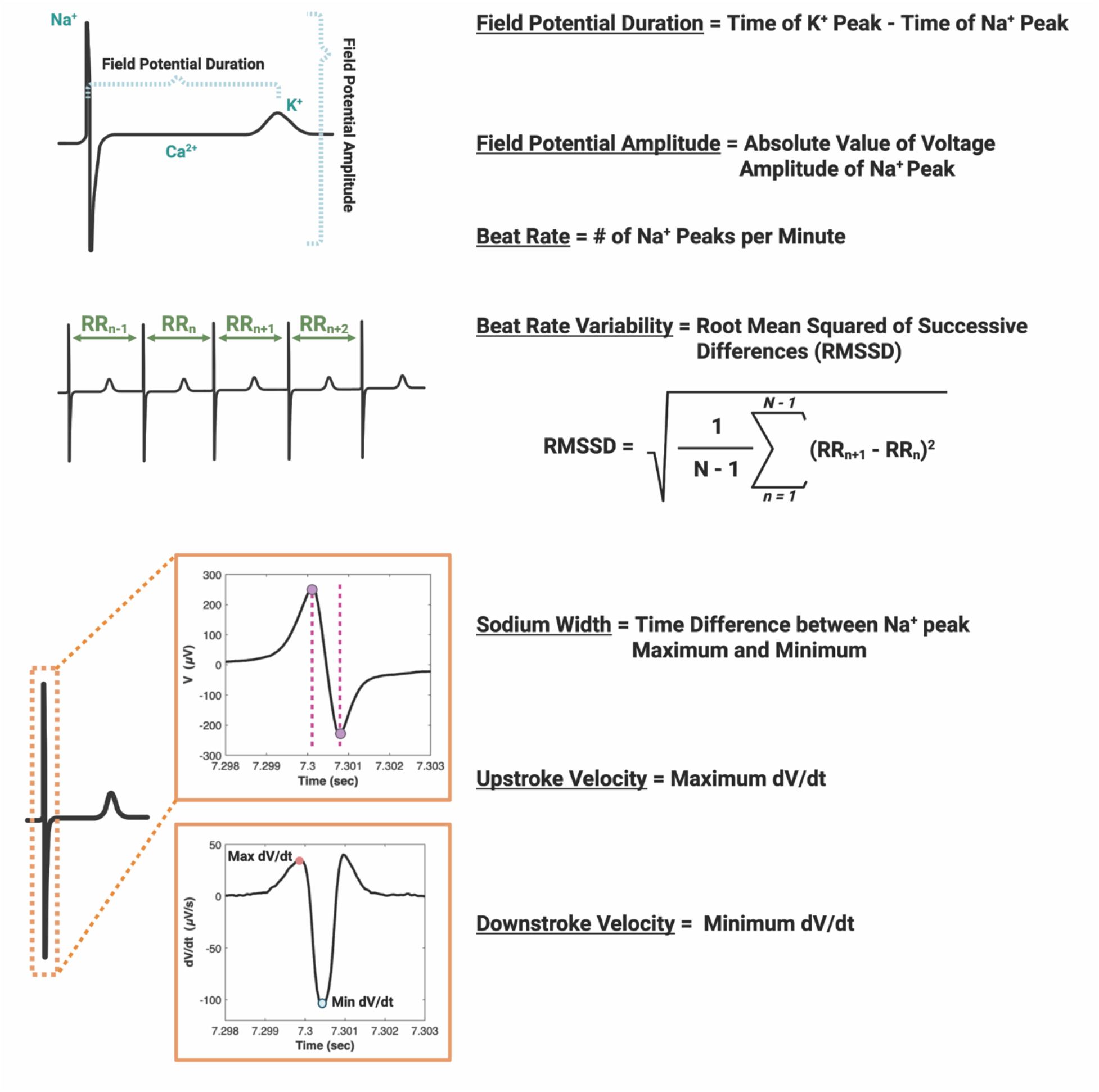
**Electrophysiology Metrics Calculated for Cardiac Extracellular Field Potentials.**

**Figure S7:**
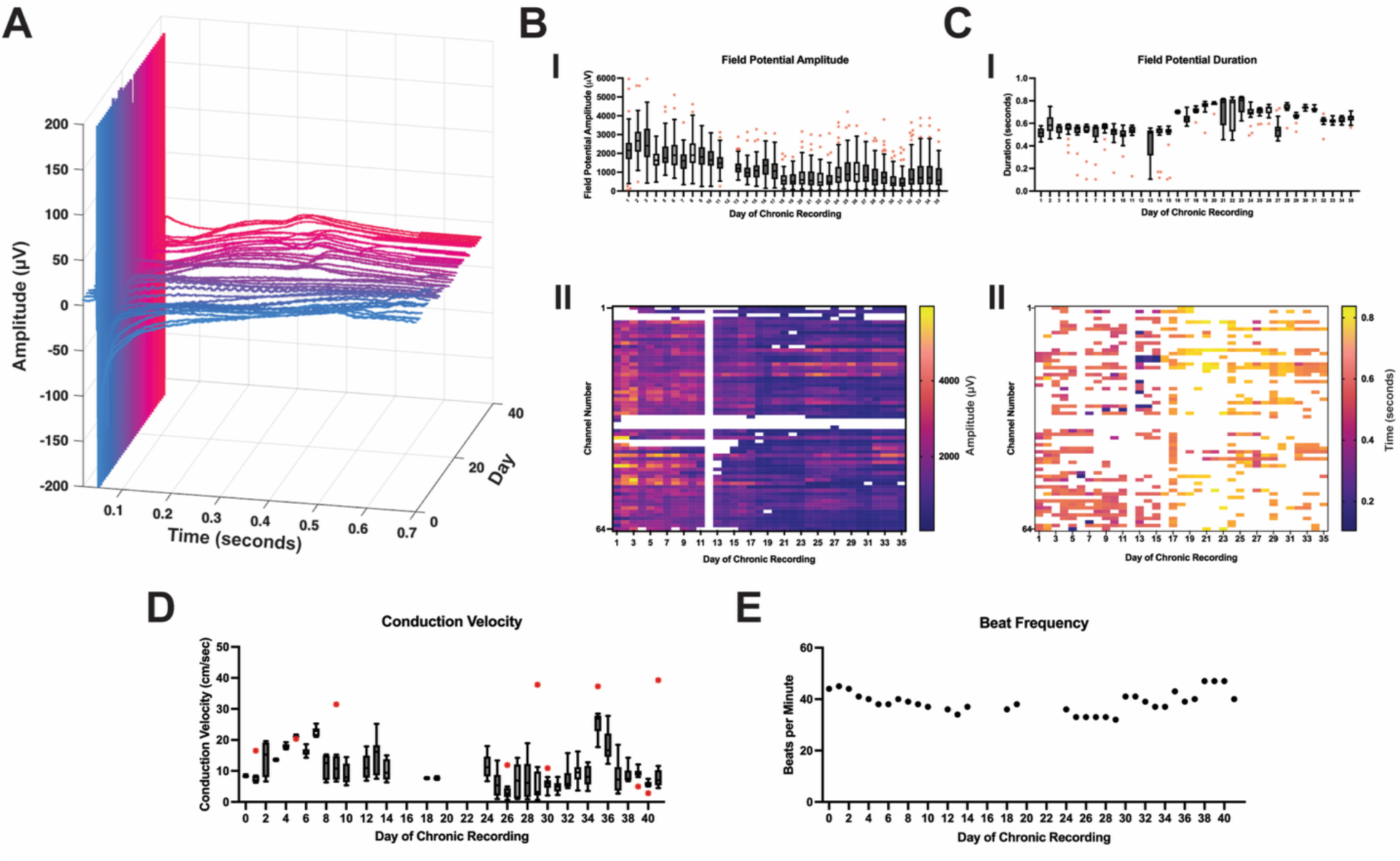
40 Day Chronic Recording from untreated iPSC-CM monolayer. **(A)** Representative extracellular field potential from a single channel over 40 days of chronic recordings. **(B)** Field Potential Amplitude (I) Boxplot of field potential amplitude across 40 days of chronic recording, with whiskers extending to 1.5× IQR and outliers shown in red. (II) Heatmap shows individual electrode amplitude values for each day of recording, white areas indicate field potential recorded. **(C)** Field potential duration, (I) Boxplot of field potential amplitude across 40 days of chronic recording, with whiskers extending to 1.5× IQR and outliers shown in red. (II) Heatmap shows individual electrode duration values for each day of recording, white areas indicate potassium peak in recorded data. **(D)** Conduction velocity over 40 days of chronic recording. **(E)** Beat frequency over 40 days of chronic recording.

**Figure S8:**
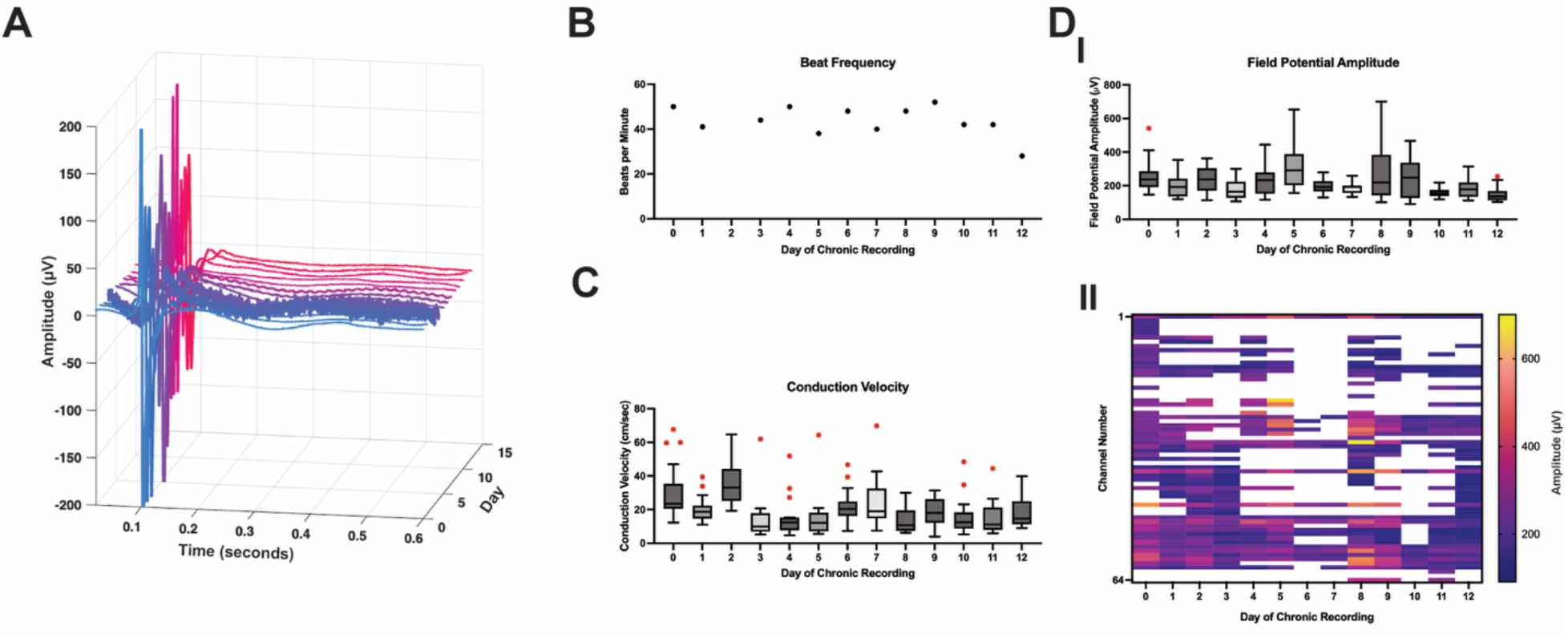
Additional Sample untreated iPSC-CM Chronic Recording. **(A)** Representative extracellular field potential from a single channel over 12 days of chronic recordings. **(B)** Beat frequency over 12 days of chronic recording. **(C)** Conduction velocity across the cardiac monolayer over 12 days of chronic recording. **(D)** Field Potential Amplitude, (I) Boxplot of field potential amplitude across 12 days of chronic recording, with whiskers extending to 1.5× IQR and outliers shown in red. (II) Heatmap shows individual electrode amplitude values for each day of recording, white areas indicate field potential recorded.

**Figure S9:**
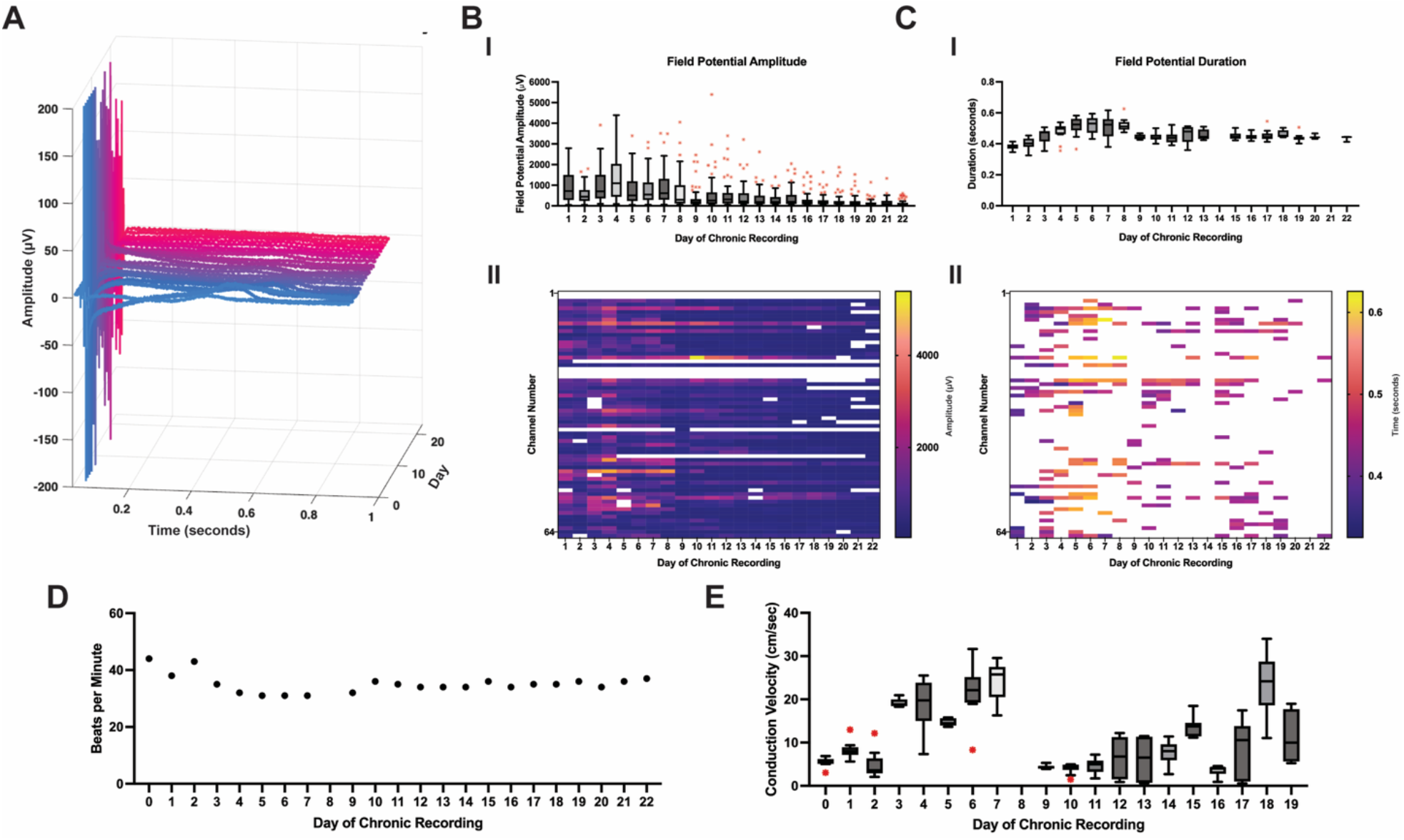
Additional Chronic Recording from untreated iPSC-CM monolayer. **(A)** Representative extracellular field potential from a single channel over 22 days of chronic recordings. **(B)** Field Potential Amplitude (I) Boxplot of field potential amplitude across 22 days of chronic recording, with whiskers extending to 1.5× IQR and outliers shown in red. (II) Heatmap shows individual electrode amplitude values for each day of recording, white areas indicate field potential recorded. **(C)** Field potential duration, (I) Boxplot of field potential amplitude across 22 days of chronic recording, with whiskers extending to 1.5× IQR and outliers shown in red. (II) Heatmap shows individual electrode duration values for each day of recording, white areas indicate potassium peak found in recorded data. **(D)** Beat frequency over 22 days of chronic recording. **(E)** Conduction velocity over 19 days of chronic recording, as after 19 days conduction velocity could not be calculated.

**Figure S10:**
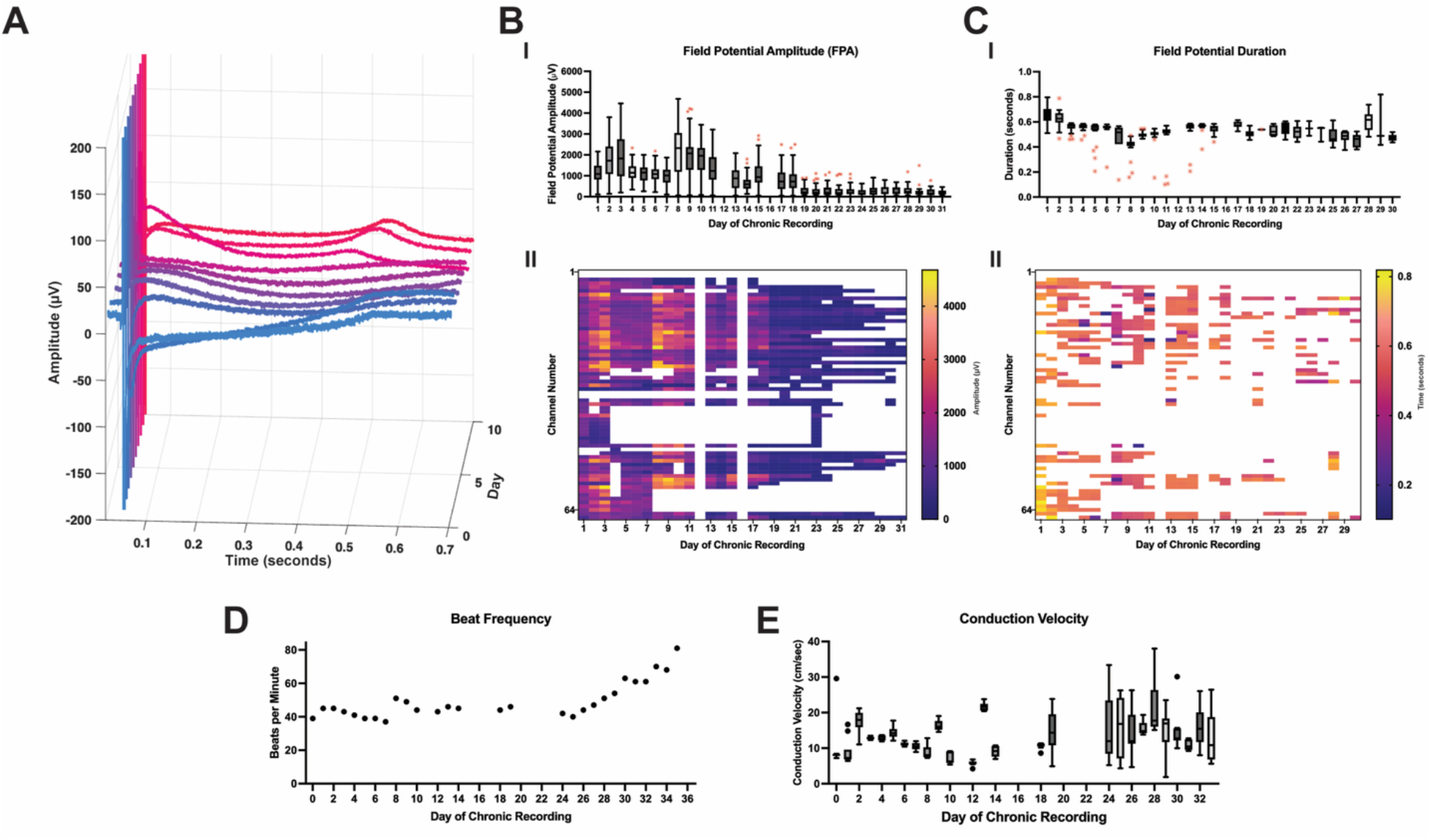
Additional Chronic Electrophysiology Recording from untreated iPSC-CM monolayer. **(A)** Representative extracellular field potential from a single channel over first 10 days of chronic recordings. **(B)** Field Potential Amplitude (I) Boxplot of field potential amplitude across 10 days of chronic recording, with whiskers extending to 1.5× IQR and outliers shown in red. (II) Heatmap shows individual electrode amplitude values for each day of recording, white areas indicate no field potential in recording data. **(C)** Field potential duration, (I) Boxplot of field potential amplitude across 10 days of chronic recording, with whiskers extending to 1.5× IQR and outliers shown in red. (II) Heatmap shows individual electrode duration values for each day of recording, white areas indicate no potassium peak in recorded data. **(D)** Beat frequency over 10 days of chronic recording. **(E)** Conduction velocity over 10 days of chronic recording.

**Figure S11:**
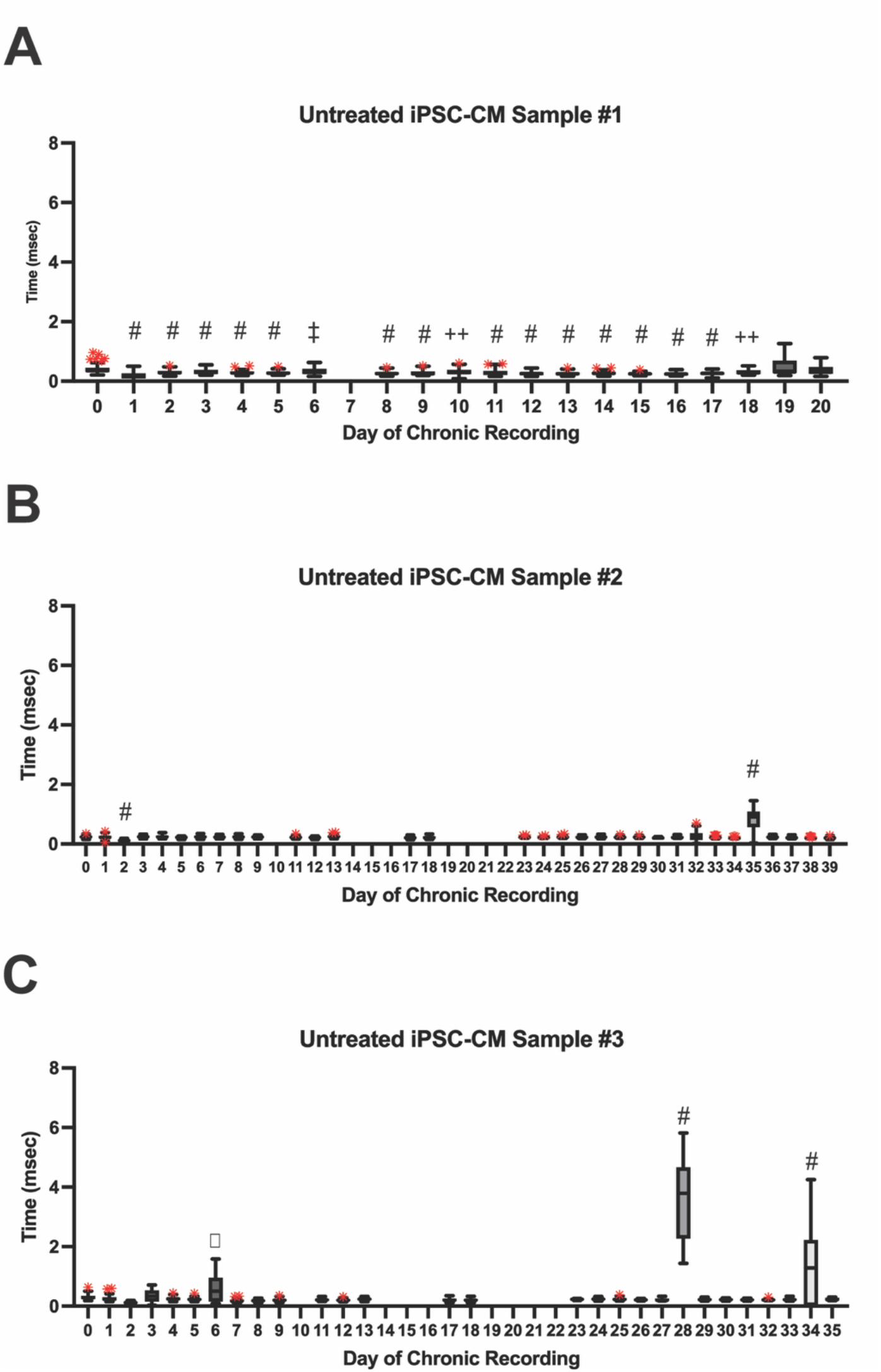
Sodium Width over time for untreated iPSC-CM. **(A-C)** Sodium width for each channel each day across the length of the chronic recording. Statistical significance was determined with a one-way ANOVA and Dunnett’s multiple comparisons test, ◻ indicates statistical significance with a p-value < 0.05, **‡** indicates statistical significance with a p-value < 0.01. **++** indicates statistical significance with a p-value < 0.001, **#** indicates statistical significance with a p-value < 0.0001.

**Figure S12:**
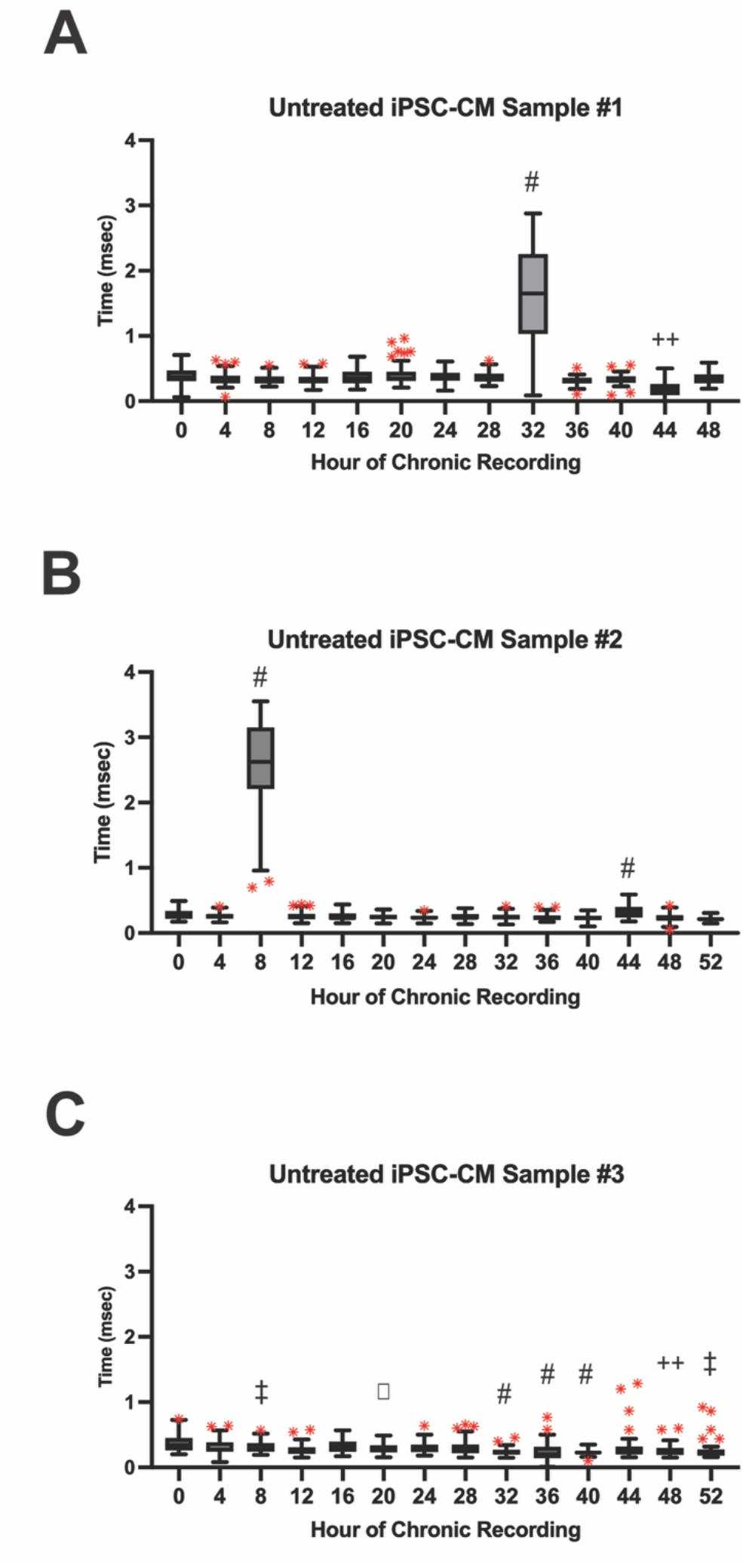
Sodium Width over time for untreated iPSC-CM. **(A-C)** Sodium width for each channel every four hours across the time frame of doxorubicin recordings. Statistical significance was determined with a one-way ANOVA and Dunnett’s multiple comparisons test, ◻ indicates statistical significance with a p-value < 0.05, **‡** indicates statistical significance with a p-value < 0.01. **++** indicates statistical significance with a p-value < 0.001, **#** indicates statistical significance with a p-value < 0.0001.

**Figure S13:**
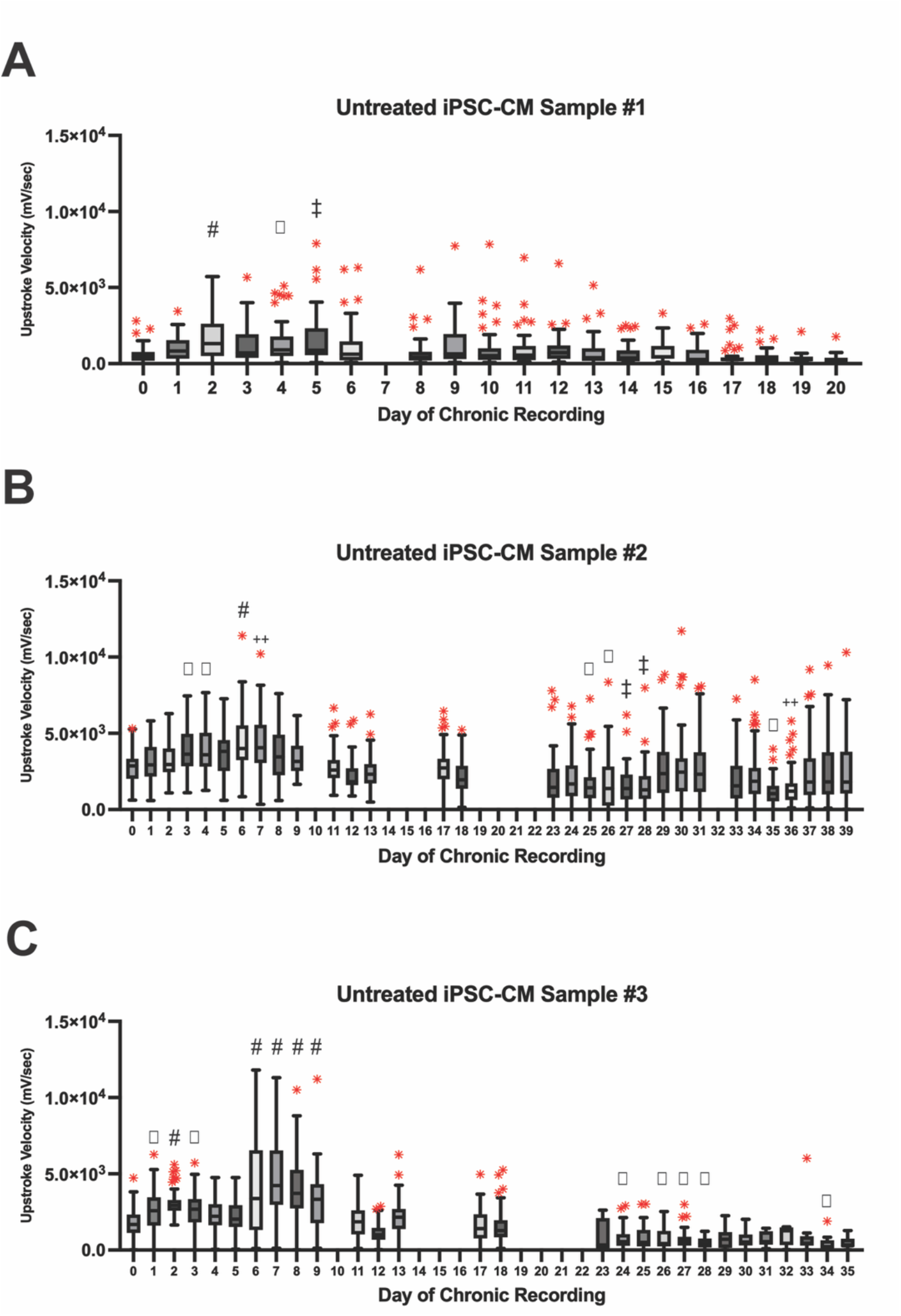
Upstroke Velocity / Max dV/dt over time for untreated iPSC-CM. **(A-C)** Upstroke velocity for each channel every day across the length of the recording. Statistical significance was determined with a one-way ANOVA and Dunnett’s multiple comparisons test, ◻ indicates statistical significance with a p-value < 0.05, **‡** indicates statistical significance with a p-value < 0.01. **++** indicates statistical significance with a p-value < 0.001, **#** indicates statistical significance with a p-value < 0.0001.

**Figure S14:**
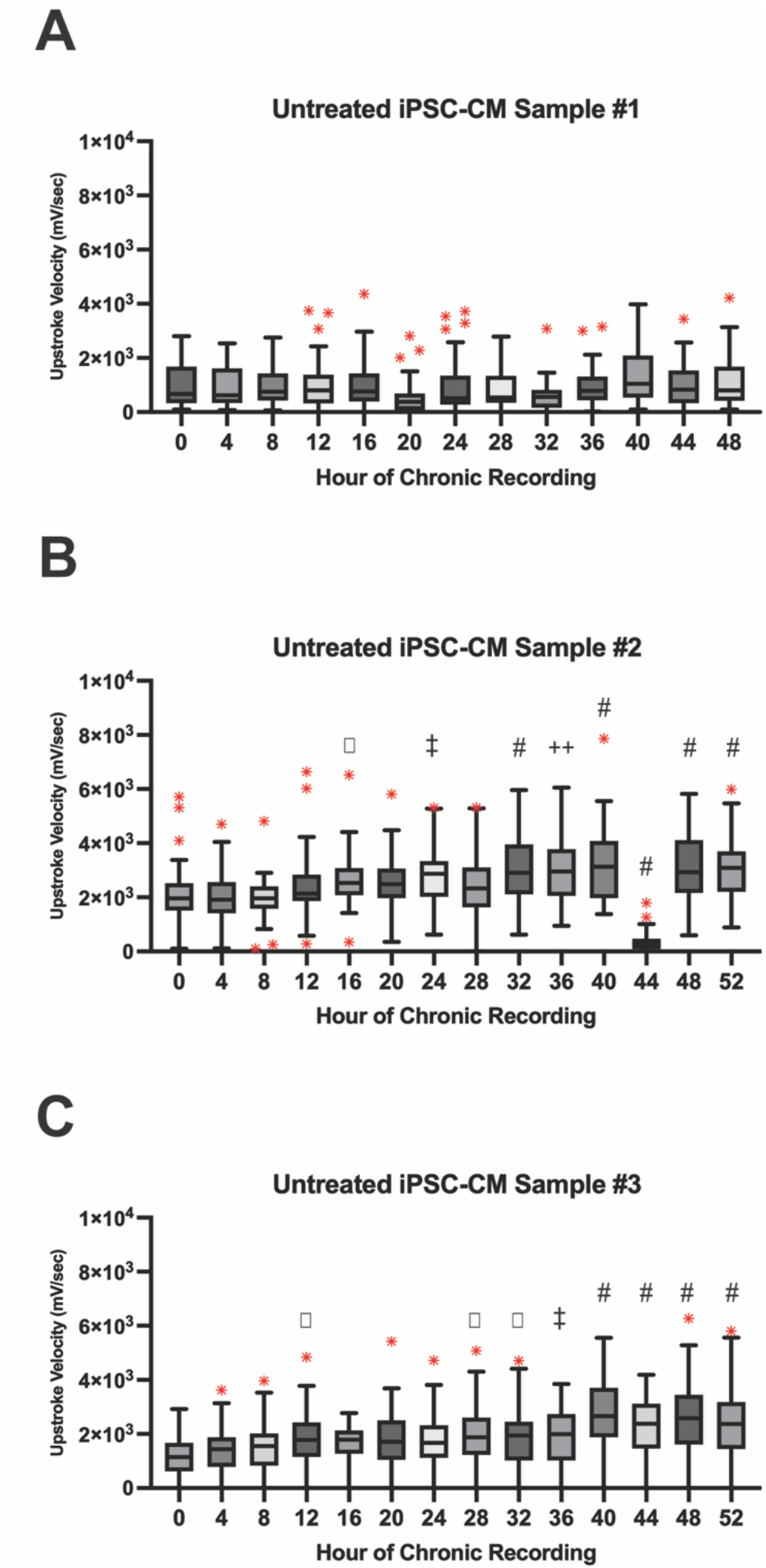
Upstroke Velocity / Max dV/dt over time for untreated iPSC-CM. **(A-C)** Upstroke velocity for each channel every four hours across the time frame of doxorubicin recordings. Statistical significance was determined with a one-way ANOVA and Dunnett’s multiple comparisons test, ◻ indicates statistical significance with a p-value < 0.05, **‡** indicates statistical significance with a p-value < 0.01. **++** indicates statistical significance with a p-value < 0.001, **#** indicates statistical significance with a p-value < 0.0001.

**Figure S15:**
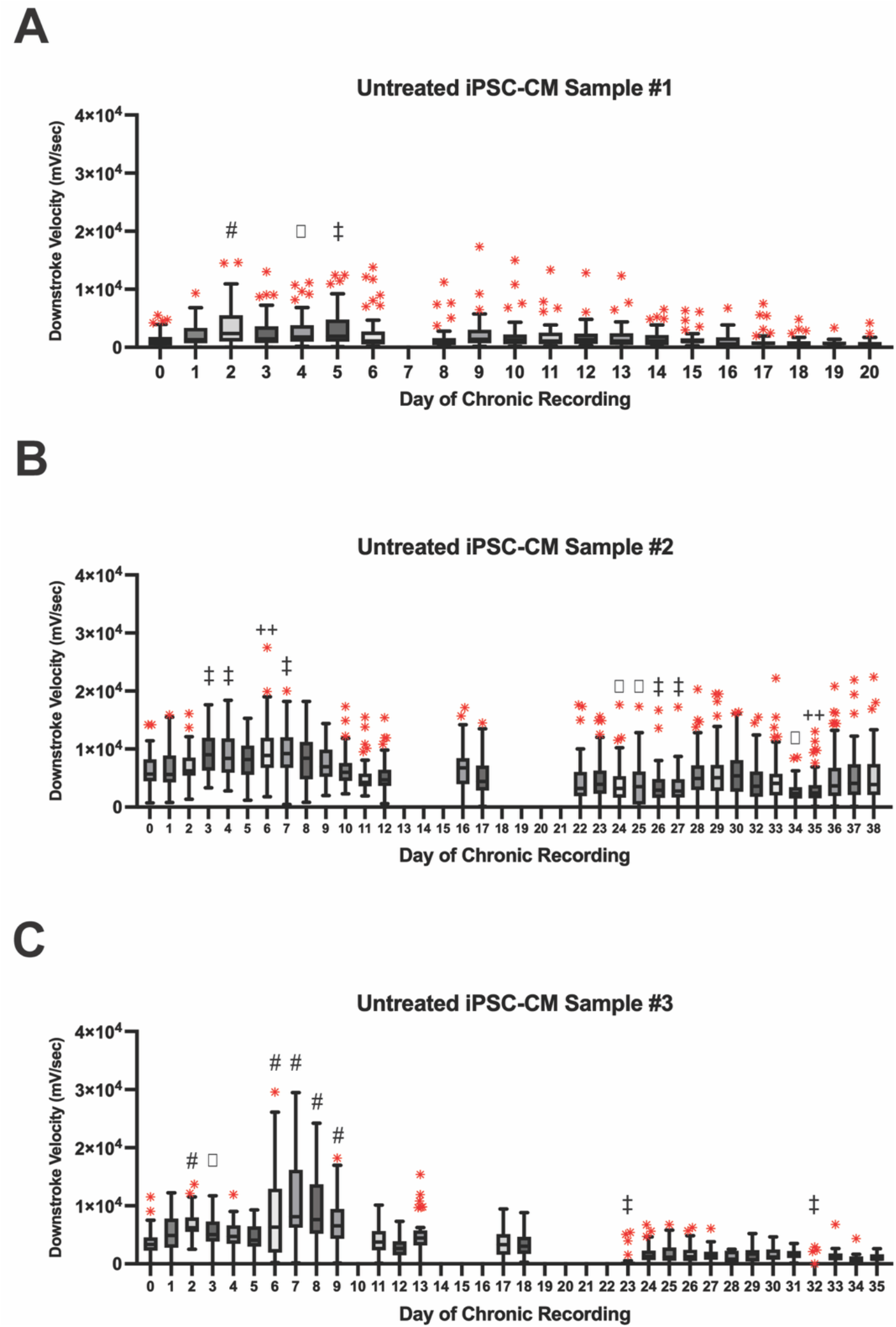
Downstroke Velocity / Min dV/dt over time for untreated iPSC-CM. **(A-C)** Downstroke velocity for each channel every day across the length of the recording. Statistical significance was determined with a one-way ANOVA and Dunnett’s multiple comparisons test, ◻ indicates statistical significance with a p-value < 0.05, **‡** indicates statistical significance with a p-value < 0.01. **++** indicates statistical significance with a p-value < 0.001, **#** indicates statistical significance with a p-value < 0.0001.

**Figure S16:**
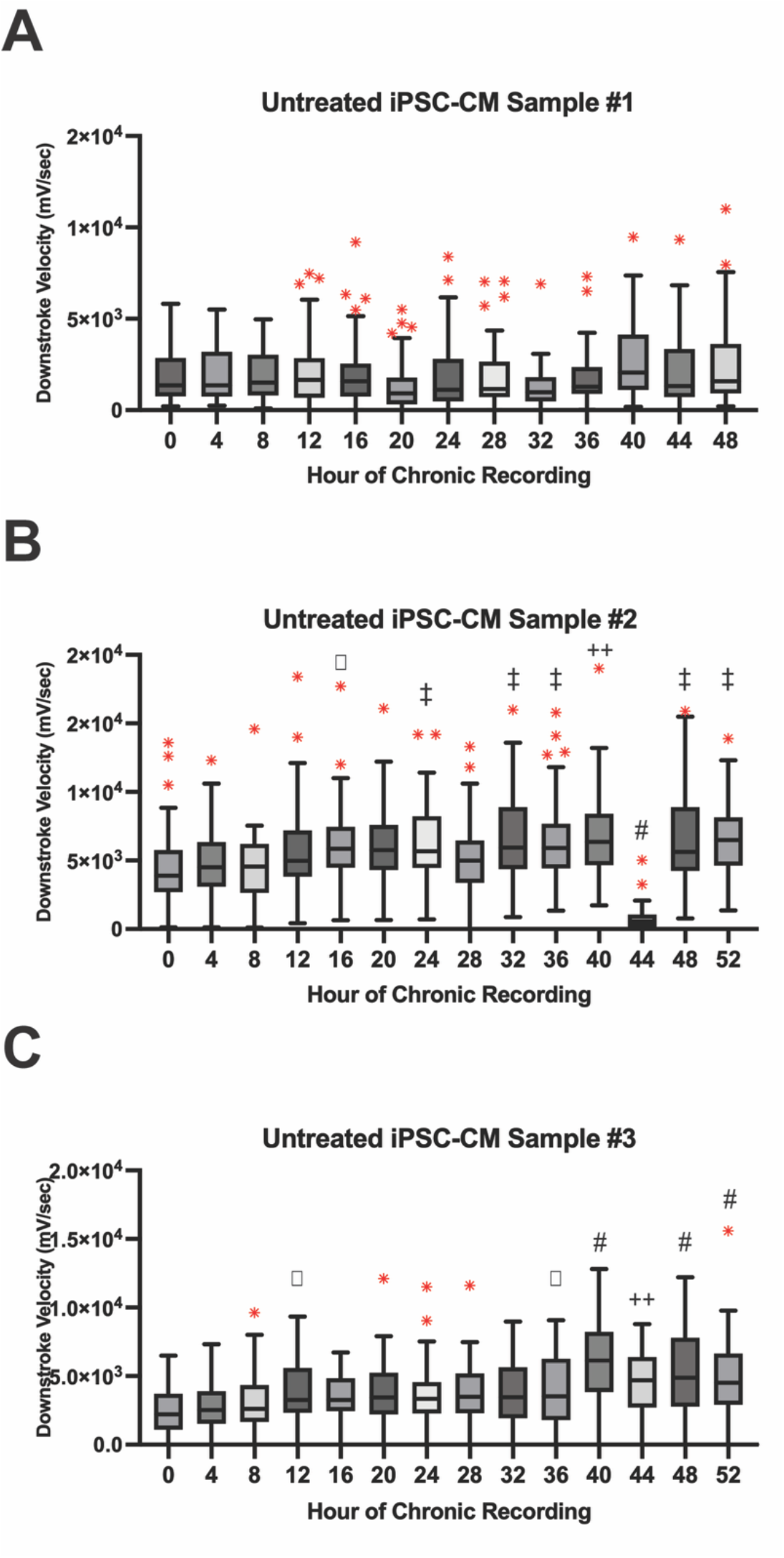
Downstroke Velocity / Min dV/dt over time for untreated iPSC-CM. **(A-C)** Downstroke velocity for each channel every four hours across the time frame of doxorubicin recordings. Statistical significance was determined with a one-way ANOVA and Dunnett’s multiple comparisons test, ◻ indicates statistical significance with a p-value < 0.05, **‡** indicates statistical significance with a p-value < 0.01. **++** indicates statistical significance with a p-value < 0.001, **#** indicates statistical significance with a p-value < 0.0001.

**Figure S17:**
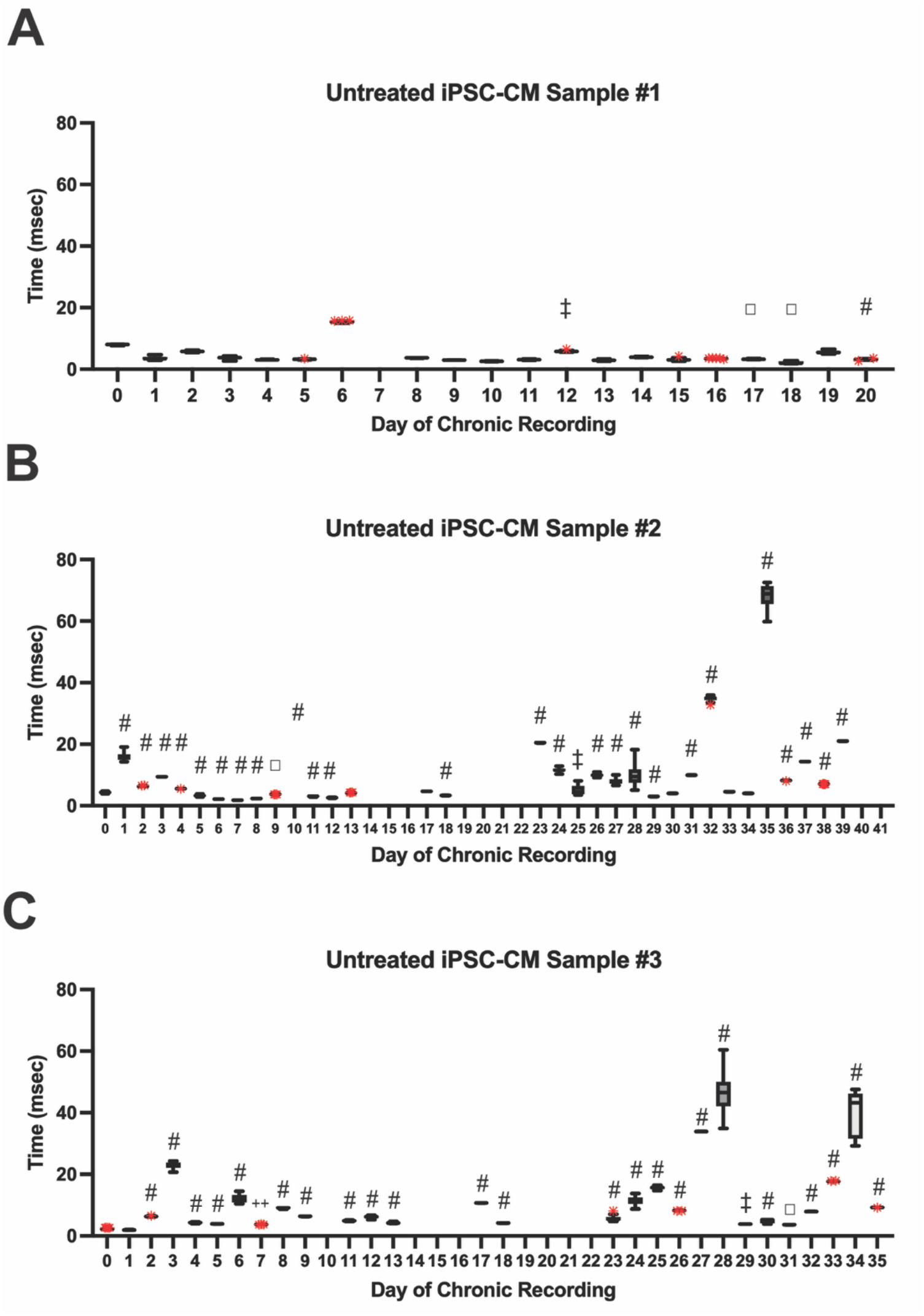
Beat Rate Variability over time for untreated iPSC-CM. **(A-C)** RMSSD calculated beat rate variability for each channel every day across the time frame of recording. Statistical significance was determined with a one-way ANOVA and Dunnett’s multiple comparisons test, ◻ indicates statistical significance with a p-value < 0.05, **‡** indicates statistical significance with a p-value < 0.01. **++** indicates statistical significance with a p-value < 0.001, **#** indicates statistical significance with a p-value < 0.0001.

**Figure S18:**
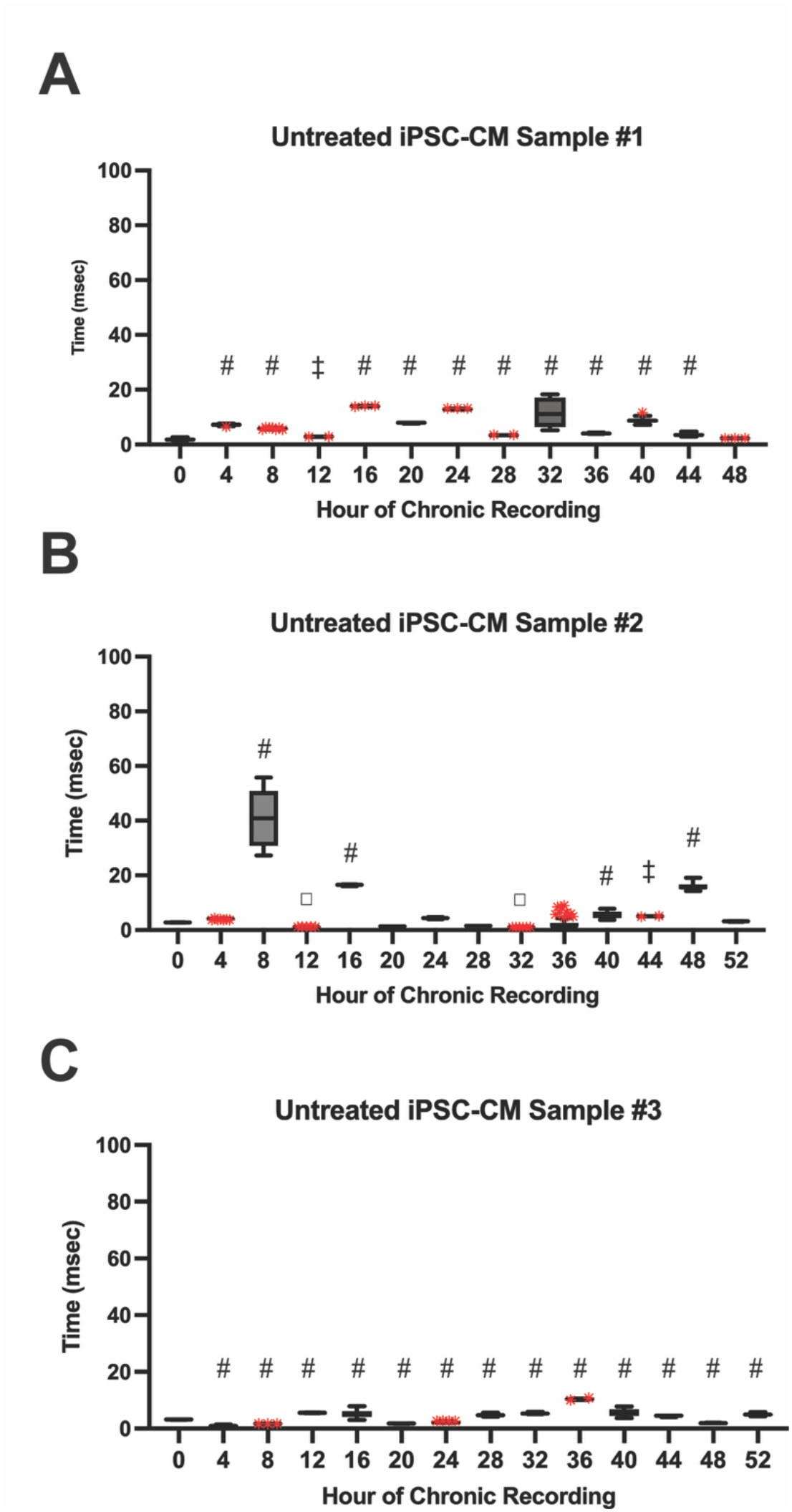
Beat Rate Variability over time for untreated iPSC-CM. **(A-C)** RMSSD calculated beat rate variability for each channel every four hours across the time frame of doxorubicin recordings. Statistical significance was determined with a one-way ANOVA and Dunnett’s multiple comparisons test, ◻ indicates statistical significance with a p-value < 0.05, **‡** indicates statistical significance with a p-value < 0.01. **++** indicates statistical significance with a p-value < 0.001, **#** indicates statistical significance with a p-value < 0.0001.

**Figure S19:**
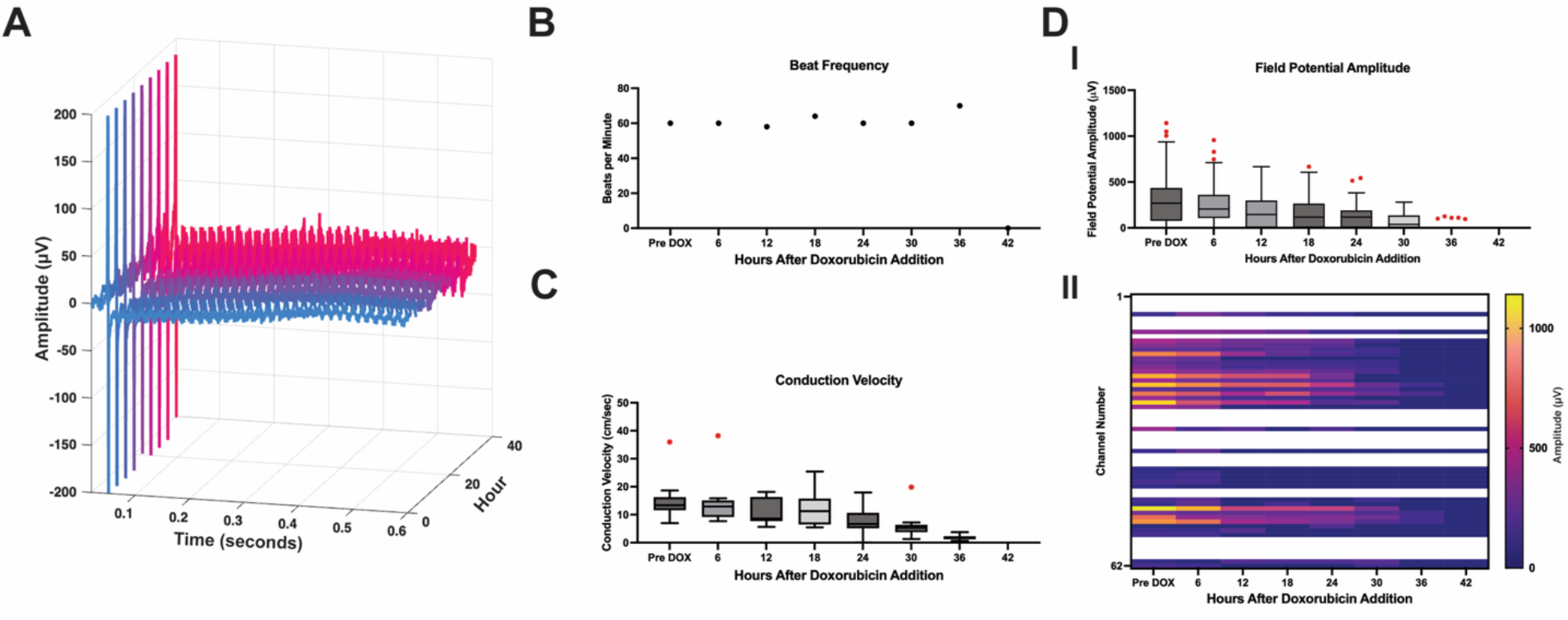
Additional 0.50 μM doxorubicin treated iPSC-CM chronic recording. (A) Representative extracellular field potential across all hours of recording. (B) Beat frequency across 36 hours of chronic recording. (C) Conduction velocity across the cardiac monolayer across 36 hours of chronic recording. (D) Field potential amplitude, (I) Boxplot of field potential amplitude across 42 hours of chronic recording, with whiskers extending to 1.5× IQR and outliers shown in red. (II) Heatmap shows individual electrode amplitude values for every 6 hours of recording, white areas indicate no field potentials in recording data.

**Figure S20:**
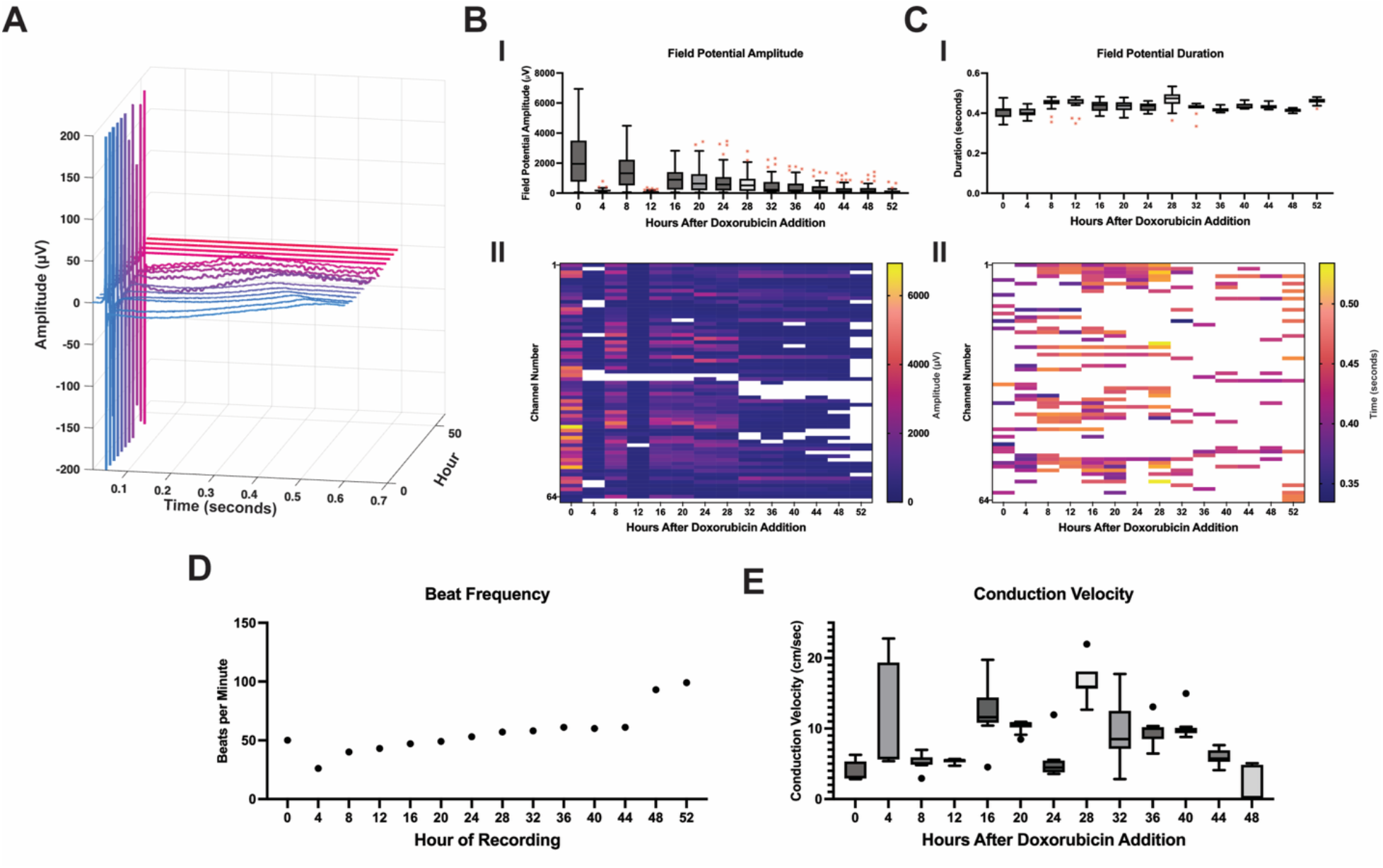
Additional 0.50 μM doxorubicin treated iPSC-CM chronic recording. **(A)** Representative extracellular field potential across all hours of recording. **(B)** Field potential amplitude, (I) Boxplot of field potential amplitude across 52 hours of chronic recording, with whiskers extending to 1.5× IQR and outliers shown in red. (II) Heatmap shows individual electrode amplitude values for every 4 hours of recording, white areas indicate no field potentials in recording data. **(C)** Field potential duration, (I) Boxplot of field potential duration across 52 hours of chronic recording, with whiskers extending to 1.5× IQR and outliers shown in red. (II) Heatmap shows individual electrode duration values for every 4 hours of recording, white areas indicate no potassium peak in recording data. (D) Beat frequency across 56 hours of chronic recording. (E) Conduction velocity across the cardiac monolayer across 48 hours of chronic recording.

**Figure S21:**
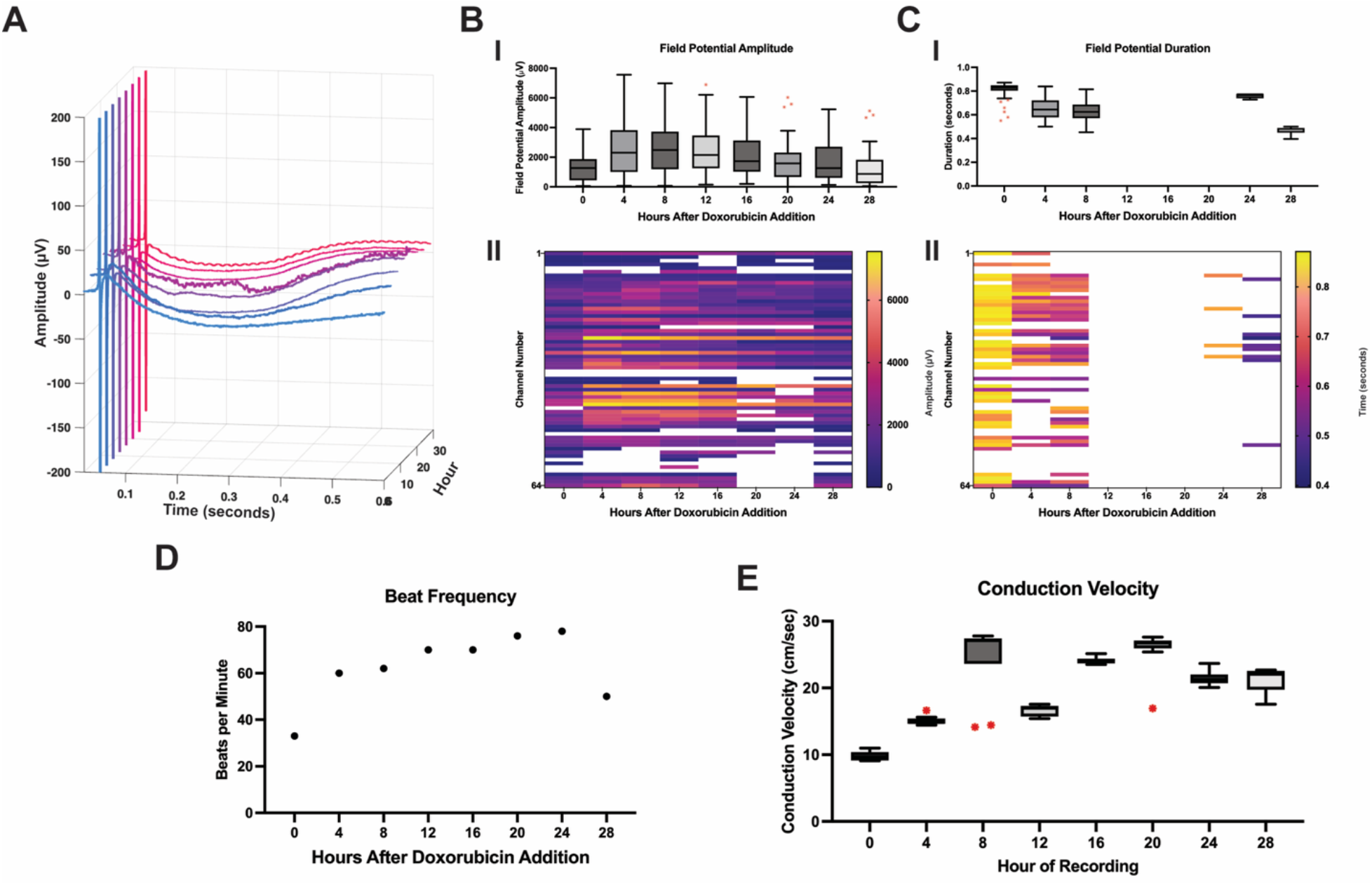
Additional 0.50 μM doxorubicin treated iPSC-CM chronic recording. **(A)** Representative extracellular field potential across all hours of recording. **(B)** Field potential amplitude, (I) Boxplot of field potential amplitude across 28 hours of chronic recording, with whiskers extending to 1.5× IQR and outliers shown in red. (II) Heatmap shows individual electrode amplitude values for every 4 hours of recording, white areas indicate no field potentials in recording data. **(C)** Field potential duration, (I) Boxplot of field potential duration across 28 hours of chronic recording, with whiskers extending to 1.5× IQR and outliers shown in red. (II) Heatmap shows individual electrode duration values for every 4 hours of recording, white areas indicate no potassium peak in recording data. (D) Beat frequency across 28 hours of chronic recording. (E) Conduction velocity across the cardiac monolayer across 28 hours of chronic recording.

**Figure S22:**
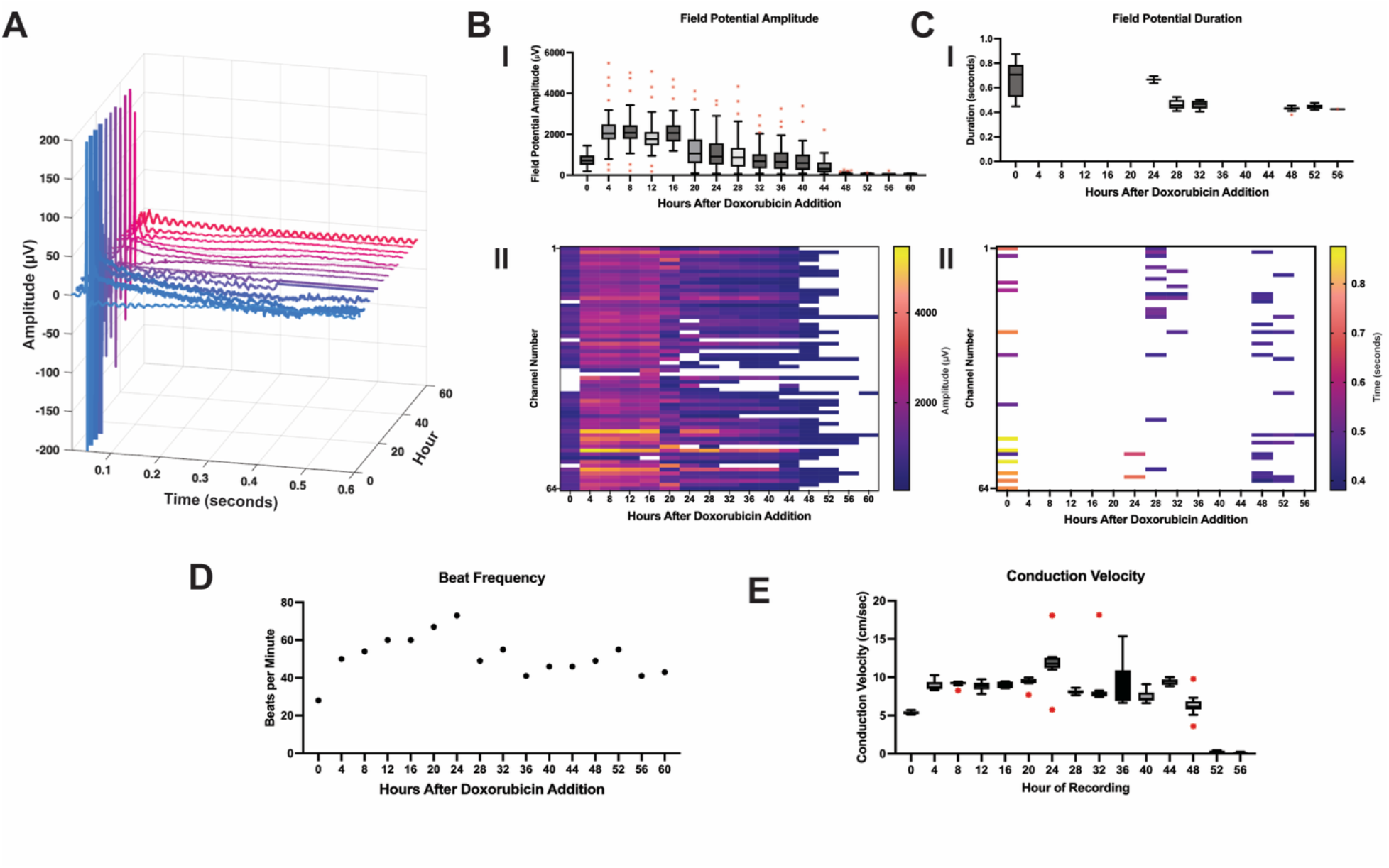
Additional 0.50 μM doxorubicin treated iPSC-CM chronic recording. **(A)** Representative extracellular field potential across all hours of recording. **(B)** Field potential amplitude, (I) Boxplot of field potential amplitude across 60 hours of chronic recording, with whiskers extending to 1.5× IQR and outliers shown in red. (II) Heatmap shows individual electrode amplitude values for every 4 hours of recording, white areas indicate no field potentials in recording data. **(C)** Field potential duration, (I) Boxplot of field potential duration across 60 hours of chronic recording, with whiskers extending to 1.5× IQR and outliers shown in red. (II) Heatmap shows individual electrode duration values for every 4 hours of recording, white areas indicate no potassium peak in recording data. (D) Beat frequency across 60 hours of chronic recording. (E) Conduction velocity across the cardiac monolayer across 56 hours of chronic recording.

**Figure S23:**
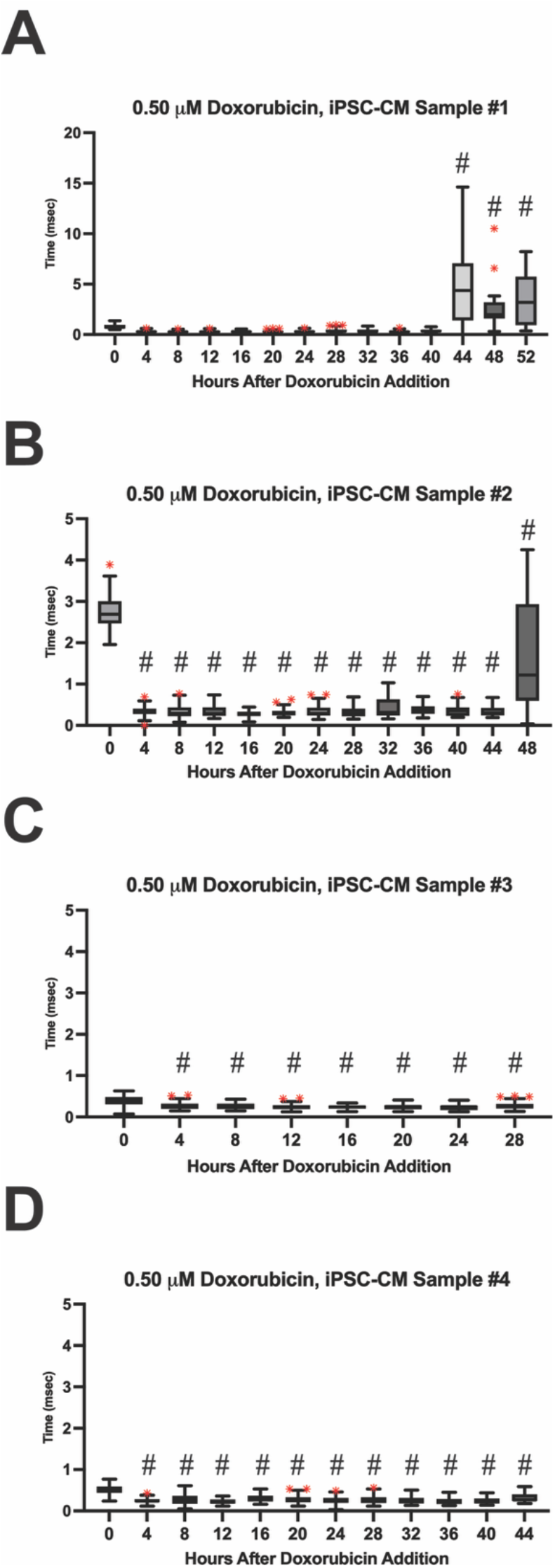
Sodium Width over time for iPSC-CM administered 0.50 µM doxorubicin. (A-D) Sodium width for each channel every four hours across the length of the chronic recording. Statistical significance was determined with a one-way ANOVA and Dunnett’s multiple comparisons test, ◻ indicates statistical significance with a p-value < 0.05, **‡** indicates statistical significance with a p-value < 0.01. **++** indicates statistical significance with a p-value < 0.001, **#** indicates statistical significance with a p-value < 0.0001.

**Figure S24:**
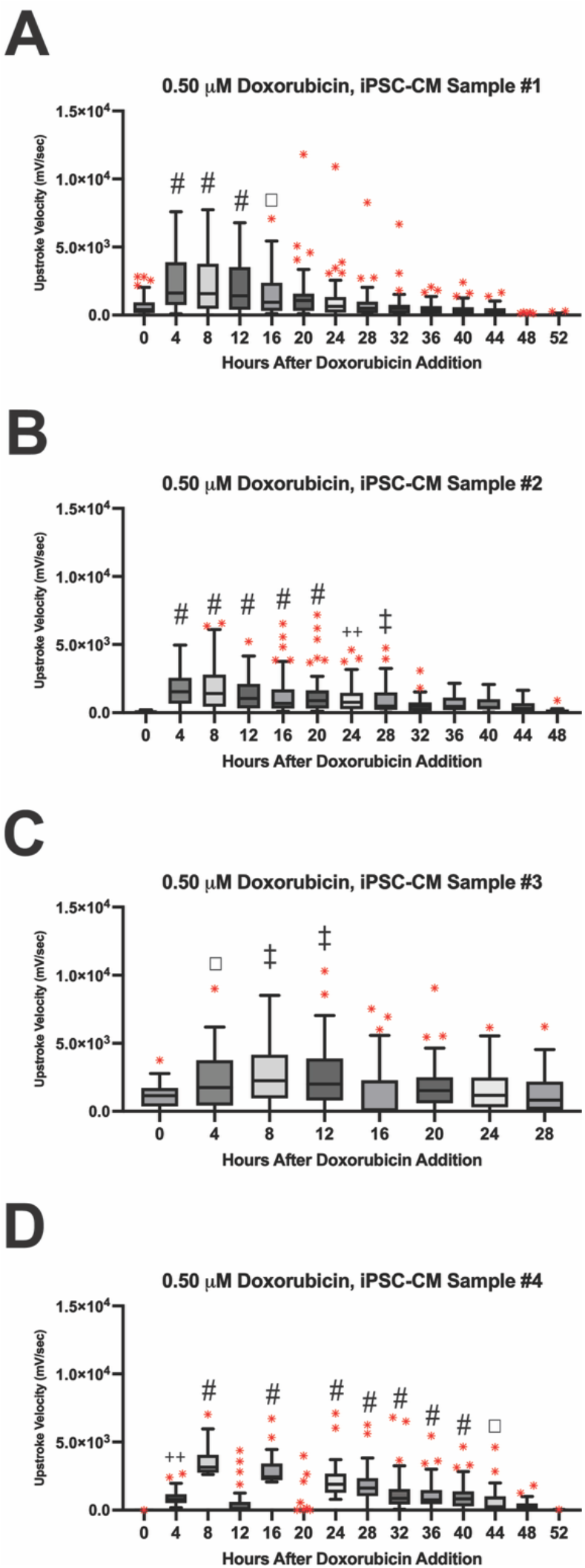
Upstroke Velocity / Max dV/dt over time for iPSC-CM administered 0.5µM doxorubicin. (A-D) Upstroke velocity for each channel every four hours across the time frame of doxorubicin recordings. Statistical significance was determined with a one-way ANOVA and Dunnett’s multiple comparisons test, ◻ indicates statistical significance with a p-value < 0.05, **‡** indicates statistical significance with a p-value < 0.01. **++** indicates statistical significance with a p-value < 0.001, **#** indicates statistical significance with a p-value < 0.0001.

**Figure S25:**
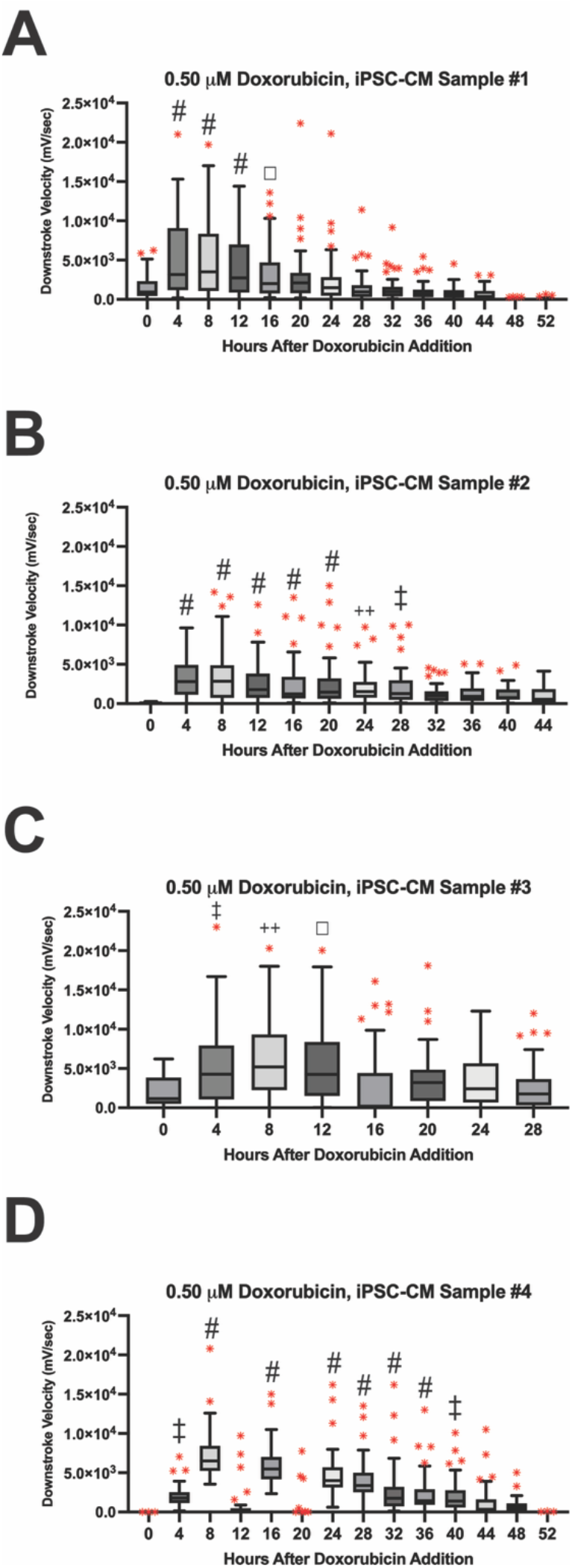
Downstroke Velocity / Min dV/dt over time for iPSC-CM administered 0.5µM doxorubicin. (A-D) Downstroke velocity for each channel every four hours across the time frame of doxorubicin recordings. Statistical significance was determined with a one-way ANOVA and Dunnett’s multiple comparisons test, ◻ indicates statistical significance with a p-value < 0.05, **‡** indicates statistical significance with a p-value < 0.01. **++** indicates statistical significance with a p-value < 0.001, **#** indicates statistical significance with a p-value < 0.0001.

**Figure S26:**
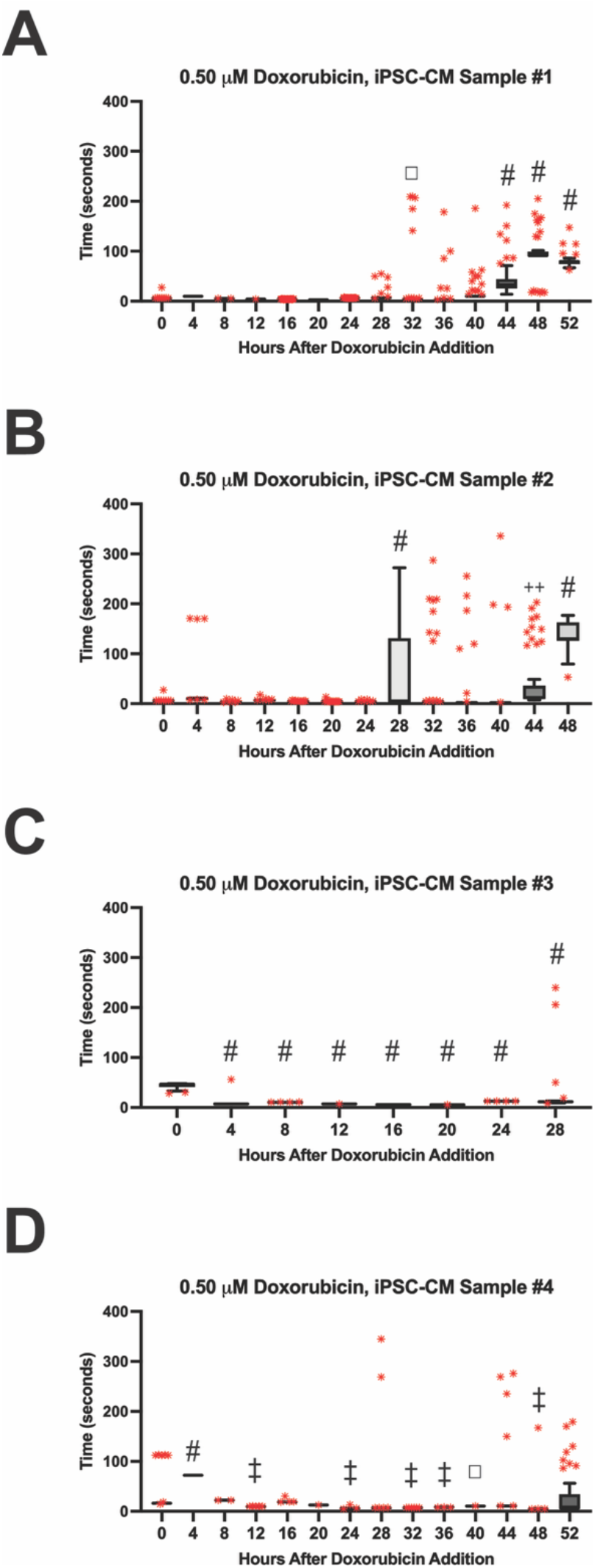
Beat Rate Variability over time for iPSC-CM administered 0.5µM doxorubicin. (A-D) RMSSD calculated beat rate variability for each channel every four hours across the time frame of doxorubicin recordings. Statistical significance was determined with a one-way ANOVA and Dunnett’s multiple comparisons test, ◻ indicates statistical significance with a p-value < 0.05, **‡** indicates statistical significance with a p-value < 0.01. **++** indicates statistical significance with a p-value < 0.001, **#** indicates statistical significance with a p-value < 0.0001.

**Figure S27:**
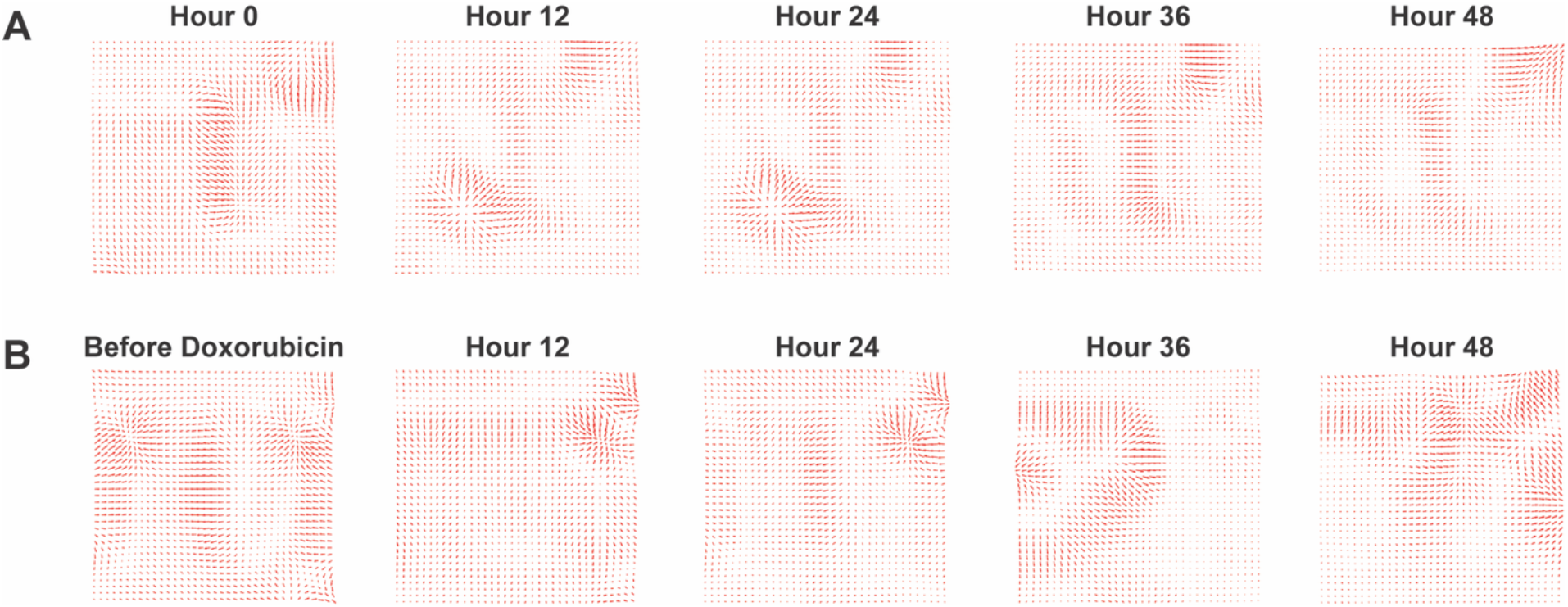
Quiver Plots from chronic electrophysiology recordings. **(A)** Quiver maps representing signal propagation waves across an untreated cardiac monolayer over time. **(B)** Quiver maps representing signal propagation waves across the cardiac monolayer over time after being administered 0.5 µM doxorubicin for 24 hours.

**Figure S28:**
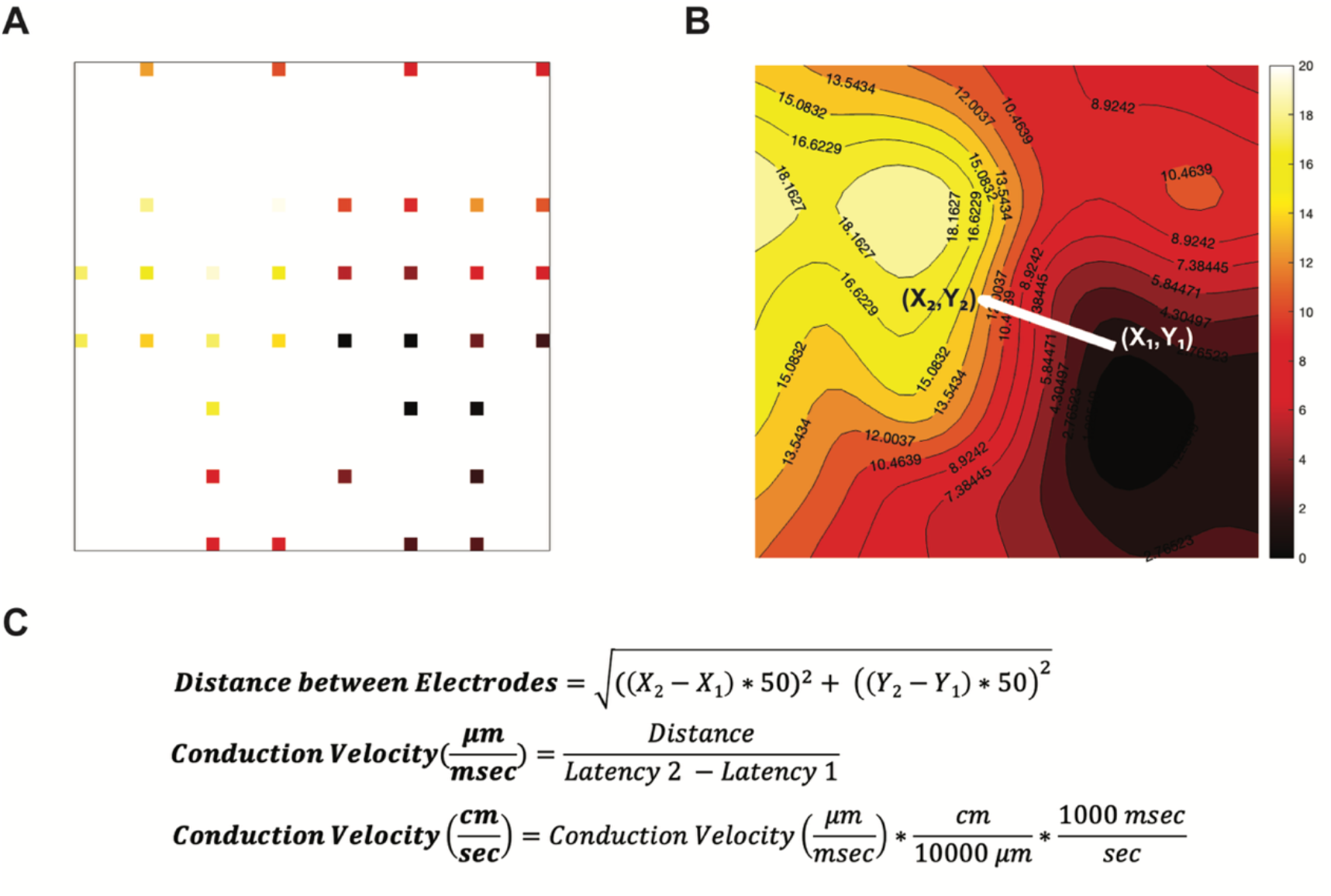
Conduction Velocity Calculations from 64 channel Microelectrode Array. **(A)** Time latency values plotted at the location of each functional electrode. White areas indicate areas without functional electrodes or areas between electrodes. **(B)** Interpolated time latency isochronal map. White arrow indicates average direction and magnitude of wave propagation across the map. **(C)** Formulas for calculating conduction velocity from the isochronal map created from time latency values.

**Figure S29:**
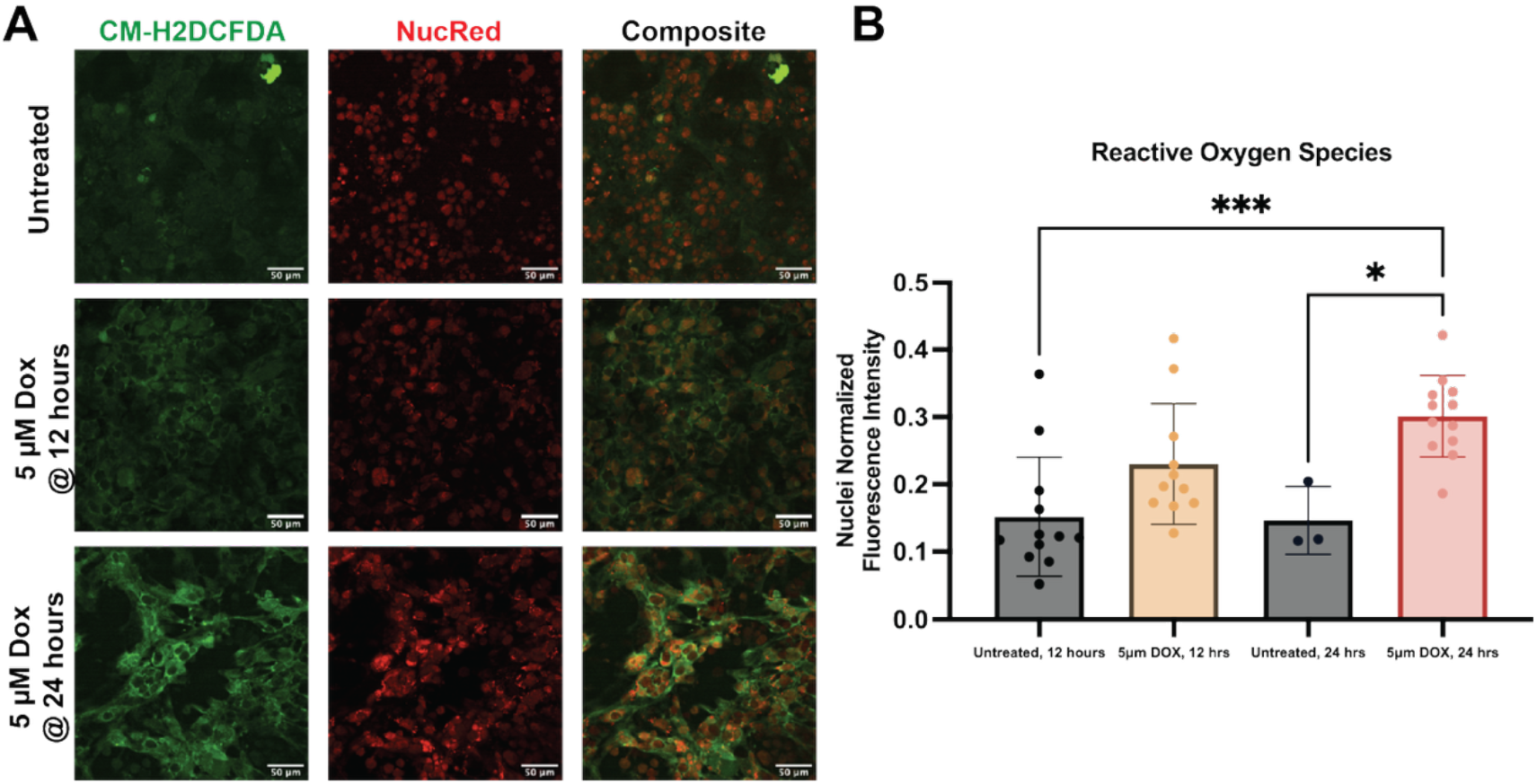
Reactive Oxygen Species Increase when Exposed to 5 µM doxorubicin. **(A)** Representative images of reactive oxygen species using CM-H2DCFA in untreated and 5.0 µM doxorubicin iPSC-CMs after 12 and 24 hours of exposure. **(B)** Quantification of the fluorescence intensity of reactive oxygen species normalized per image by the nuclei count. 3 samples were imaged per condition, with 3-4 fields of view analyzed per sample. Statistical significance was determined with a one-way ANOVA with Tukey’s multiple comparison, * indicates statistical significance with a p-value < 0.05, *** indicates statistical significance with a p-value < 0.001.

**Figure S30:**
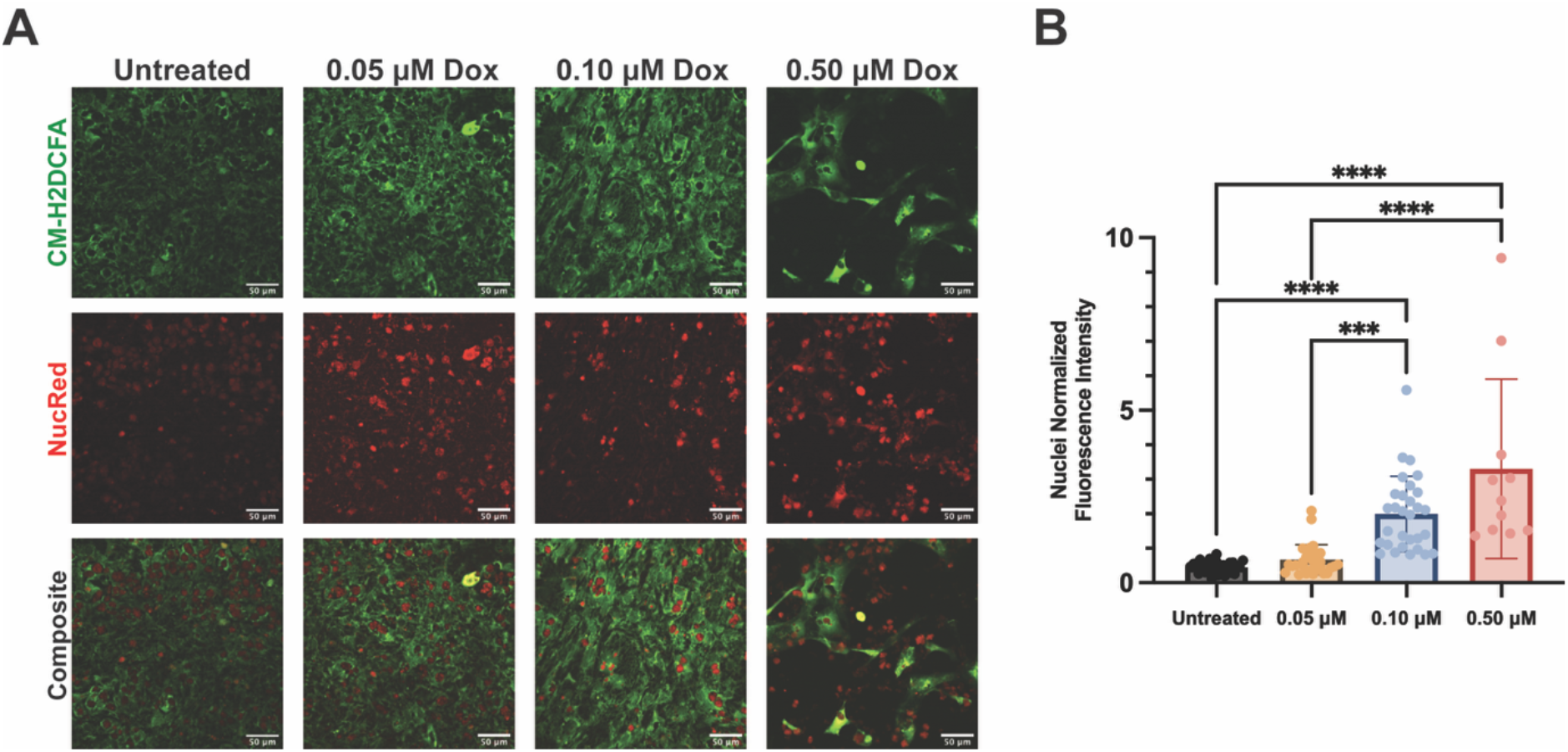
Doxorubicin induced Reactive Oxygen Species. **(A)** Representative images of reactive oxygen species using CM-H2DCFA dye to react with ROS within the cell and NucRed to stain nuclei in untreated and doxorubicin treated (0.05 µM, 0.10 µM, 0.50 µM) iPSC-CMs. **(B)** Quantification of the fluorescence intensity of reactive oxygen species normalized per image by the nuclei count. 5 samples were imaged per condition, with 6 fields of view analyzed per sample. Statistical significance was determined with a one-way ANOVA with Tukey’s multiple comparison, *** indicates statistical significance with a p-value < 0.001, **** indicates statistical significance with a p-value < 0.0001.

**Figure S31:**
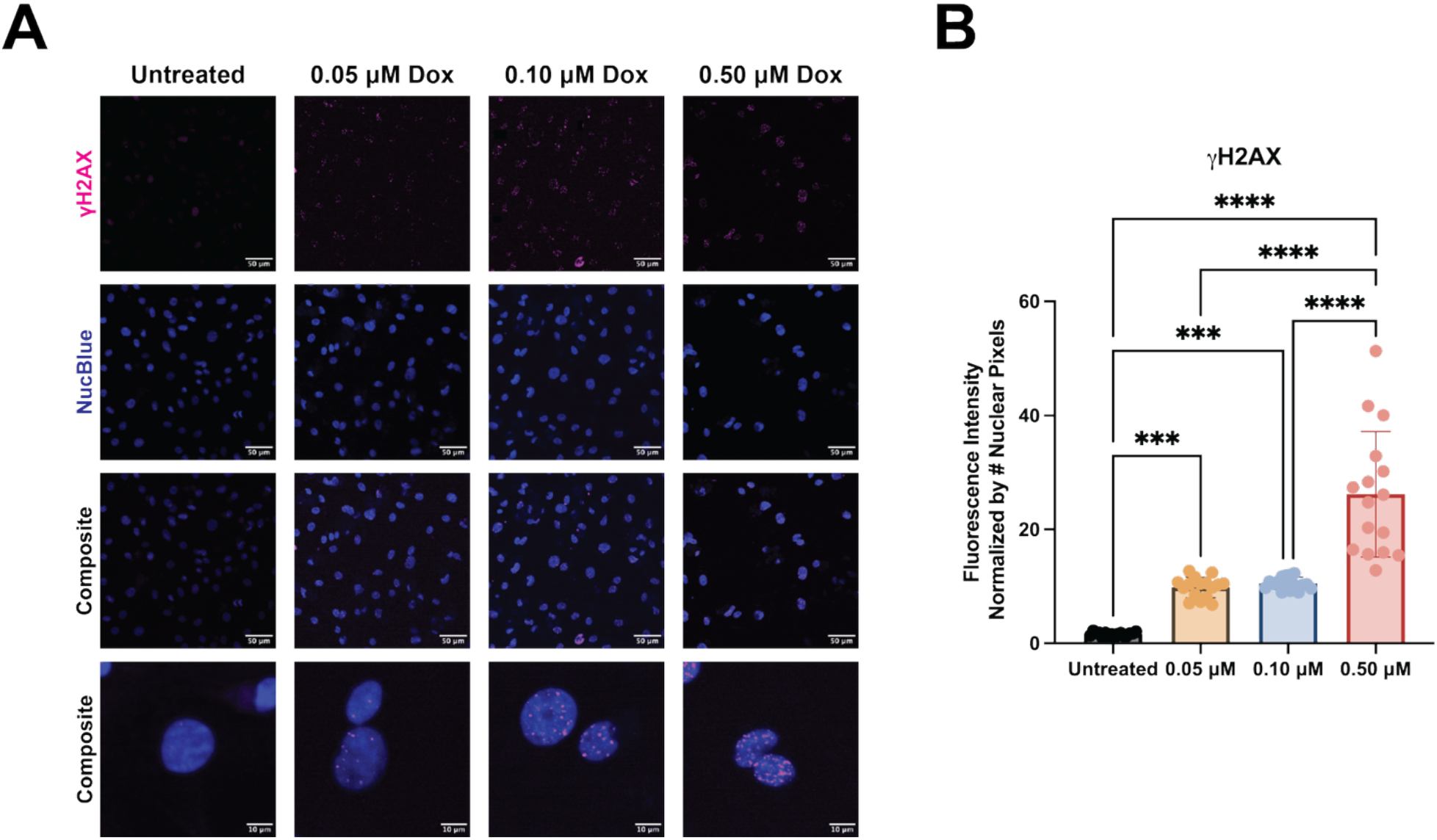
Doxorubicin Causes Increase in DNA Damage. **(A)** Representative images of immunofluorescence detection of *γ*H2AX puncta in the nuclei of untreated and doxorubicin treated (0.05 µM, 0.10 µM, 0.50 µM) iPSC-CMs. **(B)** Quantification of the *γ*H2AX puncta in each of the imaging condition normalized by the number of nuclei pixels in each image. 4 samples were imaged per condition, with 4 fields of view analyzed per sample. Statistical significance was determined with a one-way ANOVA with Tukey’s multiple comparison, ns indicates non-significance, *** indicates statistical significance with a p-value < 0.001, **** indicates statistical significance with a p-value < 0.0001.

